# Origin of the GPR15LG–GPR15 signaling axis in ancient fish ancestors

**DOI:** 10.64898/2026.06.02.729464

**Authors:** Jun-Jie Yao, Jie Yu, Hao-Zheng Li, Juan-Juan Wang, Ya-Li Liu, Zhan-Yun Guo

**Affiliations:** Research Center for Translational Medicine at East Hospital, School of Life Sciences and Technology, Tongji University, Shanghai, China; National Institute of Metrology of China, Beijing, China

**Keywords:** Activation, Binding, Fish, GPR15, GPR15LG

## Abstract

The chemokine-like peptide GPR15LG is a known agonist of G protein-coupled receptor 15 (GPR15), a ligand–receptor pair primarily implicated in mammalian mucosal immunity and lymphocyte homing. However, the evolutionary origin and phylogenetic distribution of this signaling system remain poorly understood due to the extreme sequence diversity of GPR15LG orthologs. In this study, we identified GPR15LG orthologs in several fish species for the first time according to their conserved gene synteny, genomic organization, and amino acid sequence features. A representative ortholog from the spotted gar (*Lepisosteus oculatus*), termed Lo-GPR15LG, was recombinantly prepared and functionally characterized using NanoLuc Binary Technology (NanoBiT)-based β-arrestin recruitment assay and homogenous ligand–receptor binding assay. Our results demonstrated that Lo-GPR15LG directly binds to and efficiently activates its cognate receptor, Lo-GPR15, with a dissociation constant (K_d_) of approximately 60 nM and an EC_50_ value of approximately 10 nM. Functional assays further revealed that receptor activation is critically dependent on the conserved C-terminal residues. Notably, human and fish orthologs exhibited no cross-species activity, consistent with their high sequence divergence. These findings reveal that the GPR15LG–GPR15 signaling system originated in ancient fish ancestors and has remained a conserved signaling axis throughout vertebrate evolution, suggesting a fundamental role in immunity across all vertebrate lineages.

## 1. Introduction

G protein-coupled receptor 15 (GPR15) is predominantly expressed in specific immune cell populations, including T cells, B cells, and plasma cells, according to data from the Human Protein Atlas (https://www.proteinatlas.org). Early studies identified GPR15 as a coreceptor for human immunodeficiency virus (HIV) and simian immunodeficiency virus (SIV) [1–8]. In 2013, GPR15 was shown to mediate the homing of specific T cell subsets to the large intestinal mucosa, thereby contributing to immune homeostasis [9]. Subsequent studies have consistently supported its role in regulating lymphocyte trafficking under diverse physiological and pathological conditions [10–24].

In 2017, a chemokine-like secreted peptide, chromosome 10 open reading frame 99 (C10orf99), also known as colon-derived SUSD2 binding factor (CSBF) or antimicrobial peptide 57 (AP-57) [25,26], was identified as an agonist of GPR15 [27,28]. This molecule was subsequently termed GPR15 ligand (GPR15LG or GPR15L). According to the Human Protein Atlas, GPR15LG is primarily expressed in mucosal tissues, including the esophagus, colon, rectum, appendix, and vagina (https://www.proteinatlas.org). Functionally, GPR15LG acts as a chemoattractant that recruits GPR15-expressing lymphocytes and plays a critical role in mucosal immunity [29–32]. Despite accumulating evidence supporting this ligand–receptor interaction, the GPR15LG–GPR15 pairing has not yet been formally recognized, and GPR15 remains classified as an orphan receptor in the International Union of Pharmacology database (https://www.guidetopharmacology.org).

According to the National Center for Biotechnology Information (NCBI) gene database, GPR15 orthologs are widely distributed in mammals, birds, and reptiles, but only identified in a limited number of amphibian and fish species. Instead, a less homologous “G protein-coupled receptor 15-like” receptor is broadly found across fish species. On the other hand, GPR15LG orthologs are well documented in mammals but have only been identified in a few birds and reptiles, and have not yet been reported in fish species within public databases. Consequently, the evolutionary origin and phylogenetic distribution of the GPR15LG–GPR15 signaling system remain poorly understood.

In this study, we identified putative GPR15LG orthologs in several fish species based on their conserved gene synteny, genomic organization, and amino acid sequence features. Due to lacking significant overall sequence similarity to known vertebrate counterparts, these fish orthologs were previously unrecognized, misannotated, or unannotated. Following recombinant expression, a representative ortholog from spotted gar (*Lepisosteus oculatus*), termed Lo-GPR15LG, demonstrated robust activity toward its corresponding Lo-GPR15 receptor in NanoLuc Binary Technology (NanoBiT)-based binding and activation assays, supporting the existence of a functional GPR15LG–GPR15 signaling pair in this primitive non-teleost ray-finned fish species [33]. These findings indicate that GPR15LG is an evolutionarily conserved ligand of GPR15 since originating in ancient fish ancestors.

## 2. Materials and methods

### Preparation of mature Lo-GPR15LG via bacterial overexpression

A DNA fragment encoding mature peptide of Lo-GPR15LG was chemically synthesized using *Escherichia coli*-biased codons and ligated into a small ubiquitin-related modifier (SUMO)-fusion vector via Gibson assembly, resulting in expression construct pET/6×His-SUMO-Lo-GPR15LG (Fig. S1). The expression construct for the C-terminally truncated version was generated via QuikChange mutagenesis, resulting in the expression construct pET/6×His-SUMO-Lo-[desC4]GPR15LG. The expression construct for the small fragment for NanoBiT (SmBiT)-fused version was generated by amplification of the coding region via polymerase chain reaction (PCR) using pET/6×His-SUMO-Lo-GPR15LG as template and subsequent ligation into the SUMO-fusion vector via Gibson assembly, resulting in the expression construct pET/6×His-SUMO-SmBiT-Lo-GPR15LG (Fig. S1). The coding region of Lo-GPR15LG in these expression constructs was confirmed by DNA sequencing.

Thereafter, the SUMO-fused precursors were overexpressed in the *E. coli* strain BL21(DE3) via induction by 1.0 mM isopropyl β-D-1-thiogalactopyranoside (IPTG) at 37ºC for 6–8 h. After harvest via centrifugation (5000 *g*, 10 min), the bacteria pellet was suspended in denaturation buffer (20 mM phosphate, 0.5 M NaCl, 6 M guanidine chloride, pH 7.4) and lysed by sonification. After centrifugation (10000 *g*, 30 min), the lysate supernatant was subjected to *S*-sulfonation by addition of sodium sulfite (Na_2_SO_3_) and sodium tetrathionate (Na_2_S_4_O_6_) to the final concentrations of 100 mM and 80 mM, respectively. After shaking at room temperature for 2–3 h, the mixture was subjected to immobilized metal ion affinity chromatography (Ni^2+^ column). After being washed by 10 mM imidazole (in denaturation buffer), the Ni^2+^ column was thoroughly rinsed by renaturation buffer (20 mM phosphate, 0.5 M NaCl, pH 7.4). Thereafter, the renatured SUMO-fused precursor was eluted from the Ni^2+^ column by 250 mM imidazole (in renaturation buffer) and treated with recombinant SUMO protease 6×His-R3-ULP1 at the enzyme-substrate molar ratio of 1:500. After overnight incubation at room temperature, the digestion mixture was 10-fold diluted with 20 mM phosphate buffer (pH7.4) and loaded onto a Ni^2+^ column again. After loading, the column was first eluted with 250 mM imidazole in 20 mM phosphate buffer and then eluted with 250 mM imidazole in denaturation buffer (20 mM phosphate, 0.5 M NaCl, 6 M guanidine chloride, pH 7.4).

The Lo-GRP15LG fraction eluted from the Ni^2+^ column by 250 mM imidazole in denaturation buffer was first treated with dithiothreitol (DTT) at the final concentration of 25 mM at 37ºC for ∼1 h, and then 25-fold diluted into ice-cold refolding buffer (0.5 M arginine, pH8.0, plus 2.0 mM oxidized glutathione). After refolding on ice for 7–8 h, the mixture was acidified to pH3–4 and applied to high performance liquid chromatography (HPLC). The refolded Lo-GPR15LG was eluted from a C_18_ reverse-phase column by an acidic acetonitrile gradient, manually collected, and lyophilized. After lyophilization, the purified Lo-GPR15LG proteins were dissolved in 1.0 mM aqueous hydrochloride (pH 3.0) and quantified by ultra-violet absorbance at 280 nm according to their theoretical extinction coefficient (1490 M^-1^ cm^-1^ for wild-type or C-terminally truncated Lo-GPR15LG; 2980 M^-1^ cm^-1^ for SmBiT-Lo-GPR15LG). During preparation, samples at different stages were analyzed by sodium dodecyl sulfate-polyacrylamide gel electrophoresis (SDS-PAGE).

### Recombinant expression of SUMO protease

An engineered enhanced version of SUMO protease, R3-ULP1, was prepared via bacteria overexpression according to previously reported procedures [34]. The DNA fragment encoding R3-ULP1 was chemically synthesized at Tsingke Biotechnology (Beijing, China) and ligated into pET28a vector between the cleavage sites of BamHI and NotI, resulting in the expression construct pET/6×His-R3-ULP1 (Fig. S2). Overexpression and downstream purification of the recombinant 6×His-R3-ULP1 was conducted according to the previously reported procedure [34].

### Generation of expression constructs for Lo-GPR15

The reference nucleotide sequence of the spotted gar (*L. oculatus*) GPR15, designated as Lo-GPR15, was retrieved from NCBI gene database (XM_006643117). Thereafter, a codon-optimized DNA fragment encoding Lo-GPR15 was chemically synthesized at CWbio (Taizhou, Jiangsu province, China) and ligated into pcDNA3.1(+) vector between the cleavage sites of NheI and NotI (Fig. S3). Thereafter, the Lo-GPR15 coding region was PCR amplified using appropriate primers and ligated into appropriate vectors for functional assays (Table S1). The doxycycline (Dox)-response expression construct pTRE3G-BI/Lo-GPR15-LgBiT:SmBiT-ARRB2 coexpresses a C-terminally large NanoLuc fragment for NanoBiT (LgBiT)-fused Lo-GPR15 (Lo-GPR15-LgBiT) and an N-terminally SmBiT-fused human β-arrestin 2 (SmBiT-ARRB2) for NanoBiT-based β-arrestin recruitment assays (Fig. S3). The Dox-response expression construct PB-TRE/sLgBiT-Lo-GPR15 expresses an N-terminally secretory LgBiT (sLgBiT)-fused Lo-GPR15 for NanoBiT-based ligand–receptor binding assays (Fig. S3). The coding region of Lo-GPR15 in these expression constructs was confirmed by DNA sequencing.

### NanoBiT-based β-arrestin recruitment assays

The NanoBiT-based β-arrestin recruitment assays were conducted on transiently transfected human embryonic kidney (HEK) 293T cells according to our previous procedure developed for other receptors [35–40]. Briefly, HEK293T cells were transiently cotransfected with the β-arrestin recruitment assay coexpression construct and the expression control plasmid pCMV-Tet3G (Clontech, Mountain View, CA, USA) using Lipo8000 transfection reagent (Beyotime Technology, Shanghai, China). Next day, the transfected cells were trypsinized, suspended in induction medium (complete DMEM plus 1.0 ng/mL of Dox), seeded into white opaque 96-well plates, and cultured at 37 °C for ∼24 h to ∼90% confluence. Thereafter, activation assays were conducted in following steps: removal of the induction medium, addition of pre-warmed activation solution (serum-free DMEM plus 1% bovine serum albumin) containing NanoLuc substrate (40 μL/well, containing 0.5 μL of NanoLuc substrate stock from Promega, Madison, WI, USA), measurement of bioluminescence on a SpectraMax iD3 plate reader (Molecular Devices, Sunnyvale, CA, USA) for ∼4 min, further addition of indicated peptide (10 μL/well, diluted in activation solution), and continuous measurement of bioluminescence for ∼10 min. If necessary, the measured absolute bioluminescence signals were corrected for inter well variability by forcing all curves after addition of NanoLuc substrate to same level. To generate the dose-response curve, the measured highest bioluminescence values were plotted with peptide concentrations using SigmaPlot 10.0 software (SYSTAT software, Chicago, IL, USA).

### NanoBiT-based ligand–receptor binding assays

The NanoBiT-based binding assays were conducted on transiently transfected HEK293T cells according to our previous procedure developed for other receptors [37–40]. Briefly, HEK293T cells were transiently transfected with the expression construct PB-TRE/sLgBiT-Lo-GPR15 with or without cotransfection of the tyrosylprotein sulfotransferase expression construct pTRE3G-BI/TPST1:TPST2 [40]. Next day, the transfected cells were trypsinized, suspended in induction medium (complete DMEM plus 20 ng/mL of Dox), seeded into white opaque 96-well plates, and cultured at 37 °C for ∼24 h to ∼90% confluence. Thereafter, binding assays were conducted in following steps: removal of the induction medium, addition of pre-warmed binding solution (serum-free DMEM plus 0.1% bovine serum albumin and 0.01% Tween-20) containing tracer and competitor (50 μL/well), incubation at room temperature for ∼1 h, further addition of diluted NanoLuc substrate (10 μL/well, 30-fold dilution of the substrate stock in binding solution), and measurement of bioluminescence on a SpectraMax iD3 plate reader (Molecular Devices). For saturation binding assays, the binding solution contained different concentrations of SmBiT-Lo-GPR15LG; for competition binding assays, the binding solution contained a constant concentration of SmBiT-Lo-GPR15LG and different concentrations of competitors. The measured bioluminescence data were expressed as mean ± standard deviation (SD, *n* = 3) and fitted to one-site binging model using the SigmaPlot 10.0 software (SYSTAT software).

## 3. Results

### Identification of possible fish GPR15LG orthologs

According to the NCBI gene database, the human *GPR15LG* gene is located adjacent to *GHITM, CDHR1, LRIT2*, and *LRIT1* and comprises three exons and two introns (Fig. S4). The gene synteny and genomic architecture may be evolutionarily conserved, thus we searched for potential fish GPR15LG orthologs in the NCBI gene database using these features. In the reference genome (fLepOcu1.hap2) of spotted gar (*Lepisosteus oculatus*) [33], we identified a candidate gene (*LOC107077293*) flanked by *ghitm* and *cdhr1a*, also consisting of three exons and two introns (Fig. S5). Although its transcript (XR_001478334) is annotated as a long non-coding RNA, sequence analysis revealed that it encodes a small secretory protein of 66 amino acids, here designated as Lo-GPR15LG (Table S2 and Fig. S6). Notably, this transcript contains several in-frame stop codons (Fig. S6), which may lead to nonsense-mediated mRNA decay and thus explain its mis-annotation. However, similar in-frame stop codons are also present in the human GPR15LG (Hs-GPR15LG) transcript (NM_207373), which is nonetheless protein-coding (Fig. S6), suggesting that XR_001478334 is likely translated *in vivo*.

The full-length Lo-GPR15LG contains a predicted N-terminal signal peptide of 24 residues and a 42-residue mature peptide featuring four cysteine residues arranged in a CC-C-C pattern (Table S2 and Fig. 1A). Using similar criteria, we identified two additional misannotated fish transcripts, XR_010805124 from *Amia ocellicauda* and XR_010328695 from *Anguilla rostrata*, both of which encode putative GPR15LG orthologs despite being annotated as non-coding RNAs (Table S2 and Fig. S7). Further BLAST analysis using these sequences retrieved seven additional fish homologs, all currently annotated as uncharacterized proteins with unknown functions (Fig. 1A and Table S2). Interestingly, the fish species *Huso huso* has two GPR15LG proteins that are likely encoded by two paralogous genes (Fig. 1A and Table S2).

**Fig. 1.**
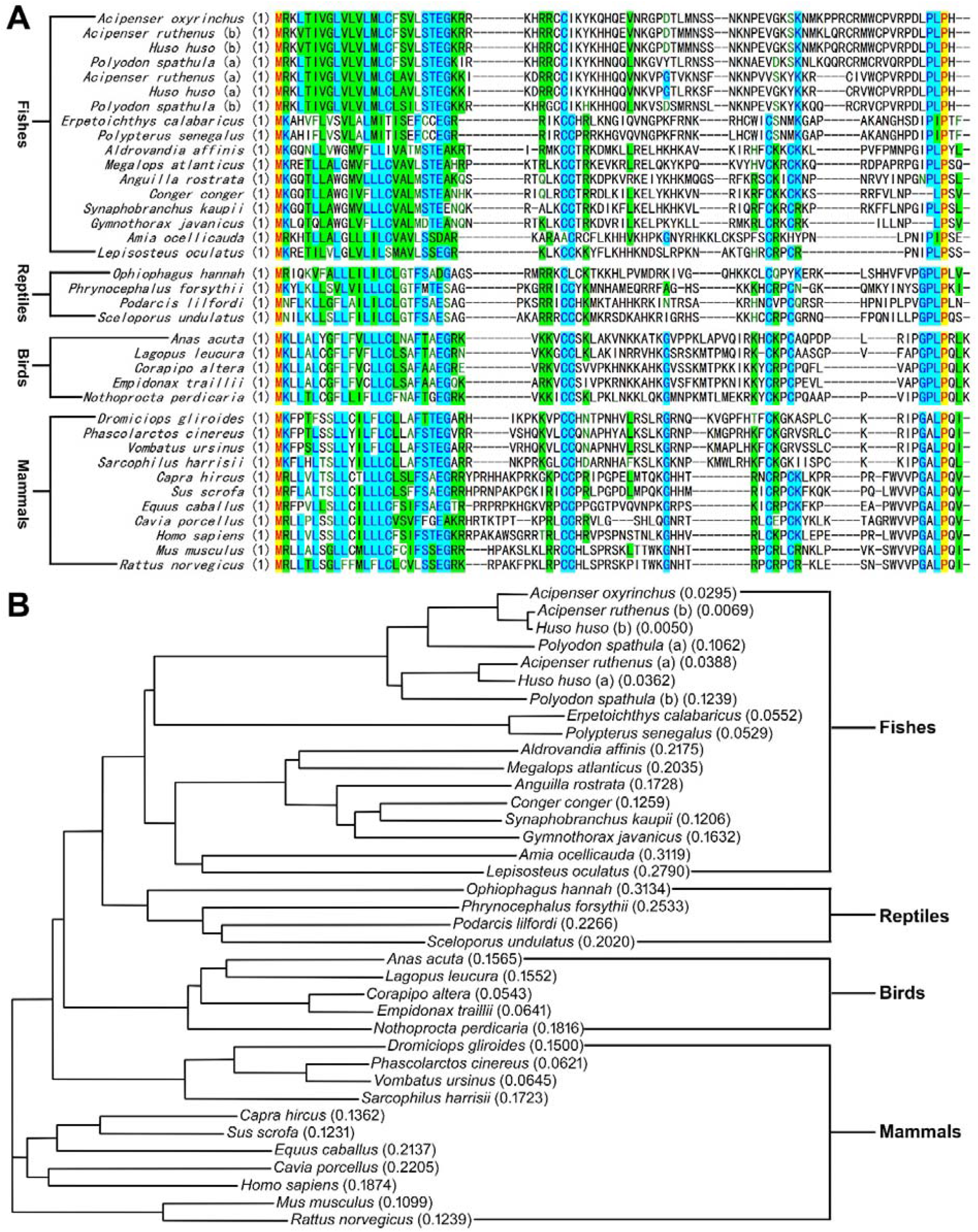
Identification of fish GPR15LG orthologs. (**A**) Amino acid sequence alignment of GPR15LG orthologs from mammals, birds, reptiles, and fishes. (**B**) Phylogenetic tree of the GPR15LG orthologs aligned in panel A. The letter “a” and “b” in parentheses indicate two GPR15LG paralogs in a species.

We also identified seven unannotated *gpr15lg* genes in five other fish species, including gray bichir (*Polypterus senegalus*), European conger (*Conger conger*), sterlet (*Acipenser ruthenus*), Mississippi paddlefish (*Polyodon spathula*), and reedfish (*Erpetoichthys calabaricus*), according to their gene synteny and RNA sequencing data (Table S2 and Fig. S8–S14). Via manually joining their exons, these previously unannotated fish GPR15LG orthologs were identified. Their genes are all adjacent to “*growth hormone-inducible transmembrane protein*” gene (*ghitm*), but four of them are composed of four exons, rather than three exons in most other species (Fig. S10–S13). Sterlet (*A. ruthenus*) and Mississippi paddlefish (*P. spathula*) have two paralogous *gpr15lg* genes in different chromosomes (Table S2), likely caused by genome duplication.

These fish GPR15LG orthologs lack significant overall sequence similarity to their counterparts from mammals, birds, and reptiles (Fig. 1A), thus they cannot be recognized previously. All GPR15LG orthologs exhibit substantial sequence divergence, particularly within the mature peptide, while retaining a relatively conserved C-terminal segment, including a highly conserved Leu–Pro dipeptide motif (Fig. 1A). This feature is consistent with the established importance of the C-terminal region in receptor binding and activation [41–43]. Phylogenetic analysis further showed that fish GPR15LG orthologs form a distinct cluster, positioned close to reptilian orthologs (Fig. 1B).

### Preparation of mature Lo-GPR15LG via bacterial overexpression

To obtain mature Lo-GPR15LG for functional characterization, we employed a bacterial overexpression strategy using a SUMO-fusion system, given the small size of the mature peptide (Fig. S1). Following IPTG induction in transformed *E. coli* cells, a prominent band of approximately 20 kDa was observed by SDS-PAGE (Fig. 2A, indicated by “f”), consistent with overexpression of the SUMO-fused precursor with a theoretical molecular weight of 17.4 kDa. However, the fusion protein exhibited susceptibility to proteolytic degradation during purification. To mitigate this, bacterial lysis was performed under denaturing conditions (6 M guanidine chloride) to inactivate endogenous proteases. In addition, the sulfhydryl groups of cysteine residues were reversibly modified via *S*-sulfonation to prevent disulfide crosslinking under denaturing conditions. After loading the denatured lysate onto a Ni^2+^ affinity column, on-column refolding of the SUMO-fused precursor was achieved by extensive washing with renaturation buffer. SDS-PAGE analysis confirmed that the soluble fusion protein could be efficiently eluted under native conditions (Fig. 2A).

**Fig. 2.**
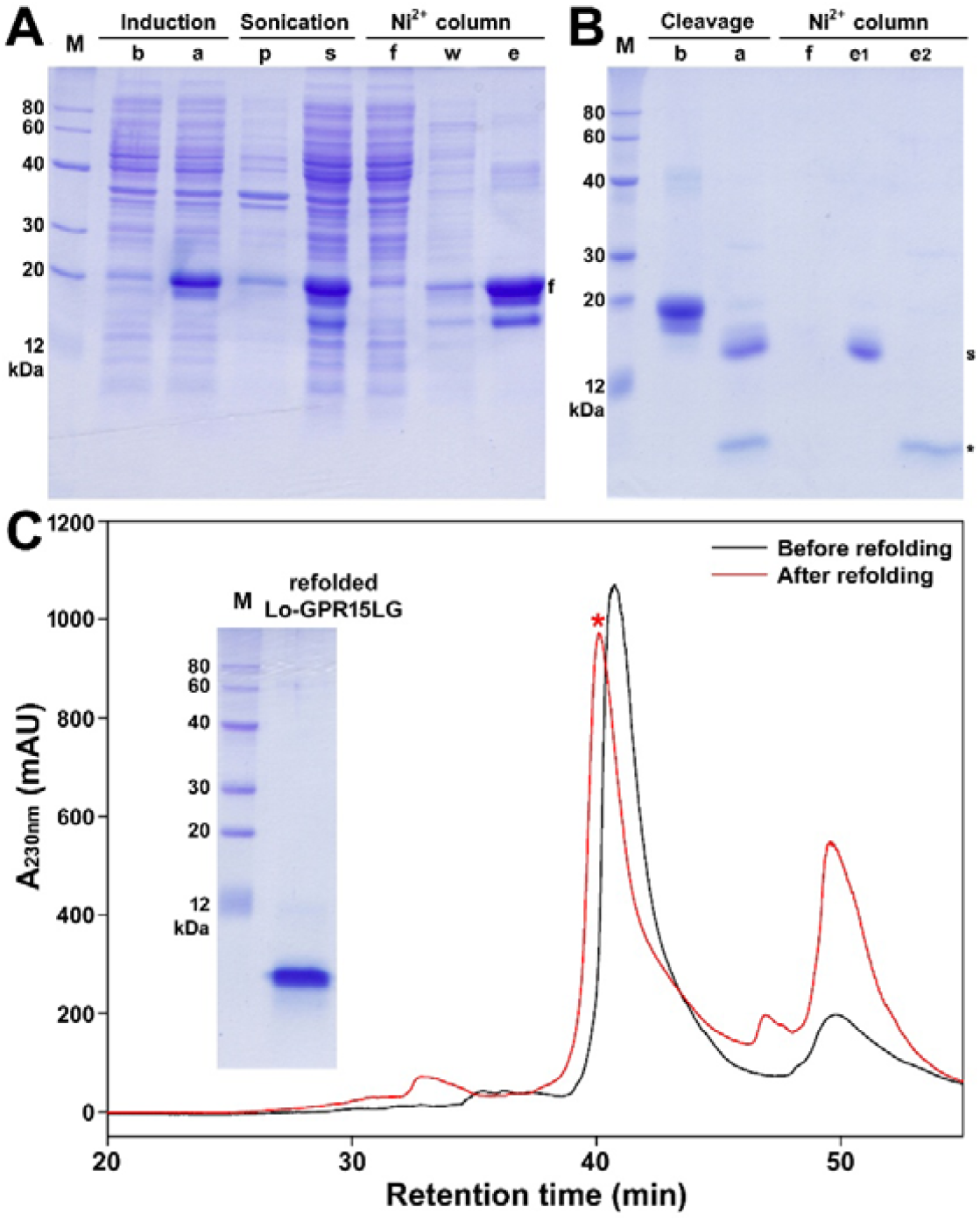
Preparation of recombinant Lo-GPR15LG. (**A**) SDS-PAGE analysis of the overexpressed 6×His-SUMO-Lo-GPR15LG at different preparation steps. Lane (M), protein ladder; lane (b), before IPTG induction; lane (a), after IPTG induction; lane (p), pellet after sonication; lane (s), supernatant after sonication; lane (f), flow-through faction from the Ni^2+^ column; lane (w), fraction washed by 10 mM imidazole; lane (e), fraction eluted by 250 mM imidazole. The letter “f” indicates band of the SUMO-fused precursor. (**B**) SDS-PAGE analysis of 6×His-SUMO-Lo-GPR15LG before or after SUMO protease treatment. Lane (M), protein ladder; lane (b), before SUMO protease treatment; lane (a), after SUMO protease treatment; lane (f), flow-through fraction from the Ni^2+^ column; lane (e1), fraction eluted by 250 mM imidazole in 20 mM phosphate buffer; lane (e2), fraction eluted by 250 mM imidazole in denaturing buffer. The letter “s” indicates the released 6×His-SUMO tag, and the asterisk indicates the released Lo-GPR15LG after SUMO protease treatment. In panel A and B, the samples were loaded onto a 15% reducing SDS-gel. After electrophoresis, the gel was stained by Coomassie Brilliant Blue R250. (**C**) HPLC analysis of Lo-GPR15LG before or after *in vitro* refolding. The major refolding peak (indicated by an asterisk) was manually collected, lyophilized, analyzed by SDS-PAGE, and used for activity assay. **Inner panel**, SDS-PAGE analysis of the refolded Lo-GPR15LG after HPLC purification.

Subsequently, the eluted SUMO-fused precursor was digested with recombinant SUMO protease to release Lo-GPR15LG. SDS-PAGE analysis revealed two major bands after cleavage (Fig. 2B): the upper band (labeled “s”) corresponding to the released 6×His-SUMO tag, and a lower band (indicated by an asterisk) corresponding to Lo-GPR15LG. To separate these components, Ni^2+^ affinity chromatography was employed. Interestingly, Lo-GPR15LG also exhibited affinity for the Ni^2+^ column despite lacking a 6×His tag. After optimization, the 6×His-SUMO fusion partner was first eluted with imidazole in 20 mM phosphate buffer, followed by elution of Lo-GPR15LG using imidazole in denaturing buffer (Fig. 2B).

The purified Lo-GPR15LG was then subjected to *in vitro* refolding to form correct intramolecular disulfide bonds. HPLC analysis showed a sharp dominant peak following refolding (Fig. 2C). This fraction corresponded to a single band on SDS-PAGE (Fig. 2C, inner panel), indicating high purity of the final product. These results demonstrated that recombinant Lo-GPR15LG can be efficiently produced through SUMO-assisted bacterial expression and subsequent purification, digestion, and refolding procedures.

### Efficient activation of Lo-GPR15 by recombinant Lo-GPR15LG measured via NanoBiT-based β-arrestin recruitment assay

To evaluate the activity of recombinant Lo-GPR15LG toward its corresponding receptor Lo-GPR15, we employed a NanoBiT-based β-arrestin recruitment assay, in which Lo-GPR15-LgBiT was coexpressed with SmBiT-ARRB2 in transfected HEK293T cells. Upon receptor activation, recruitment of SmBiT-ARRB2 to Lo-GPR15-LgBiT leads to complementation of the NanoBiT fragments and a corresponding increase in bioluminescence.

After transfected HEK293T cells were induced to coexpress Lo-GPR15-LgBiT and SmBiT-ARRB2, addition of NanoLuc substrate alone generated only a low basal signal. In contrast, subsequent addition of recombinant Lo-GPR15LG elicited a rapid and robust increase in bioluminescence in a dose-dependent manner (Fig. 3A), with an EC_50_ value of approximately 10 nM (Fig. 3A, inner panel), indicating efficient receptor activation. Specifically, truncation of four C-terminal residues markedly impaired ligand activity, as Lo-[desC4]GPR15LG produced only minimal responses even at 1000 nM (Fig. 3B), highlighting the critical role of the C-terminal region in receptor activation. In addition, cross-species activity was not observed: recombinant Hs-GPR15LG failed to activate Lo-GPR15 (Fig. 3C), and Lo-GPR15LG showed no detectable activity toward human GPR15 (Hs-GPR15) (Fig. 3D). These results demonstrate that Lo-GPR15LG potently and specifically activates its cognate receptor in a C-terminal-dependent manner, while ligand–receptor recognition is species-specific.

**Fig. 3.**
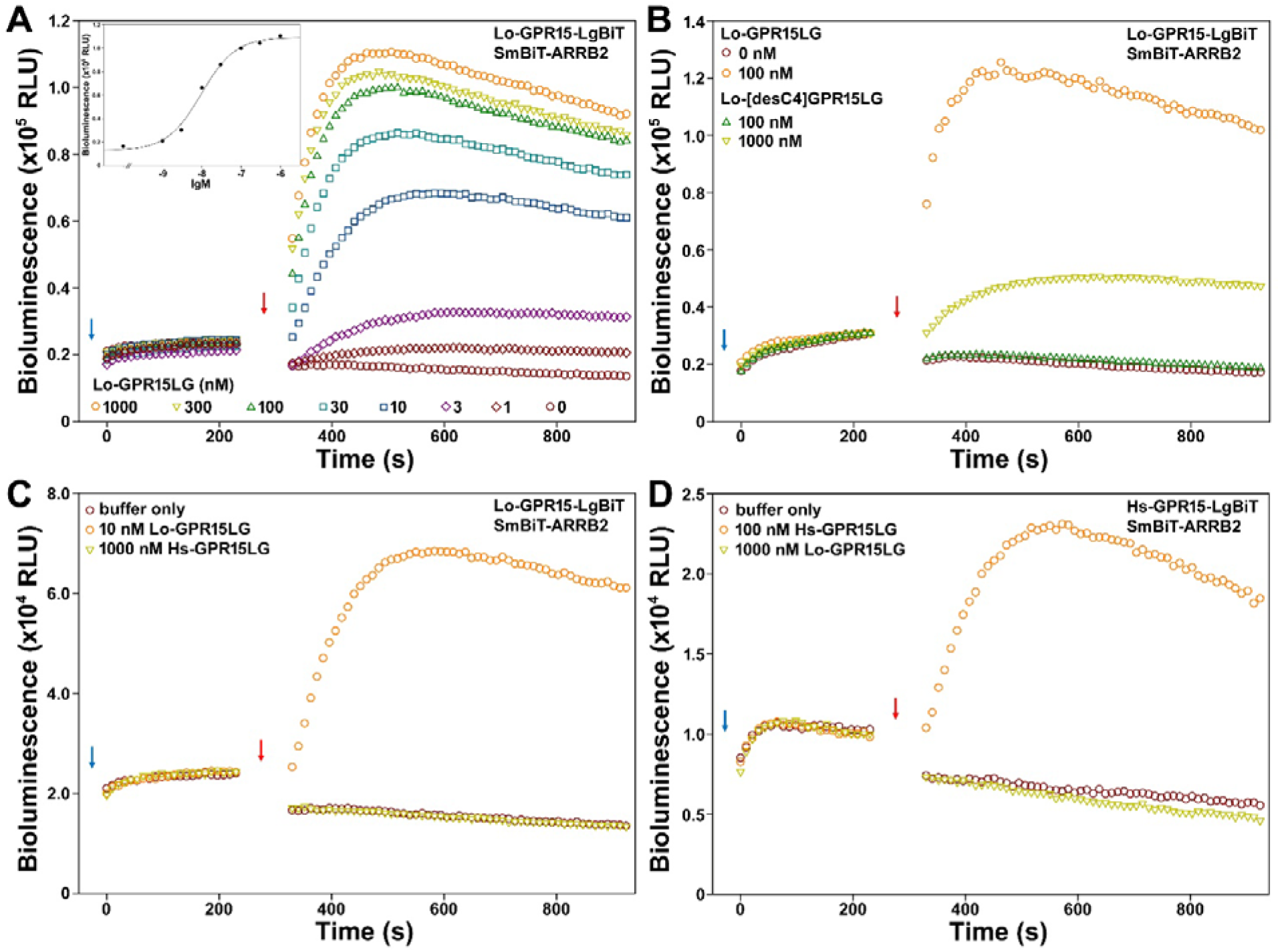
The NanoBiT-based β-arrestin recruitment assays. (**A**) Activation effect of recombinant Lo-GPR15LG towards Lo-GPR15. Inner panel, dose-response curve. (**B**) Activation effect of the C-terminally truncated Lo-[desC4]GPR15LG towards Lo-GPR15. (**C**) Activation effect of Hs-GPR15LG towards Lo-GPR15. (**D**) Activation effect of Lo-GPR15LG towards Hs-GPR15. In these β-arrestin recruitment assays, NanoLuc substrate and indicated peptide were sequentially added to living HEK293T cells coexpressing SmBiT-ARRB2 and indicated C-terminally LgBiT-fused GPR15 receptor, and bioluminescence was measured on an iD3 plate reader. Blue arrows indicate addition of NanoLuc substrate and red arrows indicate addition of peptide.

### Direct binding of recombinant Lo-GPR15LG with Lo-GPR15 measured via NanoBiT-based binding assay

To investigate the direct interaction between Lo-GPR15LG and its cognate receptor Lo-GPR15, we employed a NanoBiT-based homogeneous binding assay. In this system, a low-affinity SmBiT tag was fused to the N-terminus of Lo-GPR15LG, while a secretory LgBiT was fused to the extracellular N-terminus of Lo-GPR15. Ligand binding brings the two NanoBiT fragments into proximity, resulting in luciferase complementation and a measurable bioluminescence signal.

To assess whether the SmBiT tag affected ligand activity, SmBiT-Lo-GPR15LG was first evaluated using the β-arrestin recruitment assay. The tagged ligand induced a robust, dose-dependent increase in bioluminescence in HEK293T cells coexpressing Lo-GPR15-LgBiT and SmBiT-ARRB2 (Fig. 4A), with an EC_50_ (∼25 nM) comparable to that of untagged Lo-GPR15LG (∼10 nM) (Fig. 4A, inner panel), indicating minimal impact of the tag on ligand function.

**Fig. 4.**
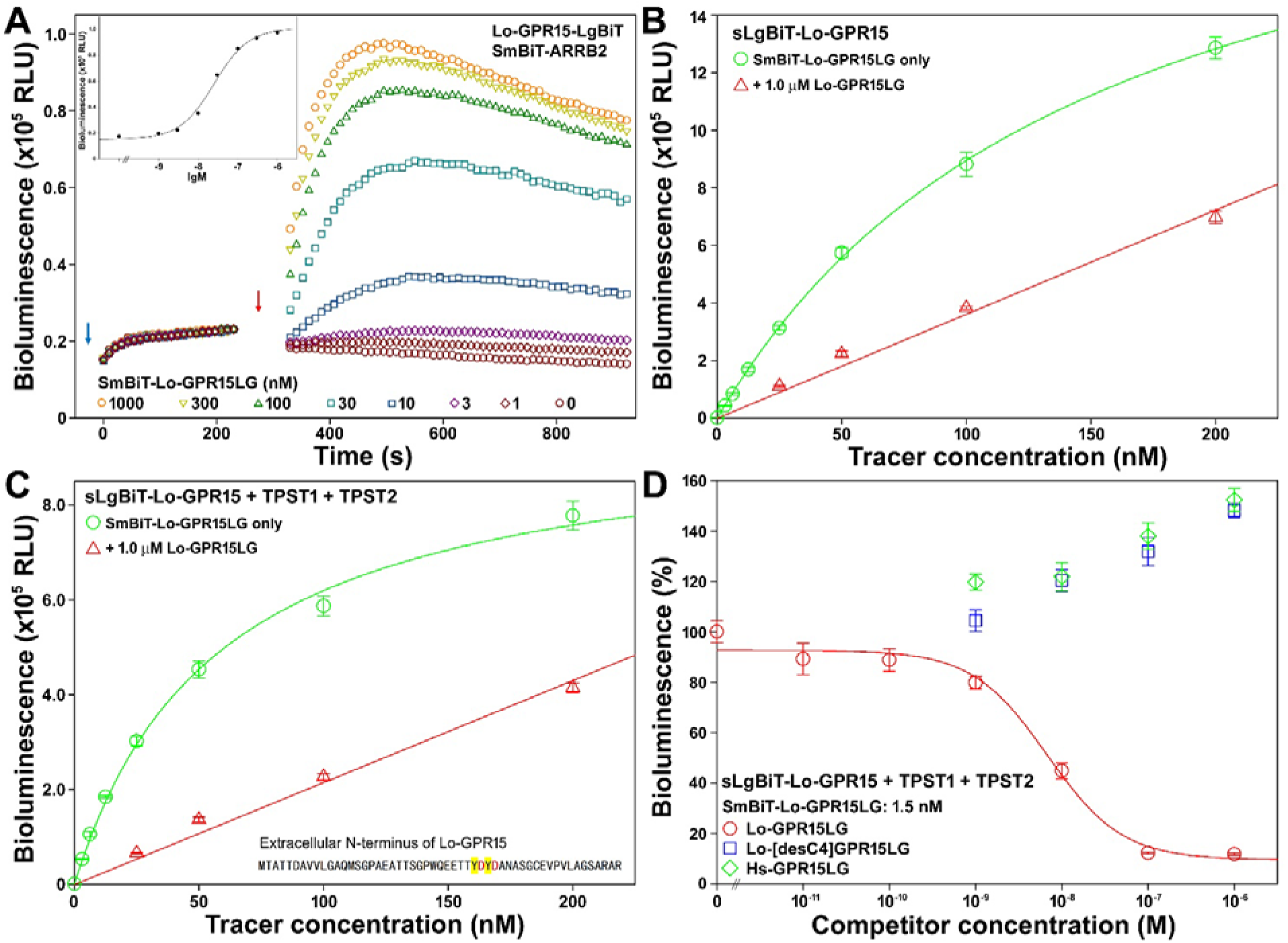
The NanoBiT-based ligand–receptor binding assays. (**A**) Activity of the recombinant SmBiT-Lo-GPR15LG towards Lo-GPR15 measured via the NanoBiT-based β-arrestin recruitment assay. **Inner panel**, dose-response curve. NanoLuc substrate and different concentrations of SmBiT-Lo-GPR15LG were sequentially added to living HEK293T cells coexpressing Lo-GPR15-LgBiT and SmBiT-ARRB2, and bioluminescence was measured on a plate reader. Blue arrow indicates addition of NanoLuc substrate, and red arrow indicates addition of peptide. (**B**,**C**) Saturation binding of SmBiT-Lo-GPR15LG with sLgBiT-Lo-GPR15 without (B) or with (C) coexpression of human TPST1 and TPST2. The measured bioluminescence data are expressed as mean ± SD (*n* = 3) and plotted using the SigmaPlot10.0 software. Total binding data (green circles) were fitted with the function of Y = B_max_X/(K_d_+X), non-specific binding data (red triangles) were fitted with linear curves. (**D**) The NanoBiT-based competition binding assays. The relative bioluminescence data are expressed as mean ± SD (*n* = 3) and fitted with sigmoidal curves using the SigmaPlot10.0 software.

In binding assays, SmBiT-Lo-GPR15LG produced a saturable increase in bioluminescence in cells expressing sLgBiT-Lo-GPR15 (Fig. 4B), with a dissociation constant (K_d_) of 153 ± 8 nM (*n* = 3). The presence of excess unlabeled Lo-GPR15LG (1.0 μM) markedly reduced the signal, confirming binding specificity. Notably, the extracellular N-terminus of Lo-GPR15 contains two tyrosine residues adjacent to negatively charged residues (Fig. 4C, inner panel), suggesting potential tyrosine sulfation [43,44]. Consistently, coexpression of TPST1 and TPST2 significantly enhanced binding affinity (Fig. 4C), reducing the K_d_ value to 57 ± 3 nM (*n* = 3).

In competition binding assays, Lo-GPR15LG inhibited binding in a sigmoidal manner with an IC_50_ value of 6.8 ± 1.1 nM (*n* = 3) (Fig. 4D). In contrast, Lo-[desC4]GPR15LG and Hs-GPR15LG showed no inhibitory effect, indicating a loss of binding capability. Interestingly, both Lo-[desC4]GPR15LG and Hs-GPR15LG caused an increase in bioluminescence in the competition binding assay (Fig. 4D). This strange phenomenon may result from recruitment of more tracers to the cell surface by the inactive peptide via interactions with both the tracer and cell surface glycosaminoglycans.

## Discussion

In this study, we identified 17 GPR15LG orthologs in 14 fish species for the first time based on conserved gene synteny and characteristic amino acid sequence features, despite the absence of significant overall sequence similarity to known GPR15LG orthologs from higher vertebrates. *GPR15LG* genes are typically located adjacent to *GHITM* and *CDHR1* and consist of three or four exons. At the protein level, GPR15LG orthologs exhibit several defining features, including a small size (generally <100 amino acids), the presence of an N-terminal signal peptide, and a highly basic mature peptide containing four cysteine residues and a conserved Leu/Ile–Pro dipeptide motif at the C-terminus (Fig. 1A). These conserved genomic and structural characteristics provide a robust framework for the identification of additional GPR15LG orthologs across diverse species in future studies.

The representative Lo-GPR15LG directly binds to and efficiently activates its cognate GPR15 receptor, indicating the presence of a functional GPR15LG–GPR15 signaling system in spotted gar. However, the physiological and pathological roles of this signaling axis in fish remain to be elucidated. The identification of a functional ligand–receptor pair in extant fish species suggests that this system likely originated in early vertebrate ancestors and has been conserved throughout evolution. These findings therefore support the notion that the GPR15LG–GPR15 signaling pathway has a long evolutionary history and may play fundamental roles across vertebrate lineages.

Mature GPR15LG orthologs are consistently highly basic proteins, with predicted isoelectric points typically exceeding 10 (Table S2). This pronounced positive charge is likely critical for their chemoattractant function, as it facilitates binding to negatively charged extracellular glycosaminoglycans, thereby establishing local concentration gradients that promote recruitment of GPR15-expressing immune cells. In addition, the basic nature of GPR15LG may enhance its interaction with receptor through electrostatic contacts with negatively charged residues in the extracellular region, including sulfated tyrosine residues in the N-terminus of GPR15 orthologs [43].

Mature GPR15LG orthologs exhibit remarkable sequence diversity, particularly within their N-terminal regions (Fig. 1A), yet retain the ability to recognize and activate their cognate receptors. Previous studies have highlighted the critical role of the relatively conserved C-terminal fragment, which inserts into the orthosteric ligand-binding pocket of the receptor [41–43]. However, the highly variable N-terminal region also contributes to receptor interaction, as evidenced by the markedly reduced activity (∼40-fold) of a synthetic C-terminal 11-mer peptide compared with the full-length protein [42]. This N-terminal region is therefore likely involved in receptor binding, primarily through electrostatic interactions with the extracellular domain of the receptor. Such multi-point electrostatic interactions appear to depend more on the overall abundance of positively charged residues in the ligand and negatively charged residues in the receptor, rather than precise amino acid sequence complementarity. Consistent with this model, GPR15LG orthologs are highly basic proteins enriched in lysine (K) and arginine (R) residues within their variable N-terminal regions (Fig. 1A), whereas GPR15 orthologs contain multiple negatively charged residues, including glutamate (E), aspartate (D), and potentially sulfated tyrosine (Y), within their variable extracellular N-termini (Table S3 and Fig. S15).

## CRediT authorship contribution statement

Jun-Ji Yao: Investigation, Methodology, Visualization. Jie Yu: Investigation, Methodology, Visualization. Hao-Zheng Li: Investigation, Methodology. Juan-Juan Wang: Investigation, Methodology. Ya-Li Liu: Formal analysis, Project administration. Zhan-Yun Guo: Supervision, Conceptualization, Writing - Original Draft, Writing - Review & Editing, Funding acquisition.

## Acknowledgments

This work was supported by grant from the National Natural Science Foundation of China (31971193).

## Supplementary data

Supplementary data for this article can be found online.

## Data availability

The data of this study are available in this manuscript, as well as the associated supplementary information.

