## Supplemental Table S1-S3 and Fig. S1-S15 for "Origin of the GPR15LG–GPR15 signaling axis in ancient fish ancestors"

### Contents:

**Table S1.** Primers and vectors for generating expression constructs for Lo-GPR15.

**Table S2.** Information about GPR15LG orthologs from mammals, birds, reptiles, and fishes.

**Table S3.** Information about GPR15 orthologs from mammals, birds, reptiles, and fishes.

**Fig. S1.** The nucleotide sequence and amino acid sequence of the Lo-GPR15LG expression constructs.

**Fig. S2.** The nucleotide sequence and amino acid sequence of the overexpressed R3-ULP1.

**Fig. S3.** The nucleotide sequence and amino acid sequences of Lo-GPR15 expression constructs.

**Fig. S4.** Gene synteny and genomic architecture of human *GPR15LG* in human genome.

**Fig. S5.** Gene synteny and genomic architecture of the possible fish *gpr15lg* gene (LOC107077293) in the reference genome (fLepOcu1.hap2) of spotted gar (*Lepisosteus oculatus*).

**Fig. S6.** The cDNA sequence and its encoded protein sequence of human or spotted gar GPR15LG.

**Fig. S7.** The cDNA sequence and its encoded protein sequence of possible fish GPR15LG orthologs from *Amia ocellicauda* (**A**) or *Anguilla rostrata* (**B**).

**Fig. S8.** Information about possible GPR15LG from *Polypterus senegalus* (gray bichir).

**Fig. S9.** Information about possible GPR15LG from *Conger conger* (European conger).

**Fig. S10.** Information about possible GPR15LG (isoform a) from *Acipenser ruthenus* (sterlet).

**Fig. S11.** Information about possible GPR15LG (isoform b) from *Acipenser ruthenus* (sterlet).

**Fig. S12.** Information about possible GPR15LG (isoform a) from *Polyodon spathula* (Mississippi paddlefish).

**Fig. S13.** Information about possible GPR15LG (isoform b) from *Polyodon spathula.* (Mississippi paddlefish).

**Fig. S14.** Information about possible GPR15LG from *Erpetoichthys calabaricus* (reedfish).

**Fig. S15.** Amino acid sequence alignment of GPR15 orthologs from different species.

**Table S1.** Primers and vectors for generating expression constructs for Lo-GPR15.

| Expression constructs | Vectors for cloning | Enzymes for vector cleavage | PCR primers (5' to 3') | PCR template | Approach for construct generation |
|---|---|---|---|---|---|
| pcDNA3.1/ Lo-GPR15 | pcDNA3.1(+) | NheI NotI | | Chemical synthesis | Gibson assembly |
| PB-TRE/ sLgBiT-Lo-GPR15 | PB-TRE/ sLgBiT-GPR182 | KpnI AgeI (removing GP182) | Forward: GGT GGC AGC GGC GGT GGT ACC ACC GCC ACC ACC GAC GCC Reverse: GAG GCT GAT CAG CGG GT T TCA GTC CTG CAG GGT GGT CAG | pcDNA3.1/ Lo-GPR15 | Gibson assembly |
| pTRE3G-BI/ Lo-GPR15-LgBiT: SmBiT-ARRB2 | pTRE3G-BI/ GPR83-LgBiT: SmBiT-ARRB2 | EcoRI AgeI (removing GPR83) | Forward: GAA CCG TCA GAT CGC CTG GAG AAT TCG GGG AGA CCC AAG CTG GCT AGC Reverse: GCT AGA CCC TCC GCC GGT ACC GAC CGG TGG GTC CTG CAG GGT GGT CAG | pcDNA3.1/ Lo-GPR15 | Gibson assembly |

PB-TRE/sLgBiT-GPR182 was generated in our laboratory based on PB-TRE/dCas9-VPR (Addgene cat#: 63800) via removal of the dCas9-VPR fragment by NheI and PmeI cleavage and then ligation of sLgBiT-GPR182 fragment, unpublished data.

pTRE3G-BI/GPR83-LgBiT:SmBiT-ARRB2 was generated in our previous study.

For synthetic PCR primers, the fragment pairing with vector is highlighted in yellow, the fragment pairing with PCR template is underlined.

**Table S2.** Information about GPR15LG orthologs from mammals, birds, reptiles, and fishes. The information is downloaded from the NCBI gene database or protein database. The fish GPR15LG orthologs were newly identified in this study. Different amino acids between two paralogs are shown in red.

| Groups | Species | NCBI information | Amino acid sequence (signal peptide shaded) | pI of mature peptide |
|---|---|---|---|---|
| Mammals | *Homo sapiens* (human) | Gene ID: 387695<br>mRNA ID: NM_207373<br>Protein ID: NP_997256 | MRLLVLSSLLCILLLCFSIFSTEGKRRPAKAWSGRRTRLCCHRVPSPNSTNLKGHHVRLCKPCKLEPEPRLWVVPGALPQV | 11.8 |
| | *Mus musculus* (house mouse) | Gene ID: 70045<br>mRNA ID: NM_001206684<br>Protein ID: NP_001193613 | MRLLALSGLLCMLLLCFCIFSSEGRRHPAKSLKLRRCCHLSPRSKLTTWKGNHTRPCRLCRNKLPVKSWVVPGALPQI | 12.2 |
| | *Rattus norvegicus* (Norway rat) | Gene ID: 290595<br>mRNA ID: NM_001106063<br>Protein ID: NP_001099533 | MRLLTLSGLFFMLFLCLCVLSSEGRKRPAKFPKLRPCCHLSPRSKPITWKGNHTRPCRPCRKLESNSWVVPGALPQI | 12.0 |
| | *Sus scrofa* (pig) | Gene ID: 100511838<br>mRNA ID: NM_001243901<br>Protein ID: NP_001230830 | MRFLALTSLLCILLLCLSFFSAEGRRHPRNPAKPGKIRICCPRLPGPDLMPQKGHHMRICRPCKFKQKPQLWVVPGALPQV | 12.0 |
| | *Equus caballus* (horse) | Gene ID: 102149943<br>mRNA ID: NM_001433570<br>Protein ID: NP_001420499 | MRFPVLLSLLCILLLCFSIFSAEGTRPRPRPKHGKVRPCCPPGGTPVQVNPKGRPSKICRPCKFKPEAPWVVPGALPQV | 11.5 |
| | *Capra hircus* (goat) | Gene ID: 102190585<br>mRNA ID: XM_005699304<br>Protein ID: XP_005699361 | MRLLVLTSLLCTLLLCLSLFSAEGRRYPRHHAKPRKGKPCCPRIPGPELMTQKGHHTRNCRPCKLKPRPRFWVVPGALPQV | 12.0 |
| | *Cavia porcellus* (domestic guinea pig) | Gene ID: 101787781<br>mRNA ID: XM_005002068;<br>Protein ID: XP_005002125 | MRLLPLSSLLCILLLCVSVFFGEAKRHRTKTPTKPRLCCRRVLGSHLQGNRTRLCEPCKYKLKTAGRWVVPGALPQV | 11.8 |
| | *Vombatus ursinus* (common wombat) | Gene ID: 114039903<br>mRNA ID: XM_027857826<br>Protein ID: XP_027713627 | MKFPSLSSLLYILFLCLLAFSTEGARRVRHQKVLCCQNAPNHVLRSLKGRNPKMAPLHKFCKGRVSSLCKRIPGALPQI | 12.2 |
| | *Sarcophilus harrisii* (Tasmanian devil) | Gene ID: 111718615<br>mRNA ID: XM_023494623<br>Protein ID: XP_023350391 | MKFLHLTSLLYILLLCLLAFSTEGARRNKPRKGLCCHDARNHAFKSLKGKNPKMWLRHKFCKGKIISPCKKIPGALPQL | 11.3 |
| | *Phascolarctos cinereus* (koala) | Gene ID: 110215496<br>mRNA ID: XM_020997060<br>Protein ID: XP_020852719 | MKFPTLSSLLYILFLCLLAFSTEGVRRVSHQKVLCCQNAPHYALKSLKGRNPKMGPRHKFCKGRVSRLCKRIPGALPQI | 12.0 |
| | *Dromiciops gliroides* (monito del monte) | Gene ID: 122739871<br>mRNA ID: XM_043981580<br>Protein ID: XP_043837515 | MKFPTFSSLLCILFLCLLAFTTEGARHIKPKKVPCCHNTPNHVLRSLRGRNQKVGPFHTFCKGKASPLCKRIPGALPQI | 11.6 |
| Birds | *Anas acuta* (northern pintail) | Gene ID: 137859209<br>mRNA ID: XM_068687991<br>Protein ID: XP_068544092 | MKLLALYGFLFVLLLCLNAFTAEGRKVKKVCCSKLAKVNKKATKGVPPKLAPVQIRKHCKPCAQPDPLRIPGPLPRLK | 11.1 |
| | *Nothoprocta perdicaria* | Gene ID: 112950473<br>mRNA ID: XM_026043014<br>Protein ID: XP_025898799 | MKLLTLCGFLLIFLLCFNAFTGEGRKVKKICCSKLPKLNKKLQKGMNPKMTLMEKRKYCKPCPQAAPPVIPGPLPQLK | 10.7 |
| | *Lagopus leucura* (white-tailed ptarmigan) | Gene ID: 122175628<br>mRNA ID: XM_042863405<br>Protein ID: XP_042719339 | MKLLALCGFLFVFLLCLSAFTAEGRNVKKGCCLKLAKINRRVHKGSRSKMTPMQIRKCKPCAASGPVFAPGPLPQLK | 11.6 |
| | *Empidonax traillii* | Gene ID: 114063575 | MKLLALCGFLFVCLLCLSAFAAEGQKARKVCCSIV | 10.8 |

| | | | | |
|---|---|---|---|---|
| | (willow flycatcher) | mRNA ID: XM_027895931;<br>Protein ID: XP_027751732 | PKRNKKAHKGVSFKMTPKNIKKYCRPCPEVLVAPG<br>PLPQLK | |
| | *Corapipo altera* (White-ruffed manakin) | Gene ID: 113950057<br>mRNA ID: XM_027649348<br>Protein ID: XP_027505149 | MKLLALCGFLFVCLLCLSAFAAEGREVRKVCCSV<br>VPKHNKKAHKGVSSKMTPKKIKKYCRPCPQFLVAP<br>GPLPQLK | 10.8 |
| Reptiles | *Sceloporus undulatus* (fence lizard) | Gene ID: 121926506<br>mRNA ID: XM_042459530<br>Protein ID: XP_042315464 | MNILKLLSLLFILILCLGTFSAESAGAKARRRCCCK<br>MKRSDKAHKRIGRHSKKHCCRPCGRNQFPQNILLP<br>GPLPQS | 11.5 |
| | *Podarcis lilfordi* | Protein ID: CAI5774416 | MNFLKLLGFLAILILCLGTFSAESAGPKSRRICCHK<br>MKTAHHKRKINTRSAKRHNCVPCQRSRHPNIPLPV<br>GPLPLN | 12.2 |
| | *Ophiophagus hannah* | Protein ID: ETE69020 | MRIQKVFALLLILILCLGTFSADGAGSRMRRKCLC<br>KTKKHLPVMDRKIVGQHKKCLCQPYKERKLSHHV<br>FVPGPLPLV | 10.9 |
| | *Phrynocephalus forsythii* | Protein ID: JRQ81_017375 | MKYLKLLSVLVILLLCLGTFMTESAGPKGRRICCY<br>KMNHAMEQRRFAGHSKKKHCRPCNGKQMKYINY<br>SGPLPKI | 10.7 |
| Fishes | *Lepisosteus oculatus* (spotted gar) | Gene ID: 107077293<br>RNA ID: XR_001478334 | MKRETILVLGLVLILSMAVLSSEGRKLKCCKKYFL<br>KHHKNDSLRPKNAKTGHRCRPCRPNIPLPSS | 11.1 |
| | *Amia ocellicauda* | Gene ID: 136716144<br>RNA ID: XR_010805124 | MRKHTLLALGLLLILCVAVLSSDARKARAACRCFL<br>KHHVKHPKGNYRHKKLCKSPFSCRKHYPNLPNIPI<br>PSE | 10.9 |
| | *Anguilla rostrata* (American eel) | Gene ID: 135249475;<br>RNA ID: XR_010328695 | MKGQTLLAWGMVLLLCVALMSTEAKQSRTQLKC<br>CTRKDPKVRKEIYKHKMQGSRFKRSCKICKNPNRV<br>YINPGNPLPSL | 11.1 |
| | *Huso huso* | Protein ID: KAK6482051<br>(isoform a) | MRKLTIVGLVLVLMLCLAVLSTEGKKIKDRRCCIK<br>YKHHQQVNKVPGTLRKSFNKNPEVGKYKKRCRV<br>WCPVRPDLPLPH | 10.9 |
| | | Protein ID: KAK6487530<br>(isoform b) | MRKVTIVGLVLVLMLCFSVLSTEGKRRKHRRCCIK<br>YKHHQEVNKGPDTMMNSSNKNPEVGKSKNMKPQ<br>RCRMWCPVRPDLPLPH | 10.9 |
| | *Acipenser oxyrinchus oxyrinchus* | Protein ID: KAK1166446 | MRKLTIVGLVLVLMLCFSVLSTEGKRRKHRRCCIK<br>YKQHQEVNRGPDTLMNSSNKNPEVGKSKNMKPP<br>RCRMWCPVRPDLPLPH | 12.1 |
| | *Synaphobranchus kaupii* | Protein ID: KAJ8365699 | MKGQTLLAWGMVLLLCVALMSTEENQKRALKCC<br>TRKDIKFLKELHKHKVLRIQRFCKRCRKPRKFFLNP<br>GIPLPSL | 11.1 |
| | *Aldrovandia affinis* | Protein ID: KAJ8406799 | MKGQNLLVWGMVFLLIVATMSTEAKRTIRMKCCT<br>RKDMKLLRELHKHKAVKIRHFCKKCKKLPVFPMN<br>PGIPLPYL | 11.1 |
| | *Gymnothorax javanicus* | Protein ID: KAJ8264355 | MKLQTQLAWGLVFLLCVALMDTEANQNRTKLKC<br>CTRKDVRILKELPKYKLLRMKRLCRKCRKILLNPL<br>PSV | 11.3 |
| | *Megalops atlanticus* | Protein ID: KAG7492735 | MKRETLLALGMVFLLCVAVLSTEAHRPKTRLKCCT<br>RKEVKRLRELQKYKPQKVYHVCKRCKKQRDPAPR<br>PGIPLPSQ | 11.1 |
| | *Polypterus senegalus* (gray bichir) | Unannotated (Fig. S8)<br>Genomic sequence: NC_053154.1; Chromosome 1; Reference ASM1683550v1 | MKAHIFLVSVLALMITISEFCCEGRRIRCCPRRKHG<br>VQVNGPKFRNKLHCWICSNMKGAPAKANGHPDIP<br>IPTF | 9.89 |

| | | | | |
|---|---|---|---|---|
| | | Primary Assembly; Position: 255,964,400-255,952,700 | | |
| | *Conger conger* (European conger) | Unannotated (Fig. S9) Genomic sequence: NC_083761.1; Chromosome 2; Reference fConCon1.1; Position: 12,906,900-12,908,900 | MKGQTLLAWGIVFLLCVALMSTEANHKRIQLRCCTRRDLKILKELYKHKVNRIKRFCKKCKKSRRFVLNPLPSV | 10.64 |
| | *Acipenser ruthenus* (sterlet) | Unannotated (Fig. S10) Genomic sequence: NC_081201.1; Chromosome 13; Reference fAciRut3.2 maternal haplotype Primary Assembly; Position: 28,136,100-28,131,800; (Isoform a; four exons) | MRKLTIVGLVLVLMLCLAVLSTEGKKIKDRRCCIKYKHHQQVNKVPGTVKNSFNKNPVVSKYKKRCIVWCPVRPDLPLPH | 10.16 |
| | | Unannotated (Fig. S11) Genomic sequence: NC_081195.1; Chromosome 7; Reference fAciRut3.2 maternal haplotype Primary Assembly; Position: 13,538,300-13,545,500; (isoform b, four exons) | MRKVTIVGLVLVLMLCFSVLSTEGKRRKHRRCCIKYKHHQEVNKGPDTMMNSSNKNPEVGKSKNMKLQRCRMWCPVRPDLPLPH | 10.26 |
| | *Polyodon spathula* (Mississippi paddlefish) | Unannotated (Fig. S12) Genomic sequence: NC_054543.1; Chromosome 10; Reference ASM1765450v1 Primary Assembly; Position: 12,726,800-12,732,700; (isoform a, four exons) | MRKLTIVGLVLVLMLCFSVLSTEGKIRKHRRCCIKYKHNQQLNKGVYTLRNSSNKNAEVDKSKNLKQQRCRMCRVQRPDLPLPH | 10.46 |
| | | Unannotated (Fig. S13) Genomic sequence: NC_054546.1; Chromosome 13; Reference ASM1765450v1 Primary Assembly; Position: 12,073,000-12,077,100; (isoform b, four exons) | MRKLTIVGLVLVLMLCLSILSTEGKKRKHRGCCIKHKHHQQLNKVSDSMRNSLNKNPEVSKYKKQQRCRVCPVRPDLPLPH | 10.32 |
| | *Erpetoichthys calabaricus* (reedfish) | Unannotated (Fig. S14) Genomic sequence: NC_041395.1; Chromosome 2; Reference fErpCal1.1 Primary Assembly; Position: 250,459,800-250,451,800 | MKAHVFLVSVLALMITISEFCCEGRRIKCCHRLKNGIQVNGPKFRNKRHCWICSNMKGAPAKANGHSDIPIPTF | 9.82 |

**Table S3.** Information about GPR15 orthologs from mammals, birds, reptiles, and fishes. The information is downloaded from the NCBI gene database and aligned in Fig. S15.

| Groups | Species | NCBI information | Amino acid sequence |
|---|---|---|---|
| Mammals | *Homo sapiens* | Gene ID: 2838<br>RNA ID: NM_005290<br>Protein ID: NP_005281 | MDPEETSVYLDYYYATSPNSDIRETHSHVPYTSVFLPVFYTAVFLTGVLGNLVLMGALHFKPGSRRLIDIFIINLA<br>ASDFIFLVTLPLWVDKEASLGLWRTGSFLCKGSSYMISVNMHCSVLLLTCMSVDRYLAIVWPVVSRKFRRTDCAYV<br>VCASIWFISCLLGLPTLLSRELTLIDDKPYCAEKKATPIKLIWSLVALIFTFFVPLLSIVTCYCCIARKLCAHYQQ<br>SGKHNKKLKKSIKIIFIVVAAFLVSWLPFNTFKFLAIVSGLRQEHYLPSAILQLGMEVSGPLAFANSCVNPFIYYI<br>FDSYIRRAIVHCLCPCLKNYDFGSSTETSDSHLTKALSTFIHAEDFARRRKRSVSL |
| | *Equus caballus* | Gene ID: 100072420<br>RNA ID: XM_005602082<br>Protein ID: XP_005602139 | MDPEATSAYLDYFYATSQNPEIEEARSPVPYTSIFLPIFYTAVFLAGVLGNLVLMSALHFKRGSRRLIDIFIINLA<br>ASDFIFLITLPLWVDKEASSGLWRTGSFLCKASSYMITVNMHCSVFLLTCMSVDRYLAIVCPAISRKFRRRDCAYG<br>VCASVWFISCLLGLPTLLSRELTMIGDKPYCAEKRATSVKWAWSLVALIFTFFAPLVSIVTCYCCIARKLCAHYQQ<br>SGKHNKKLRKSIKIILIVVAAFVFSWLPFNTFKLLSIVSGLQQELYISQDFLKHGMDVSGPLAFANSCVNPFIYYV<br>FDGYIRRAIVRCLCPCLKNYDFGSSTETSDSHLTKVLSNFVHAEDFARRRKRSVSL |
| | *Pipistrellus kuhlii* | Gene ID: 118723163<br>RNA ID: XM_036445209<br>Protein ID: XP_036301102 | MDPEATSDYWDYSSGTSPNPDDVEETRFQAPYTSLFLPAFYTAVFLIGVLGNLLLMGVLHFKPGSRRLIDIFIVNL<br>AASDFIFLITLPLWVDKEASSGLWRTGSFLCKGSSYMISVNMHCSVFLLTCMSADRYLAIVWPAASRKLRRRDCAY<br>GVCASVWCVSCLLGLPTLLSRELTLIQDKPYCAEKRATSVKLTWALVSLLFTFFAPLVSIVTCYCCIIRKLCAQYQ<br>RTGKHNKKLRKSIKIIFIVVSAFVISWLPFNTFKLLAIVSGLQQEHYVSVSVLQRGMEVSGPLAFANSCVNPFIYY<br>IFDSYIRRATVQCLCPGLKSAADFGSSTETSDSHLTKALSNFIQAGDFARRRKRSVSL |
| | *Myotis davidii* | Gene ID: 102755765<br>RNA ID: XM_006769285<br>Protein ID: XP_006769348 | MDPEATSVYWDYVYGTSPNPDDLEETHFHAPYTSVFLPIFYATVFLLGALGNLLLMSVLHFKRGSRRLIDIFILNL<br>AASDFIFLITLPLWVDKEASSGLWRTGSFLCKGSSYMISVNMHCSVFLLTCMSIDRYLAIVCPAVARKFRRRDCAY<br>GVCASVWFVSCLLGLPSLLSRELILINDKPYCAEKSATSLKLTWALVSLIATFFAPLVSIVTCYCCIARKLCAHYQ<br>QTGKHNKKLRKSIKIILIVVAAFVFSWLPFNTFKLLAIVSRLQQEHYFSSSIPQLGMEVSGPLAFANSCVNPFIYY<br>IFDSYIRRAVVHCLCPGLKNYDFGSSTETSDSHLTKALSNFIHAGDFARRRKRSVSL |
| | *Molossus molossus* | Gene ID: 118617384<br>RNA ID: XM_036243290<br>Protein ID: XP_036099183 | MDPEATSVYSDYSYVTSPSPDEMERIHAHVPYTSVFLPIFYTAVFLIGVLGNLLLMSVLHFKRGSRRLIDIFIINL<br>AASDFIFLITLPLWVDKEASLGLWRTGSFLCKGSSYMISVNMHCSIFLLTCMSVDRYLAIVRPPVSKKVRRRDYAY<br>GVCATVWFISCLLGLPTLLSRELTLIKDKPYCAEKRATSIKLTWALVSLIFTFFTPLVSIVTCYCCIARKLCAHYR<br>QTGKHKKKLRKSIKIIFIVVAAFVFSWLPFNTFKLLAIVSGLQQKLCFSSSSLQLGMEVSGPLAFANSCVNPFIYY<br>IFDSYIRRATVHCLCPGLKNYDLGNSTESSDSHLTKAFSNFIHAGDFARKRKRSVSL |
| | *Artibeus jamaicensis* | Gene ID: 119047311<br>RNA ID: XM_037142718<br>Protein ID: XP_036998613 | MDPETTSIYLDYFYTTSQSPDDIEDTQSHVTYTSVFLPVFYTTVFLAGVLGNLVLMSALYFKRGSRRLMDIFILNL<br>AASDFIFLVTLPLWVDKEASSGLWRTGSFLCKGSSYVISVNMHCSVFLLTCMSVDRYLAIVRPTVSRRFRRKDCAY<br>VVCASVWFISCLLGLPTVLSRELTLIKDKPYCAEKKATSAKLTWALVSLIFTFFAPLVSIVTCYCCITRKLCVHYQ<br>QPGKHNKKLRKSIKIIFIVVSAFVFSWLPFNTFKLLAIVSGFQQEFYSSSSFSQVGMEVSGALAFANSCVNPFIYY<br>IFDSYIRRAIVHCLCPGLKNYDFGSSTETSDSHLAKALSNLVHAEDFARRRKRSVSL |
| | *Phoca vitulina* | Gene ID: 116633517<br>RNA ID: XM_032406123<br>Protein ID: XP_032262014 | MDPEATSVYLDYFYATSKNPDLEDTHSHVPYTSVFLPVFYTAVFLTGVLGNLVLMSALHFKRGSRRLIDIFIINLA<br>VSDFIFLVTLPLWVDKEASLGLWRTGSFLCKGSSYMISVNMHCSVFLLTCMSVDRYLAIVCPAISRKFRRRDCAYG<br>VCASVWFISCLLGLPTLLSRELTLIDDKPYCAEKRATSMKLTWALVALIFTFFVPLVSIVTCYCCITRKLCAHYQQ<br>SGKHNRKLRKSIKIILIVVAAFVFSWLPFNTFKLLAIVSGLQQEVYFSSAFLQLGMEVSGPLAFANSCVNPFIYYI<br>FDSYIRRAIVHCLCPCIKNYDFGSSTETSDSHLAKALSNFIHVEDFTRRRKRSVSL |
| | *Neogale vison* | Gene ID: 122910398<br>RNA ID: XM_044255025<br>Protein ID: XP_044110960 | MDPEATSVYLDYFYATSKNPDLEDTHSHVPYTSIFLPVFYTAVFLTGALGNLILMSALHFKRGSRRLIDIFIINLA<br>VSDFIFLVTLPLWVDKEASLGLWRTGSFLCKGSSYMISVNMHCSVFLLTCMSVDRYLAIMCPAISRKFRRRDCAYG<br>VCASVWFISCLLGLPTLLSRELTLIDGKPYCAEKRATSTKLTWALVALIFTFFVPLVSIVTCYCCITRKLCAHYQQ<br>AGKHNRKLRKSIKIIFIVVAAFVFSWLPFNTFKLLAIVSGLQQEVYFSSAFLQLGMEVSGPLAFANSCINPFIYYI<br>FDSYIRRAIVHFLCPCVKNYDFGSSTETSDSHLTKVLSNIIHAEDVARRRKRSVSL |
| | *Mustela erminea* | Gene ID: 116595224<br>RNA ID: XM_032351355<br>Protein ID: XP_032207246 | MDPEATSVYLDYFYATSKNPDLEDTHSHVPYTSIFLPVFYTAVFLTGALGNLILMSALHFKRGSRRLIDIFIINLA<br>VSDFIFLVTLPLWVDKEASLGLWRTGSFLCKGSSYMISVNMHCSVFLLTCMSVDRYLAIVCPAISRKFRRRDCAYG<br>VCASVWFISCLLGLPTLLSRELTLIDGKPYCAEKRATSTKLTWALVALIFTFFVPLVSIVTCYCCITRKLCAHYQQ<br>AGKHNRKLRKSIKIIFIVVAAFVFSWLPFNTFKLLAIVSGLQQEVYFSSAFLQLGMEVSGPLAFANSCINPFIYYI<br>FDSYIRRAIVHCLCPCVKNYDFGSSTETSDSHLTKVLSNIIHAEDFARKRKRSVSL |
| | *Manis pentadactyla* | Gene ID: 118927868<br>RNA ID: XM_036916554<br>Protein ID: XP_036772449 | MDPEATIVYLDYFYATSPNPEIEETHSHVPYTSVFLPIFYTAVFLIGVLGNLLLLSALYFKKGSRRLIDIFIINLA<br>ASDFIFLVTLPLWVDKEASLGLWRTGSFLCKGSSYMISVNMHCSVFLLTCMSVDRYVTIVCPALSRKFRRKDCAYG<br>VCASVWFISCLLGLPTLLSRELTLIDSKPYCAEKRATSTKLTWALVALIFTFFVPLVSIMACYCSITRKLCAHYQQ<br>SGKHNKKLRKSMKIIFIVVAAFVFSWLPFNTFKLLAVVSGLQRELSFSSAFLQLGMEVSGPLAFANSCVNPFIYYI<br>FDSYIRRAILHCLCPCVKNYEFGSSTETSDSHLTKALSNVIHAEDFSRRRKRSVSL |
| | *Hyaena hyaena* | Gene ID: 120223199<br>RNA ID: XM_039221232<br>Protein ID: XP_039077163 | MNPEATAFYVEYFYATSQNPDIEETHSHAAYTSVFLPIFYTVVFLTGVLGNLVLMSALHFKRGSRRLIDIFITNLA<br>VSDFIFLVTLPLWVDKEASLGLWRTGSFLCKGSSYMISVNMHCSVFLLTCMSVDRYLAIMCPAVSRKFRRRDCAYG<br>VCVSVWFISCLLGLPTLLSRELTLIDDKPYCAERRATPIKLTWSLVALIFTFFVPLVSIVTCYCCIMRKLYAHYQQ<br>SGKHNKKLRKSIKIIFIVVAAFVFSWLPFNTFKLLAIVSGLQQEVYFSSSFLQLGMEVSGPLAFANSCVNPFIYYI<br>FDSYIRRSILHCVCPCLKNYDFGSSTETSDSHLPKALSNFIQVEDSARRRKRSVSL |
| | *Sus scrofa* | Gene ID:100153787<br>RNA ID: XM_021070686<br>Protein ID: XP_020926345 | METVMDPEATSVYLDYSYITSENPDIEEAPSHLPYTSVFLPIFYTAVFLIGVLGNLILMSALHFKRGSQRLIDIFI<br>INLAASDFIFLITLPVWVDKERSLGLWRTGSFLCKGSSYVISVNMHCSVFLLTCMSVDRYLAIVCPAISRKLRRRD<br>CAYGVCASVWFISCLLGLPTLLSRELTLIDGKPYCAEKKATSIKLTWALVALIFTFFAPLVSMVTCYCCITRKLCV<br>RYQQSGRHNKKLRKSIKIIFIVVAAFVFSWLPFNTFKLLAIVSGLQQELHFSSAFLQWGMEVSGPLAFANSCVNPF<br>IYYIFDSYIRRAIVHCLCPCLKNYDFGSSTETSDLTKALSNFIQGEDFARRRKRSVSL |

| | | | |
|---|---|---|---|
| | *Capra hircus* | Gene ID:102174876<br>RNA ID: XM_005674826<br>Protein ID: XP_005674883 | MDPVMDPEATSVYLEYFFVTSHNPDIEETHSHVPYTSVFLPVFYTAVFLIGVSGNLILMSALHFKRGSRRLIDIFIINLAASDFIFVITLPLWVDKEASLGLWRTGSFLCKGSSYMISVNMHCNVFLLTCMSVDRYLAIVCPAVSRKVRRRDCAYVVCASVWFVSCLLGLPTLLSRELTLIDGKPYCAEERATPIKLTWALVALIFTFFAPLVSIVSCYCCITRKLCVHYQQSGKHNKKLRKSIKIIFIVVAAFVLSWLPFNTFKLLAIVSGLQQELYLSSAFLQRGMEVSGPLAFANSCVNPFIYYIFDGYIRRAIVRCLCPCLKNYDFGSSTETSDLTKALSNFIHAEDFARKRKRSVSL |
| | *Bos taurus* | Gene ID:538740<br>RNA ID: XM_010801022<br>Protein ID: XP_010799324 | MDPVMDPEATTVYLEYLFVTSHNPDIEETHSHVPYTSVFLPVFYTAVFLIGVSGNLILMSALHFKRGSRRLIDIFIINLAASDFIFVITLPLWVDKEASLGLWRTGSFLCKGSSYVISVNMHCNVFLLTCMSVDRYLAIVCPAVSRKVRRRDCAYAVCASVWFVSCLLGLPTLLSRELTLIDGKPYCAEERATPVKLTWALVALIFTFFAPLVSIVSCYCCITRKLCVHYQQSGKHNKKLRKSMKIIFIVVAAFVLSWLPFNTFKLLAIVSGLQQELYLSSAFLQRGMEVCGPLAFANSCVNPFIYYIFDGYIRRAIVRCLCPCLKNYDFGSSTETSDLTKALSNFIHAEDFARKRKRSVSL |
| | *Monodon monoceros* | Gene ID:114891052<br>RNA ID: XM_029215088<br>Protein ID: XP_029070921 | MDPEATTVYLEYFYVTSQNPDIDETHSHVPYTSVFLPVFYTVVFLTGALGNVILMSALHFKRGSRRLIDIFIINLAASDFIFIITLPLWVDKEASLGLWRTGSFLCKGSSYMISVNMHCNVFLLTCMSVDRYLAIMCPAVSRKIRNRDCAYGVCASVWCISCLLGLPTLLSRELTLIDGKPYCAEKRATPVKLAWGLVALIFTFFAPLVSIVTCYCCITRKLCVRYQQSGKHNRKLRKSIKVIFIVVAAFVFSWLPFNTFKLLAIVSGLQQDLYFPSAFLQWGMEVSGPLAFANSGISPFIYYIFDSYIRRAIVHCLCPCLKKYDFGSSTETSDLTKALSNFIQAEDFARRRKRSVSL |
| | *Balaenoptera musculus* | Gene ID:118894400<br>RNA ID: XM_036850649<br>Protein ID: XP_036706544 | MMDPEATTVYLEYFYVTSQNPDIEETHSHVPYTSVFLPVFYTVVFLTGALGNVILMSALHFKRGSRRLIDIFIINLAASDFIFIITLPLWVDKEASLGQWRTGSFLCKGSSYMISVNMHCNVFLLTCMSVDRYLAIMCPAVSRKFRSRDCAYGVCASVWFISCLLGLPTLLSRELTLIDGKPYCAEKRATPTKLAWGLVALICTFFAPLVSIVTCYCCITRKLCVRYKQSGKHNKKLRKSIKVIFIVVAAFVFSWLPFNTFKLLAIVSGLQQELYFPSAFLQWGMEVSGPLAFANGSISPFIYYIFDSYIRRAIVRCLCPCLKTYDFGSSTETSDLTKALSNFIQAEDFARRRKRSVSL |
| | *Choloepus didactylus* | Gene ID:119533383<br>RNA ID: XM_037835461<br>Protein ID: XP_037691389 | MDPEATSVYLDYSEATSQNPEIEETHSHVPYTSVFLPVFYTAVFLTGVLGNLILMGALHFKRGSRRLIDIFIINLAASDFIFLVTLPLWVDKEASLGLWRTGSFLCKGSSYMISVNMHCNVFLLTCMSVDRYLAIVCPTVSRKFRRKDCAYGVCACIWFISFLLGLPTLLSRELTLIDDKPYCAEKIATSSKLTWALVAIIFTFFAPLLSIVTCYCCIIRKLCAHYQKSGKHNKKLRKSIKIIFIVVAAFVISWLPFNTFKLLAIISGLQQEFYFSPAILQRGMEVSGPLAFANSCVNPFIYYIFDSYMRRAIVHCLCPFLKNYDFGSSTDTSDSHLSKALSNFIHAEDFARRRKRSVSL |
| | *Orycteropus afer afer* | Gene ID:103201165<br>RNA ID: XM_007945798<br>Protein ID: XP_007943989 | MDPERTSAYLDYAYATSQIPDEEETPSHVPYTSIILPIFYTVVFLTGVLGNLILMGALHFKQGSRRLIDIFIINLAASDFIFLVTLPLWVDKEASSGLWRTGSFLCKGSSYMISVNMHCSVFLLTCMSVDRYLAIMHPAVSRKFRTRDCAYGVCASIWFISCLLGLPTLLSRELTLIDDKPYCAEKRATVIKLAWALVALIFTFFVPLLSIVTCYCCITRKLCAHYHQSGKHNRKLRKSIKTILIVVTAFTCSWLPFNTFKLLAIVSGLQQEHPVPSAILQLGMEVSGPLAFTNSCVNPFIYYIFDSYIRRAIVYCFCPCLKNYDFGSSTETSDSHLTKAFSTFIHTEDFARRRKRSVSL |
| | *Loxodonta africana* | Gene ID:100672467<br>RNA ID: XM_023559023<br>Protein ID: XP_023414791 | MDPDATSAYLDYSYATSQIPDNEETRSHVPYTSVFLPIFYTVVFLTGVLGNVVLMGALHFKRGSRRLIDIFIINLAASDFIFLVTLPLWVDKEASLGLWRTGSFLCKGSSYMISVNMHCSVFLLTCMSVDRYLAIMYPTVSRKFRRRDCAYGVCAGIWFISCLLGLPTLLSRELILIDDKPYCAEKRSTSVKLAWALVVLIFTFFAPLLSIVTCYCCITRKLCAHYHQSGKHNKKLRKSIKIIFIVVAAFVCSWLPFNIFKLLAIVSGLQQEFYISSAILQLGMEVSGPLAFTNSCVNPFIYYVFDSYIRRAIVHCLCPCLKNYDFGSSTETSDSHLTKALSNFIHTEDFARRRKRSVSL |
| | *Oryctolagus cuniculus* | Gene ID:100339663<br>RNA ID: XM_002716742<br>Protein ID: XP_002716788 | MDPEATPAYLDYYYATDPSPDIQDTGSHIPYLSVFLPIFYTAVFLTGVLGNLVLMGALHFKRGSRRLIDVFIINLAASDFIFLVTLPLWVDKEAAFGLWRTGSFLCKGSSYVISVNMHCSVLLLTCMSIDRYLAILRPAASRKYRRTDCAYGICASVWFISCLLGLPTLLSRELTLIEDKPYCAEKRATPNKLMWALVTLIFTFFVPLLSIVTCYCCITRKLCAHYQQSGKHNKKLKKSIKIIFIVVAAFVLSWLPFNTFKLLAIISGLQQELYFSTATLQQGMEVSGPLAFANSCVNPFIYYIFDSYIRRAILRCLCPCLKNYDFGNSTETSDSHLTKALSNFIHAEDFARKRKRSVSL |
| | *Nycticebus coucang* | Gene ID:128568090<br>RNA ID: XM_053565758<br>Protein ID: XP_053421733 | MDPEAASLYLDYSYATSPNPDLQETRSHVPYTSVFLPVFYTAVFLMGVLGNLVLMGALHFKQGSRRLIDIFIINLAASDFVFLVTLPLWVDKEASLGLWRTGSFLCKGSSYVISVNMYCSVFLLTGMSVDRYLAIVHPAASRKFRRRDCAHGVCASIWFVSGLLGLPTLLSRELRLSGDKLYCAEKKATLVKLMWALVALIFTFFVPLLSIVTCYCCITRKLCAHYQQSGKHNKKLKKSIKIIFIVVAAFVISWLPFNMFKLLAIVSGLQQEPSFSSPVLQLGMEVTGPLAFANSCVNPFIYYVFDSYIRRAIVHCLCPCLKNYDFGSSIETSDSHLTKALSNFIHVEDSTRRRKRSVSL |
| | *Marmota monax* | Gene ID:124100084<br>RNA ID: XM_046455400<br>Protein ID: XP_046311356 | MDPEATSVYLDYYYATSPNSDGKETHSHIPYTSVFLPIFYTAVFLTGVLGNLILIGALYFKRGSRRLIDIFIINLAASDFIFLVTLPLWVDKEASLGLWRTGSFLCKGSSYMISVNMHCSVFLLTCMSVDRYLAIMCPAISRKFRRRDCAYGVCTCVWLVSCLLGLPTVLSRELTLIDGKPYCAEKRATSEKLTWALVALIFTFFVPLLSIVTCYCCITTKLCVHYQQSGKHNKKLRKSIKIIFIVVTAFVFSWLPFNTFKLLAIISGLQPKLHFSSALLQRGMEVSGPLAFANSCVNPLIYYVFDSYIRRAVVRCLCPCLKNYDFGSSTETSDSHLTKALSNLIHTEDFVRRRKRSVSL |
| | *Heterocephalus glaber* | Gene ID:101704263<br>RNA ID: XM_004857951<br>Protein ID: XP_004858008 | MDPETTSVYLDYYYATSSNSDIKETHSSIPYTSVFLSIFYTVVFLTGVLGNLILIGALNFKRGSRRLIDIFIINLAASDFIFLVTLPLWVDKEASLGLWRTGSFLCKGSSYVISVNMHCSVFLLTCMSVDRYLAIMCPAMSRKFRRRDCAYGVCASIWFISCLLGLPTLLSRELTLIENKPYCAEKKATSVKLTWGLVTLIFTFFVPLLSIVTSYSCITRKLCAHYQQSGKHNKKLRKSMKIIFIVVAAFVFSWLPFNTFKLLAIVSRLKHEHHFSSTTLQLGIEVSGPLAFANSCVNPLIYYTFDSYIRRAIVHCLCPCLKNYDFGSSTETSDSHLTKALSNFIHAEDFVRRRKRSVSL |
| | *Chinchilla lanigera* | Gene ID:102015745<br>RNA ID: XM_005386275<br>Protein ID: XP_005386332 | MDLETTSVYLDYYYAASPNSHIKETSSQVPYTSVFLSIFYTAVFLTGVLGNLILIGALHFKRGSRRLIDIFIINLAVSDFIFLVTLPLWVDKEASLGLWRTGSFLCKSSSYMISVNMHCSVFLLTCMSVDRYLAIMYPAVSRKFRRRDCAYGVCASVWFISCLLGLPTLLSRELTLIEDKPYCAEKRATSMKLTWALVTLIFTFFVPLLSIVTCYCCITRKLCAHYQQSGKHNKKLRKSIKIIFIVVAAFVFSWLPFNTFKLLAIVSGLQNGLRFSSAALQLGMKVSGPLAFANSCVNPLIYYIFDSYIRRAIMHSLCPCLKSSDFGSNTETSDSHLSKALSNFIHADDFVRRRKRSVSL |
| | *Cavia porcellus* | Gene ID:100720613<br>RNA ID: XM_003469238<br>Protein ID: XP_003469286 | MDLKTTSDYLDYYYATSPNSHIKGSSQVPYTPVFLSVFYTAVFLTGVLGNLILIGALHCKRGSRRLIDIFIINLAASDFIFLVTLPLWVDKEASLGLWRTGSFLCKGSSYMISVNMHCSVFLLTCMSVDRYLAIMCPAVSRKFRRRDCAYRVCASIWFISCLLGLPTLLSRELTLIEDKSYCAEKRATSMRLTWALVTLIFTFFIPLLSIVICYCCITRKLYTHYQQSGKHNKKLRKSIKIIFIVVAAFVFSWLPFNTFKLLAIVSGLQNELYFSSAALQLGMKVSGPLAFANSCVNPLIYYTFDSYIHRAIVHCLCPCLRNSDIGSSTETSDSHLAKALSNFVHADDFVRRRKRSVSF |

| | | | |
|---|---|---|---|
| | *Tupaia chinensis* | Gene ID:102482783<br>RNA ID: XM_006163920<br>Protein ID: XP_006163982 | MDPEATLVYLDYYTASPDPDIKETHPHAPYTSVFLPIFYTAVFLTGVLGNLVLMGALYFKQGTRRLIDIFIINLAASDFIFLVTLPLWVDKEASLGLWRTGAFLCKGSSYMISVNMHCSVFLLTCMSVDRYLAIMCPAIARQVRRRNCAYGVCACVWFISCLLGLPTLLSRELTLIDDKPYCAEKKASFIKLAWALVALIVTFFVPLLSIVTCYCCITKKLCTHYQQSGKHNKKLKKSIKIISIVVAAFVVCWLPFNSFKLLAIVSGLQQELYFSSAILRLGMEVSGPLAFTNSCVNPFIYFIFDGYIRRAIIHCLCPCLKKYDFGSSTETSDSHLTKALSNCIHAEGDTRRRKRSVSL |
| | *Rattus norvegicus* | Gene ID:288181<br>RNA ID: NM_001105890<br>Protein ID: NP_001099360 | MEPATTLFYLDYYDATSPDPRIMETPSHTSYTSVFLPVFYTAVFLTGVLGNFILMVTLHFKHRNRRLIDIFIINLAASDFIFLVTLPLWVDKEASLGLWRTGSLLCKGSSYTISVNMHCNVFLLTCMSMDRYLAIMRPTLARRLRRRSCAYAVCAVIWIISCLLGLPTLLSRGLTHIEGKPYCAEKKPTSLKLMWGLVALIATFFVPLLSIVSSYCCITRRLCAHYQQSGKHNKKLRKSIKIIVIVVAAFTISWVPFNTVKLLAIVSGLQPESQFPSESLQQAMKVTGSLAFANSCVNPLLYYIFDSYIRRAIVRSLCPCLKIHNIGSSTETLDSHLTKALANFIHSEDFVKRRKRSVSL |
| | *Mus musculus* | Gene ID:71223<br>RNA ID: NM_001162955<br>Protein ID: NP_001156427 | MEPATALLIVDYYDYTSPDPPFLETPSHLSYTSVFLPIFYTVVFLTGVVGNFILMIALHFKRGNRRLIDIFIINLAASDFIFLVTVPLWMDKEASLGLWRTGSFLCKGSSYVISVNMHCSVFLLTCMSMDRYLAIMHPALAKRLRRRSSAYAVCAVVWIISCVLGLPTLLSRELTHIEGKPYCAEKKPTSLKLMWGLVALITTFFVPLLSIVTCYCCITRRLCAHYQQSGKHNKKLKKSIKIVIIAVAAFTVSWVPFNTFKLLAIVSGFQPEGLFHSEALQLAMNVTGPLAFASSCVNPLIYYVFDSYIRRAIVRCLCPCLKTHNFGSSTETSDSHLTKALSNFIHAEDFIRRRKRSVSL |
| | *Sarcophilus harrisii* | Gene ID:100915939<br>RNA ID: XM_003766363<br>Protein ID:XP_003766411 | MEETTYLTYLDNYASSQTPDFEDDHLHVLYTSIFLPIFYTAVFIVGVVGNLILIGALHFKRGSQRLIDIFIINLAISDFIFLVTLPLWVDKEASLGIWRTGSFLCKGSSYIISVNMHSNVFFLTCMSAERYLAIMCPSASRKFRRKDCAYGICVSVWFISCLLGLPTLLSRDLTIIEDKPYCAEEPATLSKYAWALVCLIFTFFVPLLSILTCYCSIARKLCAYYQQSRKHNKKLQKSIKIIFIVVAAFVSFWLPFNIFKVLSIISGLQEEPFFSSTILQVGMEVSGPLAFSNSCINPFIYYYFDGYIRRAIICCLCPCLKNSTFGSSTETSDSHLNKIFGNFIHGENFSRRRRRSVSL |
| | *Vombatus ursinus* | Gene ID:114022936<br>RNA ID: XM_027835378<br>Protein ID:XP_027691179 | MDETTSLTYLDNYATSQGPDFEDDHSHVLYTSVFLPIFYTAVFLIGVVGNLILIGALHFKRGSQRLIDIFIINLAVSDFIFLVTLPLWVDKEASLGLWRTGSFLCKGSSYIISVNMHSSVFFLTCMSAERYLAIMCPSASRKFRRKDCTYGVCVSVWFISCLLGLPTLLSRDLTMIEDKPYCAEEPATLSKHAWALVSLIFTFFVPLLSILTCYCSIARKLCTYYQQSGKHNKKLQKSIKIIFIVVAAFVSSWLPFNIFKVLSIISGLQEEPFFSSAMLQVGMDISGPLAFSNSCINPFIYYFFDGYIRRAIICCLCPCLKNSNLGSSTETSDSHLSKIFTNFIHGEDFSRRRRRSVSL |
| | *Phascolarctos cinereus* | Gene ID:110202008<br>RNA ID: XM_020977958<br>Protein ID:XP_020833617 | MDETTPLTYLDIYATSQGLDFEDDHLYVLYTSVFLPIFYTAVFLIGVVGNLILIGALHFKQGSQRLIDIFIINLAVSDFIFLVTLPLWVDKEASLGLWRTGSFLCKGSSYIISVNMHSSVFFLTCMSAERYLAIMCPSASRKFRRKDCTYGVCVSVWCISCLLGLPTLLSRDLIMIEDKPYCAEEPATLSKHAWALMSLIFTFFVPLLSILTCYCSIARKLCAYYQQSGKHNKKLQKSIKIIFIVVAAFVSSWLPFNIFKVLSIISGLQEEPFFSSVTLQVGMEVSGPLAFSNSCINPLIYYFFDGYIRRAIICCLCPCLKNSNLGSSTETSDSHLSKIFANFIHGEDFSRRRRRSVSL |
| | *Gracilinanus agilis* | Gene ID:123239666<br>RNA ID: XM_044666949<br>Protein ID:XP_044522884 | MDETTPLSYMDNYATSQDLDFEDDHSHVLYTSIILPIFYTVVFLIGVVGNLILMGALHFKRGSQRLIDIFIINLAVSDFIFLVTLPLWVDKEASLGLWRTGSFLCKGSSYIISVNMHSSVFLLTCMSAERYLAIMCPSASRKFRRKNCTYGVCVIVWLVSCLLGLPTVLSRKLTMIEERPYCAEEPATLSKRAWALVSLIFTFFVPLLSILTCYCSIARKLCAHYQRSGKHNKKLRKSIKIIFIVVVAFVSSWLPFNIFKLLSIISGLQEEPFLSSATLQVGMEVSGPLAFSNSCINPFIYYFFDGYIRRAIICCLCPCPKNYNLGSSTETSDSHLSKIFANFTHGEDSSRKRRRSVSL |
| | *Dromiciops gliroides* | Gene ID:122751380<br>RNA ID: XM_043998389<br>Protein ID:XP_043854324 | MDETTPLYYVDNFATSQGPEFEDDHLHVLYTSVFLPIFYTVVFLIGVVGNLILIGALHFKRGSQRLIDIFIINLAVSDFIFLVTLPLWVDKEASLGLWRTGWFLCKGSSYIISVNMHSSVFLLTCMSAERYLAIMCPSASRKFRRKDCTYGICVSVWFISCLLGLPTLLSRDLTMIEDKPYCAEEPATLSKRAWALVSLIFTFFVPLLSILTCYCSIARKLCAHYQQSGKHNKKLRKSIKIIFIVVAAFVSSWLPFNIFKLLSIISGLQEEPFFSSAILQVGMEVSGPLAFSNSCINPFIYYFFDGYIRRAIICCLCPCLKNYNLGSSTETSDSHLSKVFANFLHGEDFSRRRRSVSL |
| Birds | *Passer domesticus* | Gene ID:135295574<br>RNA ID: XM_064411336<br>Protein ID:XP_064267406 | MRTAGPETELAPLSSMTTVTFNYDYYYEDQCQYQHLQHMSTFLPILYSAVFLVGIIGNSILIAALIFKRRVQRLIDIFIINLATSDFIFLITLPFWVDMEVSDETWRVGSFLCKASSYVISVNMYCSILLLTCMSADRYLAIMHPCVARRIRTRSYFGGLCISVWVLSGCLGIPTLLSRELKTQYGKTYCTDKAVTEVKQIMSLMLLILAFFVPLLSILTFYCSITKRLCVHYQKAGKRDKKLRKSIKIVFIVVAAFVVSWVPYNLFKLMAILLRLLKQPSCFSGTVAQLGMKVSSPFAFANSCANPFIYFCFDSYIRRAMLQCLCPRVKMSSNSSDTLDTHLSHSLSSFVAGEYSTRKRKRSVSL |
| | *Cyanistes caeruleus* | Gene ID:111939404<br>RNA ID: XM_023941618<br>Protein ID:XP_023797386 | MRTAGSEMEFPELSSTAAVTFNYDYYYDDQCQHQHLQHMSVFLPVLYSAVFLVGIVGNSILIVALVFKRQVQRLIDIFIVNLAASDFIFLITLPFWVDMEVSDESWRVGSFLCKASSYVISVNMYCSILLLTCMSADRYLAIMHPSIARQIRTRAYSRALCICVWLVSCCLGMPTLLSRELRKQDGKTYCTDKAVTEVKLIMSLMLLILAFFFPLLSILTFYCSITKRLCVHYQKAGKRDKKLRKSIKIVFIVVAAFVVSWVPYNLFKLMAILLRLLKQPDCFTGTVAQLGIKVSSPFAFANSCANPFIYFCFDNYIRRAMLQCLCPWVKISSNSSETLDTNLSHSLSNFVAGEFATRKRKRSVSL |
| | *Cinclus cinclus* | Gene ID:134056051<br>RNA ID: XM_062512552<br>Protein ID:XP_062368536 | MRTTGPEMDFPLSSMATVTFNYDYYYNDQCQYQHLQHMSTFLPILYSAVFLVGIVGNSILIAALVFKRRVQRLIDIFIINLAASDFIFLITLPFWVDMEVSDESWRVGSFLCKASSYVISVNMYCSILLLTCMSADRYLAILHPSVARRIRTRSYSRALCICVWLLSCCLGMPTLLSRELKKEYGKTYCTDKAVTEVKQIMSLMLLILAFFFPLLSILTFYCSITKRLCVHYQKGGKCDKKLRKSIKIVFIVVVAFVVSWVPYNLFKLMAILLRLLKQPDCFSGRVAQLGIKVSSPFAFANSCANPFIYFCFDNYIRRAMLQCLCPQVKISSNSSDTLDTHLSHSLSNFVAGEYATRKRKCSVSL |
| | *Chamaea fasciata* | Gene ID:136284823<br>RNA ID: XM_066175247<br>Protein ID:XP_066031344 | MRTAGPEMELPLLSSVATGTFSYDYYYYDDQCQYQHLQHMSTFLPILYSAVFLVGIVGNSVLIAALVFKRRVQRLIDIFIVNLAASDFIFLITLPFWVDMEVSDESWRVGSFLCKASSYVISVNMYCSILLLTCMSADRYVAIMHPSLARRIRTRSYSRALCICVWLLSCCLGMPTLLSRELKKQYGKTYCTDKDVTEAKQIMSLMLLILAFFFPLLSILTFYCSITKRLCVHYQKTGKHDKKLRKSIKIVFIVVAAFVVSWVPFNLFKLMAILLRLLKQPDCFSGTVAHLGIKVSSPFAFANSCANPFIYFYFDNYIRRAMLQCLCPRVKISSNSSDTLDTHLSHSLSNFVAGEYATRRRKHSVSL |
| | *Struthio camelus* | Gene ID:104144263<br>RNA ID: XM_009675189<br>Protein ID:XP_009673484 | MELTPLSPVTAVTFNYEYYYEDTCQYDHLQHMSTFLPVLYTAVFLVGIVGNSILIVALVFKRRIQRLIDVFIINLAASDFIFLITLPFWVDKEASDGSWRVGSFLCKASSYIISVNMHCSILLLTCMSADRYLAIMYPSTARRVRTRSYSSGLCICVWLLSCCLGMPTLLSRELKEHYGKTYCTDKAMTETKQIASLMILILAFFFPLLSILTFYCSITRKLCVHYQKSGKHDKKLRKSIKIVFIVVAAFVISWVPFNLFKLTAIVLGLLKPPECFPDTVAQLGMKVSSPFAFANSCANPFIYYCFDNYIRRAMLRCLCPWVKISSSGNNSDTLDTRLSHSISNFVAGEYAARKRKRSVSL |

| | | | |
|---|---|---|---|
| | *Dromaius novaehollandiae* | Gene ID:112979637<br>RNA ID: XM_026093988<br>Protein ID:XP_025949773 | MRTAWPEMELTQLSPVTAVTFNYDYYEDNCQYHHLQHMSTFLPVLYTAVFLVGIIGNSILIVALVFKRRIQRLIDV<br>FIINLAASDFIFLITLPFWVDKEASDGSWRVGSFLCKASSYIISVNMYCSILLLTCMSADRYLAIMYPSTARRVRT<br>RSYASGLCICVWLLSCCLGMPTLLSRELKEHYGKMYCTDKEMTEPKQIASLMILILAFFLPLLSILTFYCSITRKL<br>CMHYQKSGKHDKKLRKSIKIVFIVVAAFVISWVPFNLFKLIAVLLGLLKPPDCFPDLVAQLGMKVSSPFAFANSCA<br>NPFIYYCFDNYIRRAMLRCLCPWVKISNSSNNSDTLDTRLSHSLSNFVAGEYATRKRKRSVSL |
| | *Apteryx rowi* | Gene ID:112974029<br>RNA ID: XM_026082543<br>Protein ID:XP_025938328 | MRTAWPEMELTQLSPVTAVTFNYDYYEDNCQYHHLQHMSTFLPVLYTVVFLVGIIGNSILIVALVCKRRIQRLIDI<br>FIINLAASDFIFLITLPFWVDKEASDGSWRVGSFLCKASSYIISVNMYCSILLLTCMSADRYLAIMYPSTARRVRT<br>RSYASGLCISVWLLSCCLGMPTLLSRELKEHYGKTYCTDKAMTEAKQIASLMILILAFFFPLLSILTFYCSITRKL<br>CVHYQKSGKHDKKLRKSIKIVFIVVAAFVISWVPFNLFKLMAILWGLLKPPDCFPDLVAQLGMKVSSPFAFANSCA<br>NPFIYYCFDNYIRRAMIRCLCPWVKLSSSSNNSDTLDTRLSHSLSNFVVGEYATRKRKRSVSL |
| | *Indicator indicator* | Gene ID:128970543<br>RNA ID: XM_054385775<br>Protein ID:XP_054241750 | MRTDQTEMELTQLSPVTTVTFNYDYYDDDNCQYQHLQHMSTFLSILYTAVFLVGTVGNGILILALIFRRQVQRLID<br>IFIINLAASDFIFLITLPFWVDMEASDGSWRVGSFLCKASSYVISVNMYCSILLLTCMSADRYLAIMHPSIARQVR<br>TRSYSSRLCICVWFLSCCLGMPTLLSRELKKQYGKTYCTDKAVTEAKQIMSLMLLILAFFFPLLSILTFYCTITRR<br>LCVHYQRAGKHGKKLRKSIKIVFIVVAAFVISWVPFNLFKLMAIILGLLKQPDCFPNMVAQLGMKVSSPFAFANSC<br>ANPFIYYCFDSCIRRAMLRWLCPWVKISSSSTNCDTLDTRLSHSLSLSPAGEHAARKRKRSVSL |
| | *Nestor notabilis* | Gene ID:104404943<br>RNA ID: XM_010013496<br>Protein ID:XP_010011798 | MGTAWPEMELTQLSSMTTVTFNYDYYYYDDNCQYQLLQHMPTFLSILYTAMFLVGIIGNSILIAALVFKRRVQRLI<br>DIFIINLAASDFIFLITLPFWVDKEASDGSWRVGAFLCKASSYVISVNMYCSILLLTCMSADRYLAIMHPSIARRV<br>RTRSYCSGLCICVWLLSCCLGMPTLLSRELKKHYGKTYCTDKAVTEDKQIMSLMLLILAFFFPLLSILTFYCSITK<br>RLCVHYQRTGKHDKKLRKSIKIVSIVVAAFIISWVPFNLFKLMAILLGLLKQPNCFPDVVAQLGMKVSSPFAFANS<br>CANPFIYYCFDNYIRRAMLRCLCPWVKSSSSGNNSDTLDTRLSYSLSNFIAGENAARKRKRSVSL |
| | *Apus apus* | Gene ID:127389321<br>RNA ID: XM_051629705<br>Protein ID:XP_051485665 | MRTACPEMELTQFSPVTTVTFTYDYYYDDTCQYQHLQHMSTFLPILYTAVFLVGIVGNSILIVALVFKRRVQRVID<br>IFIINLAASDFIFLITLPFWVDKEASDGSWRVGSFLCKASSYVISVNMYCSILLLTCMSADRYLAIMHPSIARRVR<br>TRSYSSGLCICVWLISCCLGMPTLLSRELKNQYGKTYCTDKAVTEAKQIMSLMLLILAFFFPLLSILTFYCSITRR<br>LCVHYQRAGKYDKKLRKSIKIVFIVVAVFVTSWVPFNLFKLMAILLGLLKQPDCYPNRIAQLGMMVSSPFAFANSC<br>ANPFIYYCFDNYIRRAMLQCLCPRVKISSTGNNSDTLDSRLSHSLSTFVAGEYPARKRKRSVSL |
| | *Athene cunicularia* | Gene ID:113481790<br>RNA ID: XM_026851714<br>Protein ID:XP_026707515 | MRTAGPEMELTELSPVPTVTFNYEYYYDDNCQYQHLQHMATFLPVLYTAVFLVGIIGNSILIVALVFKQRVQRLID<br>IFIINLAASDFIFLITLPFWVDMEASDGSWRVGSFLCKASSYVISVNMYCSILLLTCMSADRYLAIMYPSIARRVR<br>TRSYSSGLCICVWLLSCCLGMPTLLSRELKKQYGKTYCTDKAMTEAKQIMSLMLLILAFFFPLLSILTFYCSITRR<br>LCLHYQRAGKHDKKLRKSIKIVFIVVVAFVISWVPFNLFKLMAVLLGLLKQPDCFPNMVAHLGMKVSSPFAFANSC<br>ANPFIYYCFDNYIRRAMLRCLWPRVKIIISSSNNSDTLDTHLSHSLSNFVVGEYAARKRKRSLSL |
| | *Grus americana* | Gene ID:129207004<br>RNA ID: XM_054827285<br>Protein ID:XP_054683260 | MRIAWPEMELTQLSPVTTVTFNYDYYYDDNCQYQHLQHMSTFLPILYAAMFLVGIVGNSILIVALVFKRRVQRLID<br>VFIINLAASDFIFLITLPFWVDKEVSDGSWRVGSFLCKASSYVISVNMYCSILLLTCMSADRYLAIMHPSLARRVR<br>TRSYSSGLCICVWLLSCCLGMPTLLSRELKKQYGKTYCTDKVVTETKQIMSLMLLILAFFFPLLSILTFYCSITRR<br>LCVHYQRAGKHDKKMRKSIKIVFIVVAAFVISWVPFNLFKLMAILLGLLKQSDCFPDMVAQLGMKVSSPFAFANSC<br>ANPFIYYCFDNYIRRAMLRCLCPWVKISSSSNNSDTLDTRLSHSLSNFVVGEYAARKRKRSVSL |
| | *Harpia harpyja* | Gene ID:128144909<br>RNA ID: XM_052794325<br>Protein ID:XP_052650285 | MRTACPEMELTQLSPVTTVTFNYDYYYDDNCPYQHLQHMSTFLPILYTAVFLVGIAGNSILIVALVFKQRVQRLID<br>VFIINLAASDFIFLTTLPFWVDKEVSDESWRVGSFLCKASSYVISVNMYCSILLLTCMSADRYLAIMHPSIARRVR<br>TRSYSSGLCICVWLLSCCLGMPTLLSRELKKQYGKTYCTDKAVTETKQIVSLMLLILAFFFPLLSILTFYYSITRR<br>LCVHYQRAGKHDKKLRKSIKIVFIVVAAFVISWVPFNIFKLMAILLGLLKQPDCFPNMVAQLGMKVSSPFAFANSC<br>ANPFIYYCFDNYIRRAMLRCLCPRVKISSSGNNSDTLDTRLSHSLSNFAAGEYATRKRKRSVSL |
| | *Aptenodytes forsteri* | Gene ID:103907485<br>RNA ID: XM_009289680<br>Protein ID:XP_009287955 | MPEMELTQLSPVTTVTFNYDYYYDDNCQYQHLQHMSTFLPILYTAVFLVGIVGNSVLIVALVFKRRVQRLIDVFII<br>NLAASDFIFLITLPFWVDKEASDGSWRVGPFLCKASSYVISVNMYCSILLLTCMSADRYLAIMHPSIARRVRTRSY<br>SSGLCICVWLLSCCLGMPTLLSRELKKQYGKTYCTDKAVTEAKQIVSLTLLILAFFFPLLSILTFYCSITKRLCVH<br>YQRAGKHDKKLRKSIKIVFIVVAAFVISWVPFNLFKLMAILLGLLKQPDCFPDMVAQLGMKVSSPFAFANSCANPF<br>IYYCFDNYIRRAMLRCLCPRVKISSSGNNSDTLDTHLSHSLSNFVAGEYAARKRKRSVSL |
| | *Meleagris gallopavo* | Gene ID:100550729<br>RNA ID: XM_010719502<br>Protein ID:XP_010717804 | MRTAGPEMDLIQLSSVTTVTFNYDDYYYEDSCQYEQLQHMTAFLPILYAAVFLVGILGNSILIIALVFKRRVQRLI<br>DIFIINLAASDFIFLITLPFWVDKEASDGIWRVGSFLCKASSYIISVNMYCSILLLTCMSADRYLAIMYPSIARRV<br>RTRSYSSGLCLCVWLLSCCLGMPTLLSRELNERYGKMYCTDKDVTKSKQITSLMILILAFFFPLLSILTFYCSITK<br>RLCVHYQRSGKHDKKLRKSIKIVFIVVAAFVISWVPFNLFKLMAILLGLQKLPDCFLDMVAQVGMTVSSPFAFANS<br>CANPFIYYCFDNYIRRAMLRCLCPWVKASSGSTVSDTLDTRLSHSLSNFIAGEYTARKRKRSVSL |
| | *Gallus gallus* | Gene ID:427956<br>RNA ID: XM_004938212<br>Protein ID:XP_004938269 | MRTAGPEMDLIKLSPMTTVTFNYDDYYEDSCQYEQLQHMTAFLPVLYAAVFLVGILGNSILIVALVFKRRVQRLID<br>IFIINLAASDFIFLITLPFWVDKEASDGIWRVGSFLCKASSYIISVNMYCSILLLTCMSADRYLAIMYPSIARKVR<br>TRSYSSGLCFCVWLLSCCLGMPTLLSRELTERYGKMYCTDKAMTESKQITSLMILILAFFFPLLSILTFYCSITRR<br>LCVHYQRSGKHDKKLRKSIKIVFIVVAAFVISWVPFNLFKLMAILLGLQKLPDCFLGMVAQVGMTVSSPFAFANSC<br>ANPFIYYCFDNYIRRAMLRCLCPWVKASSGSTISDTMDTRLSHSLSNFITGDYTARKRKRSVSL |
| | *Coturnix japonica* | Gene ID:107320109<br>RNA ID: XM_015875626<br>Protein ID:XP_015731112 | MRTAGPEMDLIQLSPVTTVTFSYDDYYYEDEDSCQHKQLQHMTAFLPILYAAVFLVGILGNSILIVALVFKRRVQR<br>LIDVFIINLAASDFIFLITLPFWVDKEASDGMWRVGSFLCKASSYIISVNMYCSILLLTCMSADRYLAIMYPSIAR<br>RVRTRSYSSGLCFCVWLLSCCLGTPTLLSRELEERYGKMYCTDKDVTESKQITSLVILILAFFFPLLSILTFYCSI<br>TRRLCMHYQRSGKHDKKLRKSIKIVFIVVAAFVISWVPFNLFKLMAILLRLQKLHDCFLDMVAQLGIRVSSPFAFA<br>NSCANPFIYYCFDNYIRRAMLRCLCPWVKASNSTTISDTLDTRLSYSLSNFITGEYTARKRRRSVSL |
| | *Anser cygnoides* | Gene ID:106038001<br>RNA ID: XM_013184209<br>Protein ID:XP_013039663 | MRTAWPEMDLSQLSPVTMVTLNYDDYFYEDNCQYKRLQHMTTFLPILYTVVFLVGIIGNSILIAALVFKRRVQRLI<br>DIFIINLAASDFIFLITLPLWVDKEVSDGSWRVGSFLCKASSFIISVNMYCSILLLTCMSADRYLAIMYPAVARRV<br>RTRSYSTGLCICVWLLSCCLGIPTLLSRELQERYGKTYCADKAVTESKQIVSLMMLILAFFFPLLSILTFYCSITR<br>RLCMHYQRSGKHDKKLRKSIKIVFIVVAAFVISWVPFNLFKLMAILLGLRKLPDCFPDVVAQVGMKVSSPFAFANS<br>CANPFIYYCFDNYIRRAMLRCLCPWVKVSSSGSVSDTLDTRLSHSLSNFIVGEYAARKRKRSVSL |

| | | | |
|---|---|---|---|
| | *Anas acuta* | Gene ID:137863669<br>RNA ID: XM_068697030<br>Protein ID:XP_068553131 | MRTAWPEMDLSQLSPVTMVTLNYDDYFYEDNCQYKHLQHMTTFLPILYTVVFLVGIVGNSILIAALVFKRRVQRLIDIFIINLAASDFIFLITLPFWVDKEVSDGSWRVGSFLCKASSFIISVNMYCSILLLTCMSADRYLAIMYPAVARRIRTRSYSTGLCICVWLLSCCLGIPTLLSRELKERYGKTYCADKAVTESKQIVSLMMLILAFFFPLLSILTFYCSITRRLCMHYQRSGKHDKKLRKSIKIVFIVVAAFVISWVPFNLFKLMAILLGLRKLPDCFPDVVAEVGMQVSSPFAFANSCANPFIYYCFDNYIRRAMLRCLCPWVKVSSSGSVSDTLDTRLSHSLSNFIAGEYAARKRKRSVSL |
| Reptiles | *Pseudonaja textilis* | Gene ID:113441761<br>RNA ID: XM_026708926<br>Protein ID:XP_026564711 | MEETTPSYDYFYSFSSDTPEENCQTLKLPYMEIFVSVLYIVIFLVGTVGNGLLIGVLIFKQYVWRLTDTFIVNLAISDFSFLITLPFWIDKELASGLWRSGSILCKGSSYVISVNMYCSIFLLTWMSGDRYLTIMYPSMAKKIRTKLYPILVCISVWILSCLLGLHTLQSRELKRYNNHTYCVDKETTFNNWVGSLMLLTLAFFIPLFSILTLHYSIIKKLYEHYQKFGKHDKKLRKSIKIVFMVAIVFFFSWTPFNIFKILALMSSILELKQSFCLYYKIVYLGMELGGLFAFANSCTNPFIYFFFDDCIHRAMIQSILPCRKYNKPGSSFSSLDT |
| | *Notechis scutatus* | Gene ID:113421410<br>RNA ID: XM_026681780<br>Protein ID:XP_026537565 | MEETTPSYDYFYSFSSDTPEENCQTLKLPYMEIFVSVLYIVIFLVGTVGNGLLIGVLIFKQNVWRLTDTFIVNLAISDFSFLITLPFWIDKELASGLWRSGSILCKGSSYVISVNMYCSIFLLTWMSGDRYLTIMYPSMAKKIRTKLYPILVCISVWILSCLLGLHTLQSRELKRYNNHTYCVDKETTFNNWVGSLMLLTLAFFIPLFSILTLHYSIIKKLYEHYQKFGKHDKKLRKSIKIVFMVAIVFFFSWTPFNIFKILALMSSILELKQSFCLYYKIVYLGMELGGLFAFANSCTNPFIYFFFDDCIHRAMIQSILPCRKYNKPGSSFSSLDT |
| | *Erythrolamprus reginae* | Gene ID:139167269<br>RNA ID: XM_070751717<br>Protein ID:XP_070607818 | MEETTPIYDYFYSLNSDPLEEDCETLKLPYMEIFVSVLYIVIFLIGSVGNGLLIGALIFKQCVWRLTDTFIVNLAISDFSFLITLPFWIDKELASGLWRSGSILCKGSSYIISVNMYCSIFLLTWMSGDRYLTIMYPSMAKKIRTKLYPILICTSIWILSCLLGLPTLQSRELGRYNNNTYCMDKETITNNWIASLMLLTLAFFIPLFSILTLHYSIIKKLYMHYQKFGKQDKKLRKSIKIVFVVAIVFSLSWTPFNIFKILALVSSILEPKEPKESFCVYKKMVYLGMELGGLFAFANSCANPFIYFFFDDCIHRAMIQTILPCRKYNKPGSSFSSLDT |
| | *Crotalus tigris* | Gene ID:120300815<br>RNA ID: XM_039327326<br>Protein ID:XP_039183260 | MEETTPSYDYFYYLSPDAPEENCQILKLPNMEILVSVLYIVIFLVGIVGNGLLIGVLISKQRVWRLTDTFIVNLAISDFSFLITLPFWIDKELASGLWRSGFILCKGSSYIISVNMYCSIFLLTWMSGDRYLTIMYPSMARKIRTKLYPILVCICVWILSCLLGLPTLQSRELRRYNNNTYCVDKETIFNNWIGSLILLILAFFIPLFSILTLHYSIIKKLYVHYQKFGKHDKKLRKSIKIVFIVAVVFLFSWTPFNIFKILALISNNSELQQSFCLYYKIAHLGMELGGLFAFANSCTNPFIYFFFDNCIRRTIIQSILPCRKYNRPGSSFSSLDT |
| | *Python bivittatus* | Gene ID:103065587<br>RNA ID: XM_007435872<br>Protein ID:XP_007435934 | MEETTPSYDYFYSLSPDAPEENCQVLKLPYMETFVSVLYIAIFLVGTASNGLLIGVLIFKQRVWRLTDTFIVNLAISDFSFLITLPFWVDKELASGLWRSGSVLCKGSSYIISVNMYCSIFLLTWMSGDRYLTIMYPSVARKIRTKLYPIFVCISVWILSCLLGLPTLQSRELRRYNNNTYCVDKETISSNWIGSLLLLILAFFIPLFSILILNYFIIKKLYVHYQKFGKHDKKLRKSIKIVFTITVVFLFSWIPFNIFKILALISSTQELKQPFCLHYKIAYLGMELGGLLAFTNSCTNPFIYYFFDDCIRRATMQSIIPCRKTNRPGSSFASLDTCLRFSES |
| | *Candoia aspera* | Gene ID:134499109<br>RNA ID: XM_063305665<br>Protein ID:XP_063161735 | MEETTLSYDYFYSLSPDAPEEDCQTLKLPYMEIFVSVLYIAIFLVGTAGNGLLIGILIFKQHVWRLTDTFIVNLAISDFSFLITLPFWVDKELASGLWRSGSVLCKGSSYIISVNMYCSIFLLTWMSGDRYLTIMHPSMARKIRTKHYPIFVCISVWIVSCLLGLPTLQSRELRRYNNNTYCVDKETLYNNWIGSLLLLILAFFIPLFSILTLHYSIIKKLYVHYQKFGKHDKKLRKSIKIVFMVAVVFLFSWTPFNIFKILALTSSIQELKQPFCLHYKIAYLGMELGGLLAFTNSCTNPFIYYFFDDCIRRAMMQSILPCRKAKRPGSSFASLDTCLRSSESSFLPENVVLRKRRKSV |
| | *Tiliqua scincoides* | Gene ID:136645750<br>RNA ID: XM_066622144<br>Protein ID:XP_066478241 | MEGPNFSNYDYYDYNYSSPEPPEDCPALQVPYKYIFLPAVYGTVFLLGVVGNAVLMGALIFKRRIWRLVDIFIINLAASDFIFLITLPFWVDKEMYSGLWRSGSFFCKGSSYVISVNMFCSIFLLTCLCIDRYLAIMYPSMARKIRTKLYSIILCVCVWILSCLLGLPTLLSRELRSFEDGNMYCMEKDPTFTNRIVALLVLILAFFVPLFIILMFYCSITKKLCVHYHKSGKHDKKLRKSIKIVFIVVIAFVFSWTPFNIFKLLALISGIQELKPPFCLPFKVAHLGMELSAPFAFANSCVNPFIYYFFDDYIYRTVLKLSCMKTNSLGSSSDSSLSRLSYFMNTFEHGEDVSRRRRRSVSF |
| | *Zootoca vivipara* | Gene ID:118085314<br>RNA ID: XM_035115845<br>Protein ID:XP_034971736 | MEKSTSSYDYYFYTDLPEDNCSAVELPYKHIFLPALYTIVFLVGIAGNALLIGALIFKRRIQRLIDIFIVNLAVSDFIFLITLPFWVDKERTSGLWRSGSFLCKGSSYTISVNMYCTIFLLTCMSSDRYLAIMYPSVARRIRTKLYSIILCTCVWILSCLLGLPTLLSRELRRYDNSTYCVDKEITSTHQIGSVLLLILAFFVPLFIILTFYCSITKKLCVHYQKSGKHDKKLKKSIKIVFTVVIIFVFSWAPFNIFKFLALMSAIQELKPPFCLPYKVAHIGMELSGPFAFANSCTNPFIYYFFDDYIRRAMMQCITPCTKANNFGTSSESSDTRLSYSLTAFAHREDVSRKRRRSVSF |
| | *Podarcis raffonei* | Gene ID:128412768<br>RNA ID: XM_053386102<br>Protein ID:XP_053242077 | MEKSTSSYDYYFYTDPPEDNCSAVELPYKHIFLPALYTIVFLVGIAGNALLIGALIFKRRVQRLIDIFIVNLAVSDFIFLITLPFWVDKERTSGLWRSGSFLCKGSSYTISVNMYCTVFLLTCMSSDRYLAIMYPSVARRIRTKLYSIILCTCVWILSCLLGLPTLLSRELRRYDNSTYCVDKEITSTHRIGSLLLLILAFFVPLFIILTFYCSITKKLCVHYQKSGKHDKKLKKSIKIVFTVVIIFVLSWAPFNIFKFLALMSAIQELKPPFCLPYKVAHIGMELSGPFAFANSCTNPFIYYFFDDYIRRAVMQCITPCTKANNFGTSSDSSDTHLSYSLTAFAHREDVSRKRRRSVSF |
| | *Podarcis muralis* | Gene ID:114595834<br>RNA ID: XM_028726566<br>Protein ID:XP_028582399 | MEKSTSSYDYYFYTDSPEDNCSAVELPYKHIFLPALYTIVFLVGIAGNALLIGALIFKRQVQRLIDIFIVNLAVSDFIFLITLPFWVDKERTSGLWRSGSFLCKGSSYTISVNMYCTVFLLTCMSSDRYLAIMYPSVARRIRTKLYSIILCTCVWILSCLLGLPTLLSRELRRYDNSTYCVDKEITSTHRIGSLLLLILAFFVPLFIILTFYCSITKKLCVHYQKSGKHDKKLKKSIKIVFTVVIIFVLSWAPFNIFKFLALMSAIQELKPPFCLPYKVAHIGMELSGPFAFANSCTNPFIYYFFDDYIRRAMMQCITPCTKANNFGTSSDSSDTRLSYSLTAFAHREDVSRKRRRSVSF |
| | *Lacerta agilis* | Gene ID:117045478<br>RNA ID: XM_033146558<br>Protein ID:XP_033002449 | MEKSTSSYDYYFYTDPPEDNCSAVELPYKHIFLPALYTIVFLVGIAGNALLIGALIFKRRVQRLIDIFIVNLAVSDFIFLVTLPFWVDKERTSGLWRSGSFLCKGSSYTISVNMYCTIFLLTCMSSDRYLAIMYPSVARRIRTKLYSIILCTCVWILSCLLGLPTLLSRELRRYDNSTYCVDKEIISTHQIGSLLLLILAFFVPLFIILTFYCSITKKLCVHYQKSGKHDKKLKKSIKIVFTVVIIFVLSWAPFNIFKFLALMSAIQELKPPFCLPYKVAHIGMELSGPFAFANSCTNPFIYYFFDDYIRRAMMQCITPCTKANNFGTSSDSSDTRLSYSLSAFAHREDVSRKRRRSVSF |
| | *Hemicordylus capensis* | Gene ID:128349503<br>RNA ID: XM_053305893<br>Protein ID:XP_053161868 | METTTFTDDYSSGTEAPEDICQGVQLPYKHIFLPALYGTVFLVGIAGNALLMGALLFRRRTQRLIDIFIINLAVSDFIFLITLPFWVDKEMNSGLWRTGSFLCKGSSYIIAVNMYCSIFLLTCMSADRYLAIMHSSMARKIRTKFYSIILCICVWILSCLLGLPTLLSRELRHDYDGNTYCINKESTLTNWIGSLLLLILAFFVPLFSILTFYCSITRKLCVHYQKSGKRDKKLKKSLKVVFIVVIAFVFSWIPYNVFKFLVVISHIQGLKSSFCLPYEVALLGMELSSPFAFANSCTNPFIYYYFDDYIHRAMMQCMLPCMKISSSGTSIDTSDTRLSYSINTSAQGEDNSRKRRRSVSF |

| | | | |
|---|---|---|---|
| | *Heteronotia binoei* | Gene ID:132568979<br>RNA ID: XM_060235164<br>Protein ID:XP_060091147 | MEGSTVNYYDDFFSTEAADNNCPAVLLPYKDIFLPALYASVFLVGIVGNTLLMGALIFKRRIQRLIDIFIVNLAASDFVFLITLPFWVDKEIFSGLWRSGLFICKGSSYIISVNMYCSIFLLTCMSADRYLAIMHPSVARRIRTKLHSVTLCISVWILSCLLGLPTLLSRELGNYNDNTYCEDKKATFINQIGSLLQIILAFFLPLLSILLFYCSITRKLCIHCKKSGKHNKKLKKSIKVVFIVVIAFVFSWAPYNIFKFLSVVSGIQELKPPFCLTYKVAYLGMELSGPFAFANSCTNPLIYYFFDDYIRRAMMQCMLPCVKASSLGTSSDSTDTRLSYSLTVHGEDVSRRRRRSLSL |
| | *Gekko japonicus* | Gene ID:107112445<br>RNA ID: XM_015413575<br>Protein ID:XP_015269061 | MEESTVSYYDDFFSTEAPDDNCPPVQLPYKNIFLPALYATVFLVGIAGNTLLMGALIFKRGIQRLIDIFIVNLAASDFVFLITLPFWVDKEMSSGLWRSGLFICKGSSYIISVNMYCSIFLLTCMSADRYLAIMHPSVARRIRTKLYSATLCICVWILSCLLGLPTLLSRELGNYDGNTYCEDKKATFTDRIGSLLLLVLAFFFPLLSILLFYCSITKKLCIHYKKSGKHNKKLKKSIKVVFVVVIAFVFSWAPYNIFKFLSLVSGIQELKPPFCLIYKVAYLGMELSGPFAFANSCTNPFIYYFFDDYIRRAMVQCMLPCVKASSLGTSSDSTDTRLSYSLTVHGEDVSRRRRRSLSL |
| | *Euleptes europaea* | Gene ID:130485483<br>RNA ID: XM_056858689<br>Protein ID:XP_056714667 | MEGSTVNYYDDYFSTEAPDHNCSAVELPYKDIFLPVLYAGVFLVGIAGNTLLMGALIFKRRIQRLIDIFIVSLAASDFVFLITLPFWVDKEMSSGLWRSGLFTCKGSSYIISVNMYCSIFLLTCMSADRYLAILYPSAARRIRTKLYSVTLCICVWILSCLLGLPTLLSRELGNNDGNAYCEDKKATLTNRIVSLLLLILAFFLPLLSILLFYCSITRKLCIHYKKSGKHNKKLKKSIKVIFIVVIAFVFSWAPFNVFKFLSVVSGIQELKPPFCLTYKVAHLGMELSGPFAFANSCTNPFIYYFFDDYIRGAMLRCTLPCMKASSLGISSDTTDTRLSYSLNVHGEDVSRKRRRSLSL |
| | *Eublepharis macularius* | Gene ID:129326536<br>RNA ID: XM_054974748<br>Protein ID:XP_054830723 | MEGSTVSYYDDFISTEAPDDYCPGVQLPYKGIFLPVLYATVFLVGIVGNGLLMCALIIKRRAQRLIDIFIVNLAASDFVFLITLPFWVDKEMSSGIWRSGSFICKGSSYIISVNMYCSIFLLTCMSADRYLAIIHPSVARRIRTKLYSIILCICVWILSCLLGLPTLLSRELEENDGNTYCKDKKVTSTNRIGSLFLLISAFFFPLLSILLFYCSITRKLCMHYKKAGKHNKKLKKSIKIVFIVVIAFVFSWAPFNIFKFLSVVSGIQELKPPFCLTYEVAYLGMELSGPFAFANSCTNPFIYFFFDGYIRTAMLQCMLPCLKAHSPGTSSDTMDTCLSYSLTVHGEDVSRRRRRSLSL |
| | *Sceloporus undulatus* | Gene ID:121925033<br>RNA ID: XM_042456738<br>Protein ID:XP_042312672 | MEGSTSSYDYFYSFSTEAPEESCPTKELPYKSIVLPVLYVTVFLVGIVGNALLIGALIFKQRIWRLIDIFIVNLAISDFIFLITLPFWVDKEIYSGLWRSGSFVCKGSSYIISVNMYCSIFLLTCMSCDRYLAIMHPSAARKIRTRLYSIILCVGVWILSCLLGLPTLLSRELRRFDDNPYCVDKDTASSQRIESLLLVVFAFFVPLFCILTFYCSITKKLCVHYQKSGKHDKKLRKSIRIVFIVVIAFVFSWAPFNVFKFVALTFGIQNVKPPFCLPYKIAYLGMELGGPLAFANSCTNPFIYYYFDDHIRRAMLRCIFPCIKTSSLRTSSDTSDSHLSYTLTAFAHGEDASRKRRRSVSF |
| | *Anolis sagrei* | Gene ID:132770048<br>RNA ID: XM_067466604<br>Protein ID:XP_067322705 | MEETMSNYYDLYSPGTEAPEEICLPIELPYKNIVLPVLYSTVFFVGILGNTLLIGALVFKQRIQRPIDIFIVNLAASDFIFLITLPIWVDKEIYSGLWRSGSFVCKGSSYIISVNMYCSIFLLTVMSGDRYLAIMYPSVARRIRTKLYSINLCVCVWILSFLLGLPTLLSRELQRYADEYYCMDKDPTPKKRIESLLLLALAFFVPLFSILTFYCSVTKKLCVHYQKSGKHEKKLKKSIKIVFIVVIAFVFSWAPFNVFKIVAVMSGIQDLKPPFCLPYKVGIFGMELSGPFAFANSCTNPLIYYYFDDHIRKAMLRCILPCIKTKSFGSSSDTSDTHS |
| | *Anolis carolinensis* | Gene ID:100567777<br>RNA ID: XM_003219200<br>Protein ID:XP_003219248 | MEETMPSYYDLYFPSSETPEEICLPVEVPYKNIVLPVLYATVFFVGILGNTLLIGALIFKQRIRRPIDIFIVNLAVSDFIFLITLPIWVDKEIYSGLWRSGSFVCKGSSYIISVNMYCSIFLLTVMSGDRYLAIMYPSVARRIRTKLYSINLCVCVWILSFLLGLPTLLSRELQRYADKSYCIDKDPTLKKRIESLLLLALAFFVPLFSILMFYCSVTKKLCVHYQKSGRHEKKLKKSIKIVFIVVIAFVFSWAPFNVFKFVAIMSDIQDLKPPFCLLYKVGNFGMELSGPFAFANSCTNPLIYYYFDDHIRKAMLRCILPCIKTMSSGSSSDTSDTQS |
| | *Terrapene triunguis* | Gene ID:112108285<br>RNA ID: XM_024202343<br>Protein ID:XP_024058111 | MSETPFNYNDSYEYFFPTESPEEYCQALHMPYMGIFLPILYATVFLVGIVGNSILMGALVFKFGVRRLIDTFILNLAASDFVFLLTLPLWVHKEIWLGVWRSGSFLCKGSSYIISVNMYCSIFLLTCMSADRYLAIMYPSVARKVRTRFYTNGLCICVWILSCLLGLPTLLSRELRQYNGQAYCTDVELTPAKRIVSLVMLILAFFFPLLSILTFYCSITKKLCMHYQKSGKHDKKLRKSIKIVFIVVAAFVFSWIPYNIFKLLAIISGLQDLKPPFCLPYVLAQAGMEVSSPFAFANSCANPFIYYCFDGYIRRNISQCLCPWVKGHSTGSSSDTLDTRISYSLSTFIHGEDAIRKRRRSLSF |
| | *Mauremys mutica* | Gene ID:123352090<br>RNA ID: XM_044991865<br>Protein ID:XP_044847800 | MSETIFPYNDSYEYSFPTENPEEYCQALDMPYMGIFLPILYATVFLVGIVGNSILMGALVFKLGIRRLIDIFILNLAASDFVFLLTLPLWVHKEIWQGVWRSGSFLCKGSSYIISVNMYCSIFLLTCMSADRYLAIMYPSVARKVRTRFYTNGLCICVWLLSCLLGLPTLLSRELRQYNGQAYCTDVELTPIKRIVSLVTLILAFFFPLLSILTFYCSITKKLCMHYQKSGKHDKKLRKSIKIVFIVVAAFVFSWIPYNVFKLLAIISGLQDLKPPFCLPYVLAQVGMEVSSPFAFANSCANPFIYYCFDGYIRRNISQRLCPWVKGQSTGSSSDTLDTRSISYSLSTFIHGEDAIRKRRRSLSF |
| | *Chelonia mydas* | Gene ID:102948133<br>RNA ID: XM_007062124<br>Protein ID:XP_007062186 | MSETILSYNDSYEYSFSTESPEDYCQALSIPYMGIFLPILYATVFLVGIVGNAILMGALIFKFRVRRLIDIFILNLAASDFVFLLTLPLWVHKEIWLGVWRSGSFLCKGSSYIISVNMYCSIFLLTCMSADRYLAIIHPSVARKVRTRFYTNGLCICVWILSCLLGLPTLLSRELRQYNGQAYCTDVELTPAKRIVSLVTLILAFFFPLLSILTFYCSITKKLCMHYQKSGKHDKKLRKSIKIVFIVVAAFVFSWIPYNIFKLLAIISGLQDLKPPFCLPYVLAQVGMEVSSPFAFANSCANPFIYYCFDGYIHRNILRCLCPWVKGHSTGSSSDSLNTRISHSMSTFINGEDAIRKRRRSLSL |
| | *Caretta caretta* | Gene ID:125630184<br>RNA ID: XM_048835776<br>Protein ID:XP_048691733 | MSETTLSYNDSYEYSFSTESPEEYCQALYIPYMGIFLPILYATVFLVGIVGNAILMGALVFKFRVRRLIDIFILNLAASDFVFLLTLPLWVHKEIWLGVWRSGSFLCKGSSYIISVNMYCSIFLLTCMSADRYLAIMHPSVARKVRTRFYTNGLCICVWILSCLLGLPTLLSRELRQYNGQAYCIDVDLTPAKRIVSLVTLILAFFFPLLSILTFYCSITKKLCMHYQKSGKHDKKLRKSIKIVFIVVAAFVFSWIPYNIFKLLAIISGLQDLKPPFCLPYVLAQAGMEVSSPFAFANSCANPFIYYCFDGYIHRNILRCLCPWVKGHSTGSSSDSLNTRISHSMSTFIHGEDTIRKRRRSLSL |
| | *Gavialis gangeticus* | Gene ID:109304298<br>RNA ID: XM_019526777<br>Protein ID:XP_019382322 | MDSEYLYPMMTGTSYAYDYSLPTESPEESCQAEPVLYANIFLPLLYAIVFVVGIIGNSILIAALFRRGIQRLIDIFIANLAASDFVFLITLPLWVDKEKSGGNWRSGSFLCKGSSYVISVNMYCSTLLLTCMSTDRYLAIMHPSVARKVRTRFNSIGMCICIWVLSGFLGLPTLLTRQLVLDEDTGQSYCTDSEQMPMKWITSLIILILAFFFPLLCILTFYCSVTRKLCMHYQKSGKHNRKLRKSIKTVFIVVAAFVVSWIPFNLFKLLANISRLQHPNPLFCFPYKVAQIGMQVSGPLAFANSCANPFIYYFFDCYMRRAMTRCIYSQVKAHSVRSSSDTMDTRLSYSLSYFIYGEDATRKRRRSMSF |
| | *Crocodylus porosus* | Gene ID:109319150<br>RNA ID: XM_019548656<br>Protein ID:XP_019404201 | MDSEYLYPVTTGTSYAYDYSLPTESPEESCQAEPVPYANIFLPLLYAIVFVVGIIGNSILIAAVFRRGIQRLIDIFIANLAASDFVFLITLPLWVDKEKSGGSWRSGSFLCKGSSYVISVNMYCSTLLLTCMSTDRYLAIMHPSVARKVRTRFNSIGMCICIWVLSGFLGLPTLLTRQLVLEEETGQSYCTDSEQMPMKWITSLIILILAFFFPLLCILTFYCSVTRKLCMHYQKSGKHDRKLRKSIKIVFIVVAAFVFSWIPFNLFKLLANISRLQHPLFCFPYKVAQMGMQVSGPLAFANSCANPFIYYFFDCYMRRAMTRCIYSQVKAHSVRSSSDTMDTRLSHSLSYFIYGEDATRKRRRSMSF |

| | | | |
|---|---|---|---|
| | *Alligator sinensis* | Gene ID:102368112<br>RNA ID: XM_006020153<br>Protein ID:XP_006020215 | MTGTSYVAYDDYSLPTESPEESCQAEPVPHANIFLPLLYAIIFVVGIIGNSILIAALFRRGVQRLIDVFIVNLAASDFVFLITLPLWVDKEKSGGNWRSGSFLCKGSSYVISVNMYCSTLLLTCMSTDRYLAIMHPFVARKVRTRFNSIGMCICVWVLSGFLGLPILLTRQLVLDKETGKSHCTDSDLMPMRWITSLIILILAFFFPLLCILTFYCSVTRKLCMHYQKSGKHDKKLRKSIKIVFIVVAAFVFSWIPFNLFKLLAIISQLLGPNLPCLPYKVALVGMQVSGPLAFANSCANPFIYYFFDCYIRRAMMRCIYSQVKAHSVRSSSDTMDTRLSHSLSYFIYGEDATRKRRRSMSF |
| Amphibians | *Microcaecilia unicolor* | Gene ID:115460609<br>RNA ID: XM_030190377<br>Protein ID:XP_030046237 | MENVAVMDYLDYGSTPLYDNSSSEEECELSHLPYTSVSLSLIYSILFLLGTAGNIILIGALSFRKHTVRLVDIFVINLAISDLVFVVTLPLWVDREVSDGAWRSGSFLCKASSYIISVNMYSSIFFLGCMSLDRYLAIVHPLHSRKLRTRFYACLFCTLVWLVSSILGIPILVFTKWIMLEDGAAYCVDMEVTSTRLILSLLSLILAFFLPLITILTFYCSITKKLCLHYCRAGKQDRKLRRSFKIIVIVVAIFICSWVPYNTFRLLGILNELLQEPSCMAATVAQLGIETSGPIAFTNSCVNPIIYYVFDGYIRRSIKHFLCFCAAPGRLKRSSVTSETXLSKSLAVHLQSKEKNWTRKRKLSISF |
| Fishes | *Polypterus senegalus* | Gene ID:120538035<br>RNA ID: XM_039767425<br>Protein ID:XP_039623359 | MEYSSSTVEYETYYYENDSTEQSEGCRLPAFPWTWIIRTFLYILVVILGVPGNIALIWIMVRRLSVFRRPCESFVVNLAISDLLLLLGLLVWIDSEIHGGSWRSGWLCCKITAYFMALSMQSGIFFLTAMSIDRYLAVVHSNIYRKIKKKLYVTASCFLVWLVSILIALPVFRARTLSLDVNGIWRCREEIDIHLQRFSLVNLLAFFFSLLGILYCYCSIMRTLCLHYRRTRRQNHKLQRSIKVVFLVVVVFCFSWVPFNVFKIVKIVLTMMNKENSCTMDVALTGLGVLVPFAFSNSCANPFIYTWADASLKKLAVRCLCPCLPRLQEVMVVSQSSEMGSRSSQGSESSWHKKERQNTTNCQLSVKA |
| | *Lepisosteus oculatus* | Gene ID:102691053<br>RNA ID: XM_006643117<br>Protein ID:XP_006643180 | MTATTDAVVLGAQMSGPAEATTSGPWQEETTYDYDANASGCEVPVLAGSARARVGLYCAFFVAGVAGNLALLAALWARWRRRGWGRQRGWRPSETLAANLAAADLLFLTALPFWIDSELSGGTWRAGELSCKGAPFLVALSMNVGVLMLTFVSVDRYLAVVNPSLYRRLSRALFTGLGCLSVWLVSPLLALPVLRARVLQGVELEDGTEAMWCQEEDGSSSPGQSLLLLLFSFFLPLLLVLLCYCRITRTLCLHARRSSSLDTHLCRSFKIIFLVVGAFVLSWVPFNTFRLVGVVQQLLARDSSPSCVAHKVAQLGMEITAPLAFANSCANPFIYALADRSLRRDSARCLCPCLVKAPVSWGAISFSSQQGASWSGWRTRDRAAGEPVGLTTLQD |
| | *Latimeria chalumnae* | Gene ID:102351034<br>RNA ID: XM_006012249<br>Protein ID:XP_006012311 | MGDSVEHTYDDDKYYLFTVHYDYFPSTSFPPEDQCSDFDLSLPHGLLPILYCLVFVVGAVGNAIVIGAIVFKRGVKRLVDIFISHLAVSDFIFLVTLPLWVDKEVVGGPWRSGWFFCKFSAYIITLNMYSSVFFLTCMSLDRFLAAVLPLQSRVFRTKHNAKVCCTCVWMLSATLAAPVLHSRVLKKYEEKEYCNEDAGSSTVAFSMISLIVAFFLPLAVILSCYCTIIWKLCHQWQKFHKQQQKLRRSLKIVFIVVVVFVFSWMPFNLFRALAIANNLQVEVTCQSYTLARLGMQLTAPLAFSNSCANPIIYAFFDRYIRRAMLQCLCPCIKPPAHWQSSDTSESHLHKSHPSAEKGRRDNSR |

**6×His-SUMO-Lo-GPR15LG**

```
1    ATG CAT CAC CAT CAC CAC CAT ATG GCT AGC ATG TCG GAC TCA GAA GTC AAT CAA GAA GCT AAG CCA GAG GTC AAG
     TAC GTA GTG GTA GTG GTG GTA TAC CGA TCG TAC AGC CTG AGT CTT CAG TTA GTT CTT CGA TTC GGT CTC CAG TTC
      M   H   H   H   H   H   H   M   A   S   M   S   D   S   E   V   N   Q   E   A   K   P   E   V   K

76   CCA GAA GTC AAG CCT GAG ACT CAC ATC AAT TTA AAG GTG TCC GAT GGA TCT TCA GAG ATC TTC TTC AAG ATC AAA
     GGT CTT CAG TTC GGA CTC TGA GTG TAG TTA AAT TTC CAC AGG CTA CCT AGA AGT CTC TAG AAG AAG TTC TAG TTT
      P   E   V   K   P   E   T   H   I   N   L   K   V   S   D   G   S   S   E   I   F   F   K   I   K

151  AAG ACC ACT CCT TTA AGA AGG CTG ATG GAA GCG TTC GCT AAA AGA CAG GGT AAG GAA ATG GAC TCC TTA AGA TTC
     TTC TGG TGA GGA AAT TCT TCC GAC TAC CTT CGC AAG CGA TTT TCT GTC CCA TTC CTT TAC CTG AGG AAT TCT AAG
      K   T   T   P   L   R   R   L   M   E   A   F   A   K   R   Q   G   K   E   M   D   S   L   R   F

226  TTG TAC GAC GGT ATT AGA ATT CAA GCT GAT CAG ACC CCT GAA GAT TTG GAC ATG GAG GAT AAC GAT ATT ATT GAG
     AAC ATG CTG CCA TAA TCT TAA GTT CGA CTA GTC TGG GGA CTT CTA AAC CTG TAC CTC CTA TTG CTA TAA TAA CTC
      L   Y   D   G   I   R   I   Q   A   D   Q   T   P   E   D   L   D   M   E   D   N   D   I   I   E

301  GCT CAC AGA GAA CAG ATT GGT GGA CGT AAA CTG AAG TGT TGC AAG AAA TAT TTT CTG AAA CAT CAC AAG AAC GAC
     CGA GTG TCT CTT GTC TAA CCA CCT GCA TTT GAC TTC ACA ACG TTC TTT ATA AAA GAC TTT GTA GTG TTC TTG CTG
      A   H   R   E   Q   I   G   G   R   K   L   K   C   C   K   K   Y   F   L   K   H   H   K   N   D

376  TCC TTA CGT CCG AAG AAT GCC AAA ACC GGT CAT CGC TGC CGT CCA TGT CGC CCT AAC ATT CCG TTG CCG TCT AGC
     AGG AAT GCA GGC TTC TTA CGG TTT TGG CCA GTA GCG ACG GCA GGT ACA GCG GGA TTG TAA GGC AAC GGC AGA TCG
      S   L   R   P   K   N   A   K   T   G   H   R   C   R   P   C   R   P   N   I   P   L   P   S   S

451  TAA
     ATT
      *
```

**6×His-SUMO-SmBiT-Lo-GPR15LG**

```
1    ATG CAT CAC CAT CAC CAC CAT ATG GCT AGC ATG TCG GAC TCA GAA GTC AAT CAA GAA GCT AAG CCA GAG GTC AAG
     TAC GTA GTG GTA GTG GTG GTA TAC CGA TCG TAC AGC CTG AGT CTT CAG TTA GTT CTT CGA TTC GGT CTC CAG TTC
      M   H   H   H   H   H   H   M   A   S   M   S   D   S   E   V   N   Q   E   A   K   P   E   V   K

76   CCA GAA GTC AAG CCT GAG ACT CAC ATC AAT TTA AAG GTG TCC GAT GGA TCT TCA GAG ATC TTC TTC AAG ATC AAA
     GGT CTT CAG TTC GGA CTC TGA GTG TAG TTA AAT TTC CAC AGG CTA CCT AGA AGT CTC TAG AAG AAG TTC TAG TTT
      P   E   V   K   P   E   T   H   I   N   L   K   V   S   D   G   S   S   E   I   F   F   K   I   K

151  AAG ACC ACT CCT TTA AGA AGG CTG ATG GAA GCG TTC GCT AAA AGA CAG GGT AAG GAA ATG GAC TCC TTA AGA TTC
     TTC TGG TGA GGA AAT TCT TCC GAC TAC CTT CGC AAG CGA TTT TCT GTC CCA TTC CTT TAC CTG AGG AAT TCT AAG
      K   T   T   P   L   R   R   L   M   E   A   F   A   K   R   Q   G   K   E   M   D   S   L   R   F

226  TTG TAC GAC GGT ATT AGA ATT CAA GCT GAT CAG ACC CCT GAA GAT TTG GAC ATG GAG GAT AAC GAT ATT ATT GAG
     AAC ATG CTG CCA TAA TCT TAA GTT CGA CTA GTC TGG GGA CTT CTA AAC CTG TAC CTC CTA TTG CTA TAA TAA CTC
      L   Y   D   G   I   R   I   Q   A   D   Q   T   P   E   D   L   D   M   E   D   N   D   I   I   E

301  GCT CAC AGA GAA CAG ATT GGT GGA GTG ACC GGC TAC CGT CTG TTT GAA GAA ATT CTG GGC GGC AGC GGT GGT GGC
     CGA GTG TCT CTT GTC TAA CCA CCT CAC TGG CCG ATG GCA GAC AAA CTT CTT TAA GAC CCG CCG TCG CCA CCA CCG
      A   H   R   E   Q   I   G   G   V   T   G   Y   R   L   F   E   E   I   L   G   G   S   G   G   G

376  CGT AAA CTG AAG TGT TGC AAG AAA TAT TTT CTG AAA CAT CAC AAG AAC GAC TCC TTA CGT CCG AAG AAT GCC AAA
     GCA TTT GAC TTC ACA ACG TTC TTT ATA AAA GAC TTT GTA GTG TTC TTG CTG AGG AAT GCA GGC TTC TTA CGG TTT
      R   K   L   K   C   C   K   K   Y   F   L   K   H   H   K   N   D   S   L   R   P   K   N   A   K

451  ACC GGT CAT CGC TGC CGT CCA TGT CGC CCT AAC ATT CCG TTG CCG TCT AGC TAA
     TGG CCA GTA GCG ACG GCA GGT ACA GCG GGA TTG TAA GGC AAC GGC AGA TCG ATT
      T   G   H   R   C   R   P   C   R   P   N   I   P   L   P   S   S   *
```

**Fig. S1.** The nucleotide sequence and amino acid sequence of the Lo-GPR15LG expression constructs. The amino acid sequence of Lo-GPR15LG is shown in red, that of SmBiT in blue.

```
1   ATG GGC AGC AGC CAT CAT CAT CAT CAT CAC AGC AGC GGC CTG GTG CCG CGC GGC AGC CAT ATG GCT AGC ATG ACT
    TAC CCG TCG TCG GTA GTA GTA GTA GTA GTG TCG TCG CCG GAC CAC GGC GCG CCG TCG GTA TAC CGA TCG TAC TGA
     M   G   S   S   H   H   H   H   H   H   S   S   G   L   V   P   R   G   S   H   M   A   S   M   T

76  GGT GGA CAG CAA ATG GGT CGC GGA TCC CTT GTT CCT GAA CTG AAC GAA AAA GAT GAT GAC CAG GTA CAA AAA GCT
    CCA CCT GTC GTT TAC CCA GCG CCT AGG GAA CAA GGA CTT GAC TTG CTT TTT CTA CTA CTG GTC CAT GTT TTT CGA
     G   G   Q   Q   M   G   R   G   S   L   V   P   E   L   N   E   K   D   D   D   Q   V   Q   K   A

151 CTG GAT TCT CGT GAA AAC ACT CAG TTA ATG AAT CGT GAT AAC ATC GAG ATC ACA GTA CGT GAC TTT AAG ACC TTG
    GAC CTA AGA GCA CTT TTG TGA GTC AAT TAC TTA GCA CTA TTG TAG CTC TAG TGT CAT GCA CTG AAA TTC TGG AAC
     L   D   S   R   E   N   T   Q   L   M   N   R   D   N   I   E   I   T   V   R   D   F   K   T   L

226 GAA CCA CGC CGT TGG CTG AAT GAC ACT ATC ATT GAG TTC TTT ATG AAA TAC ATT GAA AAA TCT ACC CCT AAC ACC
    CTT GGT GCG GCA ACC GAC TTA CTG TGA TAG TAA CTC AAG AAA TAC TTT ATG TAA CTT TTT AGA TGG GGA TTG TGG
     E   P   R   R   W   L   N   D   T   I   I   E   F   F   M   K   Y   I   E   K   S   T   P   N   T

301 GTG GCG TTC AAC AGC TTT TTC TAT ACC AAC TTA AGC GAA CGC GGT TAT CAA GGC GTC CGC CGC TGG ATG AAG CGT
    CAC CGC AAG TTG TCG AAA AAG ATA TGG TTG AAT TCG CTT GCG CCA ATA GTT CCG CAG GCG GCG ACC TAC TTC GCA
     V   A   F   N   S   F   F   Y   T   N   L   S   E   R   G   Y   Q   G   V   R   R   W   M   K   R

376 AAG AAG ACG CAG ATT GAT AAA CTT GAT AAA ATC TTT ACA CCG ATC AAT CTG AAC CAG TCC CAC TGG GCG TTG GGC
    TTC TTC TGC GTC TAA CTA TTT GAA CTA TTT TAG AAA TGT GGC TAG TTA GAC TTG GTC AGG GTG ACC CGC AAC CCG
     K   K   T   Q   I   D   K   L   D   K   I   F   T   P   I   N   L   N   Q   S   H   W   A   L   G

451 ATC ATT GAT CTG AAA AAG AAA ACT ATC GGT TAC GTA GAT TCA TTA TCG AAC GGT CCG AAT GAT TCC AGC AAA CAG
    TAG TAA CTA GAC TTT TTC TTT TGA TAG CCA ATG CAT CTA AGT AAT AGC TTG CCA GGC TTA CTA AGG TCG TTT GTC
     I   I   D   L   K   K   K   T   I   G   Y   V   D   S   L   S   N   G   P   N   D   S   S   K   Q

526 ATC CTG ACT GAC CTG CAG AAA TAT GTT GAA GAG GAA AGC AAG CAT ACG ATC GGT GAA GAC TTT GAT CTG CGT CAT
    TAG GAC TGA CTG GAC GTC TTT ATA CAA CTT CTC CTT TCG TTC GTA TGC TAG CCA CTT CTG AAA CTA GAC GCA GTA
     I   L   T   D   L   Q   K   Y   V   E   E   E   S   K   H   T   I   G   E   D   F   D   L   R   H

601 CTG GAT TGT CCG CAG CAA CCA AAT GGC TAC GAC TGT GGC ATC TAC GTT TGC ATG AAC ACT CTC TAT GGT AGT GCA
    GAC CTA ACA GGC GTC GTT GGT TTA CCG ATG CTG ACA CCG TAG ATG CAA ACG TAC TTG TGA GAG ATA CCA TCA CGT
     L   D   C   P   Q   Q   P   N   G   Y   D   C   G   I   Y   V   C   M   N   T   L   Y   G   S   A

676 GAT GCG CCG TTG GAT TTC GAC TCT AAA GAT GCG GAA CGC ATG CGT CGT TTT ATT GCC CAT CTG ATT TTA ACC GAC
    CTA CGC GGC AAC CTA AAG CTG AGA TTT CTA CGC CTT GCG TAC GCA GCA AAA TAA CGG GTA GAC TAA AAT TGG CTG
     D   A   P   L   D   F   D   S   K   D   A   E   R   M   R   R   F   I   A   H   L   I   L   T   D

751 GCT CTG AAA TAA GCG GCC GC
    CGA GAC TTT ATT CGC CGG CG
     A   L   K   *
```

**Fig. S2.** The nucleotide sequence and amino acid sequence of the overexpressed R3-ULP1. The BamHI and NotI cleavage sites are shaded.

### Untagged Lo-GPR15 in pcDNA3.1(+)

```
1    GCT AGC ATG ACC GCC ACC ACC GAC GCC GTG GTT CTG GGC GCC CAG ATG AGC GGC CCT GCC GAG GCC ACA ACA AGC
     CGA TCG TAC TGG CGG TGG TGG CTG CGG CAC CAA GAC CCG CGG GTC TAC TCG CCG GGA CGG CTC CGG TGT TGT TCG
              M   T   A   T   T   D   A   V   V   L   G   A   Q   M   S   G   P   A   E   A   T   T   S

76   GGA CCT TGG CAG GAG GAA ACC ACA TAC GAC TAC GAC GCC AAT GCC TCT GGA TGT GAA GTG CCT GTG CTG GCC GGA
     CCT GGA ACC GTC CTC CTT TGG TGT ATG CTG ATG CTG CGG TTA CGG AGA CCT ACA CTT CAC GGA CAC GAC CGG CCT
      G   P   W   Q   E   E   T   T   Y   D   Y   D   A   N   A   S   G   C   E   V   P   V   L   A   G

151  AGC GCC AGA GCC AGA GTG GGA CTG TAC TGC GCC TTC TTC GTG GCC GGC GTT GCT GGC AAC CTG GCA CTG CTC GCC
     TCG CGG TCT CGG TCT CAC CCT GAC ATG ACG CGG AAG AAG CAC CGG CCG CAA CGA CCG TTG GAC CGT GAC GAG CGG
      S   A   R   A   R   V   G   L   Y   C   A   F   F   V   A   G   V   A   G   N   L   A   L   L   A

226  GCC CTG TGG GCC CGT TGG CGG CGG AGA GGC TGG GGC AGA CAG AGA GGC TGG CGG CCT TCT GAA ACC CTG GCC GCC
     CGG GAC ACC CGG GCA ACC GCC GCC TCT CCG ACC CCG TCT GTC TCT CCG ACC GCC GGA AGA CTT TGG GAC CGG CGG
      A   L   W   A   R   W   R   R   R   G   W   G   R   Q   R   G   W   R   P   S   E   T   L   A   A

301  AAC CTG GCT GCT GCT GAT CTG CTG TTC CTG ACA GCC CTT CCA TTT TGG ATC GAC TCT GAG CTG AGC GGC GGA ACA
     TTG GAC CGA CGA CGA CTA GAC GAC AAG GAC TGT CGG GAA GGT AAA ACC TAG CTG AGA CTC GAC TCG CCG CCT TGT
      N   L   A   A   A   D   L   L   F   L   T   A   L   P   F   W   I   D   S   E   L   S   G   G   T

376  TGG AGA GCC GGC GAG CTG TCT TGT AAA GGC GCC CCT TTC CTG GTG GCC CTG TCC ATG AAC GTG GGC GTG CTG ATG
     ACC TCT CGG CCG CTC GAC AGA ACA TTT CCG CGG GGA AAG GAC CAC CGG GAC AGG TAC TTG CAC CCG CAC GAC TAC
      W   R   A   G   E   L   S   C   K   G   A   P   F   L   V   A   L   S   M   N   V   G   V   L   M

451  CTG ACA TTC GTG TCT GTG GAC CGG TAC CTG GCG GTC GTG AAC CCT AGC CTG TAT AGA CGG CTG AGC AGA GCT CTC
     GAC TGT AAG CAC AGA CAC CTG GCC ATG GAC CGC CAG CAC TTG GGA TCG GAC ATA TCT GCC GAC TCG TCT CGA GAG
      L   T   F   V   S   V   D   R   Y   L   A   V   V   N   P   S   L   Y   R   R   L   S   R   A   L

526  TTC ACC GGC CTG GGC TGT CTG TCC GTG TGG CTG GTC AGC CCT CTG CTG GCC CTG CCC GTG CTG AGA GCC CGG GTG
     AAG TGG CCG GAC CCG ACA GAC AGG CAC ACC GAC CAG TCG GGA GAC GAC CGG GAC GGG CAC GAC TCT CGG GCC CAC
      F   T   G   L   G   C   L   S   V   W   L   V   S   P   L   L   A   L   P   V   L   R   A   R   V

601  CTG CAA GGC GTG GAA CTG GAG GAT GGC ACC GAG GCC ATG TGG TGC CAG GAG GAG GAC GGC AGC TCC AGC CCC GGC
     GAC GTT CCG CAC CTT GAC CTC CTA CCG TGG CTC CGG TAC ACC ACG GTC CTC CTC CTG CCG TCG AGG TCG GGG CCG
      L   Q   G   V   E   L   E   D   G   T   E   A   M   W   C   Q   E   E   D   G   S   S   S   P   G

676  CAG AGC CTG CTG CTG CTG CTG TTT AGC TTT TTC CTG CCT CTG CTG CTG GTG CTG CTG TGC TAC TGC AGA ATT ACC
     GTC TCG GAC GAC GAC GAC GAC AAA TCG AAA AAG GAC GGA GAC GAC GAC CAC GAC GAC ACG ATG ACG TCT TAA TGG
      Q   S   L   L   L   L   L   F   S   F   F   L   P   L   L   L   V   L   L   C   Y   C   R   I   T

751  AGA ACT CTG TGC CTG CAC GCC AGG CGG AGC TCT TCT CTT GAT ACA CAC CTG TGC CGG AGC TTC AAG ATC ATC TTC
     TCT TGA GAC ACG GAC GTG CGG TCC GCC TCG AGA AGA GAA CTA TGT GTG GAC ACG GCC TCG AAG TTC TAG TAG AAG
      R   T   L   C   L   H   A   R   R   S   S   S   L   D   T   H   L   C   R   S   F   K   I   I   F

826  CTG GTG GTG GGC GCC TTC GTG CTG TCC TGG GTC CCC TTC AAC ACC TTT AGA CTG GTG GGC GTG GTG CAG CAG CTG
     GAC CAC CAC CCG CGG AAG CAC GAC AGG ACC CAG GGG AAG TTG TGG AAA TCT GAC CAC CCG CAC CAC GTC GTC GAC
      L   V   V   G   A   F   V   L   S   W   V   P   F   N   T   F   R   L   V   G   V   V   Q   Q   L

901  CTG GCT AGA GAC AGC AGC CCC AGC TGC GTG GCC CAC AAG GTG GCT CAA CTG GGA ATG GAA ATC ACC GCC CCT CTG
     GAC CGA TCT CTG TCG TCG GGG TCG ACG CAC CGG GTG TTC CAC CGA GTT GAC CCT TAC CTT TAG TGG CGG GGA GAC
      L   A   R   D   S   S   P   S   C   V   A   H   K   V   A   Q   L   G   M   E   I   T   A   P   L

976  GCT TTC GCC AAT AGC TGC GCC AAC CCC TTC ATC TAC GCC CTG GCC GAC AGA AGC CTC AGA AGA GAT TCT GCC CGG
     CGA AAG CGG TTA TCG ACG CGG TTG GGG AAG TAG ATG CGG GAC CGG CTG TCT TCG GAG TCT TCT CTA AGA CGG GCC
      A   F   A   N   S   C   A   N   P   F   I   Y   A   L   A   D   R   S   L   R   R   D   S   A   R

1051 TGC CTG TGT CCT TGC CTG GTG AAG GCC CCA GTG TCC TGG GGA GCC ATC AGC TTC AGC AGC CAG CAG GGC GCC AGC
     ACG GAC ACA GGA ACG GAC CAC TTC CGG GGT CAC AGG ACC CCT CGG TAG TCG AAG TCG TCG GTC GTC CCG CGG TCG
      C   L   C   P   C   L   V   K   A   P   V   S   W   G   A   I   S   F   S   S   Q   Q   G   A   S

1126 TGG TCC GGC TGG CGC ACC AGA GAT AGA GCC GCT GGC GAG CCT GTG GGC CTG ACC ACC CTG CAG GAC TGA GCGGCCGC
     ACC AGG CCG ACC GCG TGG TCT CTA TCT CGG CGA CCG CTC GGA CAC CCG GAC TGG TGG GAC GTC CTG ACT CGCCGGCG
      W   S   G   W   R   T   R   D   R   A   A   G   E   P   V   G   L   T   T   L   Q   D   *
```

### sLgBiT-Lo-GPR15 in PB-TRE

1 GCT AGC ATG AAC TCC TTC TCC ACA AGC GCC TTC GGT CCA GTT GCC TTC TCC CTG GGC CTG CTC CTG GTG TTG CCT
CGA TCG TAC TTG AGG AAG AGG TGT TCG CGG AAG CCA GGT CAA CGG AAG AGG GAC CCG GAC GAG GAC CAC AAC GGA
M N S F S T S A F G P V A F S L G L L L V L P

76 GCT GCC TTC CCT GCC CCA GTC TTC ACA CTC GAA GAT TTC GTT GGG GAC TGG GAA CAG ACA GCC GCC TAC AAC CTG
CGA CGG AAG GGA CGG GGT CAG AAG TGT GAG CTT CTA AAG CAA CCC CTG ACC CTT GTC TGT CGG CGG ATG TTG GAC
A A F P A P V F T L E D F V G D W E Q T A A Y N L

151 GAC CAA GTC CTT GAA CAG GGA GGT GTG TCC AGT TTG CTG CAG AAT CTC GCC GTG TCC GTA ACT CCG ATC CAA AGG
CTG GTT CAG GAA CTT GTC CCT CCA CAC AGG TCA AAC GAC GTC TTA GAG CGG CAC AGG CAT TGA GGC TAG GTT TCC
D Q V L E Q G G V S S L L Q N L A V S V T P I Q R

226 ATT GTC CGG AGC GGT GAA AAT GCC CTG AAG ATC GAC ATC CAT GTC ATC ATC CCG TAT GAA GGT CTG AGC GCC GAC
TAA CAG GCC TCG CCA CTT TTA CGG GAC TTC TAG CTG TAG GTA CAG TAG TAG GGC ATA CTT CCA GAC TCG CGG CTG
I V R S G E N A L K I D I H V I I P Y E G L S A D

301 CAA ATG GCC CAG ATC GAA GAG GTG TTT AAG GTG GTG TAC CCT GTG GAT GAT CAT CAC TTT AAG GTG ATC CTG CCC
GTT TAC CGG GTC TAG CTT CTC CAC AAA TTC CAC CAC ATG GGA CAC CTA CTA GTA GTG AAA TTC CAC TAG GAC GGG
Q M A Q I E E V F K V V Y P V D D H H F K V I L P

376 TAT GGC ACA CTG GTA ATC GAC GGG GTT ACG CCG AAC ATG CTG AAC TAT TTC GGA CGG CCG TAT GAA GGC ATC GCC
ATA CCG TGT GAC CAT TAG CTG CCC CAA TGC GGC TTG TAC GAC TTG ATA AAG CCT GCC GGC ATA CTT CCG TAG CGG
Y G T L V I D G V T P N M L N Y F G R P Y E G I A

451 GTG TTC GAC GGC AAA AAG ATC ACT GTA ACA GGG ACC CTG TGG AAC GGC AAC AAA ATT ATC GAC GAG CGC CTG ATC
CAC AAG CTG CCG TTT TTC TAG TGA CAT TGT CCC TGG GAC ACC TTG CCG TTG TTT TAA TAG CTG CTC GCG GAC TAG
V F D G K K I T V T G T L W N G N K I I D E R L I

526 ACC CCC GAC GGC TCC ATG CTG TTC CGA GTA ACC ATC AAC AGT GGT GGC GGC TCT GGT GGT GGC AGC GGC GGT GGT
TGG GGG CTG CCG AGG TAC GAC AAG GCT CAT TGG TAG TTG TCA CCA CCG CCG AGA CCA CCA CCG TCG CCG CCA CCA
T P D G S M L F R V T I N S G G G S G G G S G G G

601 ACC ACC GCC ACC ACC GAC GCC GTG GTT CTG GGC GCC CAG ATG AGC GGC CCT GCC GAG GCC ACA ACA AGC GGA CCT
TGG TGG CGG TGG TGG CTG CGG CAC CAA GAC CCG CGG GTC TAC TCG CCG GGA CGG CTC CGG TGT TGT TCG CCT GGA
T T A T T D A V V L G A Q M S G P A E A T T S G P

676 TGG CAG GAG GAA ACC ACA TAC GAC TAC GAC GCC AAT GCC TCT GGA TGT GAA GTG CCT GTG CTG GCC GGA AGC GCC
ACC GTC CTC CTT TGG TGT ATG CTG ATG CTG CGG TTA CGG AGA CCT ACA CTT CAC GGA CAC GAC CGG CCT TCG CGG
W Q E E T T Y D Y D A N A S G C E V P V L A G S A

751 AGA GCC AGA GTG GGA CTG TAC TGC GCC TTC TTC GTG GCC GGC GTT GCT GGC AAC CTG GCA CTG CTC GCC GCC CTG
TCT CGG TCT CAC CCT GAC ATG ACG CGG AAG AAG CAC CGG CCG CAA CGA CCG TTG GAC CGT GAC GAG CGG CGG GAC
R A R V G L Y C A F F V A G V A G N L A L L A A L

826 TGG GCC CGT TGG CGG CGG AGA GGC TGG GGC AGA CAG AGA GGC TGG CGG CCT TCT GAA ACC CTG GCC GCC AAC CTG
ACC CGG GCA ACC GCC GCC TCT CCG ACC CCG TCT GTC TCT CCG ACC GCC GGA AGA CTT TGG GAC CGG CGG TTG GAC
W A R W R R R G W G R Q R G W R P S E T L A A N L

901 GCT GCT GCT GAT CTG CTG TTC CTG ACA GCC CTT CCA TTT TGG ATC GAC TCT GAG CTG AGC GGC GGA ACA TGG AGA
CGA CGA CGA CTA GAC GAC AAG GAC TGT CGG GAA GGT AAA ACC TAG CTG AGA CTC GAC TCG CCG CCT TGT ACC TCT
A A A D L L F L T A L P F W I D S E L S G G T W R

976 GCC GGC GAG CTG TCT TGT AAA GGC GCC CCT TTC CTG GTG GCC CTG TCC ATG AAC GTG GGC GTG CTG ATG CTG ACA
CGG CCG CTC GAC AGA ACA TTT CCG CGG GGA AAG GAC CAC CGG GAC AGG TAC TTG CAC CCG CAC GAC TAC GAC TGT
A G E L S C K G A P F L V A L S M N V G V L M L T

1051 TTC GTG TCT GTG GAC CGG TAC CTG GCG GTC GTG AAC CCT AGC CTG TAT AGA CGG CTG AGC AGA GCT CTC TTC ACC
AAG CAC AGA CAC CTG GCC ATG GAC CGC CAG CAC TTG GGA TCG GAC ATA TCT GCC GAC TCG TCT CGA GAG AAG TGG
F V S V D R Y L A V V N P S L Y R R L S R A L F T

1126 GGC CTG GGC TGT CTG TCC GTG TGG CTG GTC AGC CCT CTG CTG GCC CTG CCC GTG CTG AGA GCC CGG GTG CTG CAA
CCG GAC CCG ACA GAC AGG CAC ACC GAC CAG TCG GGA GAC GAC CGG GAC GGG CAC GAC TCT CGG GCC CAC GAC GTT
G L G C L S V W L V S P L L A L P V L R A R V L Q

1201 GGC GTG GAA CTG GAG GAT GGC ACC GAG GCC ATG TGG TGC CAG GAG GAG GAC GGC AGC TCC AGC CCC GGC CAG AGC
CCG CAC CTT GAC CTC CTA CCG TGG CTC CGG TAC ACC ACG GTC CTC CTC CTG CCG TCG AGG TCG GGG CCG GTC TCG
G V E L E D G T E A M W C Q E E D G S S S P G Q S

1276 CTG CTG CTG CTG CTG TTT AGC TTT TTC CTG CCT CTG CTG CTG GTG CTG CTG TGC TAC TGC AGA ATT ACC AGA ACT
GAC GAC GAC GAC GAC AAA TCG AAA AAG GAC GGA GAC GAC GAC CAC GAC GAC ACG ATG ACG TCT TAA TGG TCT TGA

```
     L   L   L   L   L   F   S   F   F   L   P   L   L   L   V   L   L   C   Y   C   R   I   T   R   T

1351 CTG TGC CTG CAC GCC AGG CGG AGC TCT TCT CTT GAT ACA CAC CTG TGC CGG AGC TTC AAG ATC ATC TTC CTG GTG
     GAC ACG GAC GTG CGG TCC GCC TCG AGA AGA GAA CTA TGT GTG GAC ACG GCC TCG AAG TTC TAG TAG AAG GAC CAC
      L   C   L   H   A   R   R   S   S   S   L   D   T   H   L   C   R   S   F   K   I   I   F   L   V

1426 GTG GGC GCC TTC GTG CTG TCC TGG GTC CCC TTC AAC ACC TTT AGA CTG GTG GGC GTG GTG CAG CAG CTG CTG GCT
     CAC CCG CGG AAG CAC GAC AGG ACC CAG GGG AAG TTG TGG AAA TCT GAC CAC CCG CAC CAC GTC GTC GAC GAC CGA
      V   G   A   F   V   L   S   W   V   P   F   N   T   F   R   L   V   G   V   V   Q   Q   L   L   A

1501 AGA GAC AGC AGC CCC AGC TGC GTG GCC CAC AAG GTG GCT CAA CTG GGA ATG GAA ATC ACC GCC CCT CTG GCT TTC
     TCT CTG TCG TCG GGG TCG ACG CAC CGG GTG TTC CAC CGA GTT GAC CCT TAC CTT TAG TGG CGG GGA GAC CGA AAG
      R   D   S   S   P   S   C   V   A   H   K   V   A   Q   L   G   M   E   I   T   A   P   L   A   F

1576 GCC AAT AGC TGC GCC AAC CCC TTC ATC TAC GCC CTG GCC GAC AGA AGC CTC AGA AGA GAT TCT GCC CGG TGC CTG
     CGG TTA TCG ACG CGG TTG GGG AAG TAG ATG CGG GAC CGG CTG TCT TCG GAG TCT TCT CTA AGA CGG GCC ACG GAC
      A   N   S   C   A   N   P   F   I   Y   A   L   A   D   R   S   L   R   R   D   S   A   R   C   L

1651 TGT CCT TGC CTG GTG AAG GCC CCA GTG TCC TGG GGA GCC ATC AGC TTC AGC AGC CAG CAG GGC GCC AGC TGG TCC
     ACA GGA ACG GAC CAC TTC CGG GGT CAC AGG ACC CCT CGG TAG TCG AAG TCG TCG GTC GTC CCG CGG TCG ACC AGG
      C   P   C   L   V   K   A   P   V   S   W   G   A   I   S   F   S   S   Q   Q   G   A   S   W   S

1726 GGC TGG CGC ACC AGA GAT AGA GCC GCT GGC GAG CCT GTG GGC CTG ACC ACC CTG CAG GAC TGA AAC CCG CTG ATC
     CCG ACC GCG TGG TCT CTA TCT CGG CGA CCG CTC GGA CAC CCG GAC TGG TGG GAC GTC CTG ACT TTG GGC GAC TAG
      G   W   R   T   R   D   R   A   A   G   E   P   V   G   L   T   T   L   Q   D   *

1801 AGC CTC GAC TGT GAA ACG GGG GAG GCT AAC TGA AAC
     TCG GAG CTG ACA CTT TGC CCC CTC CGA TTG ACT TTG
```

**Lo-GPR15-LgBiT in pTRE3G-BI coexpressing with SmBiT-ARRB2**

```
1    GAA CCG TCA GAT CGC CTG GAG AAT TCG GGG AGA CCC AAG CTG GCT AGC ATG ACC GCC ACC ACC GAC GCC GTG GTT
     CTT GGC AGT CTA GCG GAC CTC TTA AGC CCC TCT GGG TTC GAC CGA TCG TAC TGG CGG TGG TGG CTG CGG CAC CAA
                                                                  M   T   A   T   T   D   A   V   V

76   CTG GGC GCC CAG ATG AGC GGC CCT GCC GAG GCC ACA ACA AGC GGA CCT TGG CAG GAG GAA ACC ACA TAC GAC TAC
     GAC CCG CGG GTC TAC TCG CCG GGA CGG CTC CGG TGT TGT TCG CCT GGA ACC GTC CTC CTT TGG TGT ATG CTG ATG
      L   G   A   Q   M   S   G   P   A   E   A   T   T   S   G   P   W   Q   E   E   T   T   Y   D   Y

151  GAC GCC AAT GCC TCT GGA TGT GAA GTG CCT GTG CTG GCC GGA AGC GCC AGA GCC AGA GTG GGA CTG TAC TGC GCC
     CTG CGG TTA CGG AGA CCT ACA CTT CAC GGA CAC GAC CGG CCT TCG CGG TCT CGG TCT CAC CCT GAC ATG ACG CGG
      D   A   N   A   S   G   C   E   V   P   V   L   A   G   S   A   R   A   R   V   G   L   Y   C   A

226  TTC TTC GTG GCC GGC GTT GCT GGC AAC CTG GCA CTG CTC GCC GCC CTG TGG GCC CGT TGG CGG CGG AGA GGC TGG
     AAG AAG CAC CGG CCG CAA CGA CCG TTG GAC CGT GAC GAG CGG CGG GAC ACC CGG GCA ACC GCC GCC TCT CCG ACC
      F   F   V   A   G   V   A   G   N   L   A   L   L   A   A   L   W   A   R   W   R   R   R   G   W

301  GGC AGA CAG AGA GGC TGG CGG CCT TCT GAA ACC CTG GCC GCC AAC CTG GCT GCT GCT GAT CTG CTG TTC CTG ACA
     CCG TCT GTC TCT CCG ACC GCC GGA AGA CTT TGG GAC CGG CGG TTG GAC CGA CGA CGA CTA GAC GAC AAG GAC TGT
      G   R   Q   R   G   W   R   P   S   E   T   L   A   A   N   L   A   A   A   D   L   L   F   L   T

376  GCC CTT CCA TTT TGG ATC GAC TCT GAG CTG AGC GGC GGA ACA TGG AGA GCC GGC GAG CTG TCT TGT AAA GGC GCC
     CGG GAA GGT AAA ACC TAG CTG AGA CTC GAC TCG CCG CCT TGT ACC TCT CGG CCG CTC GAC AGA ACA TTT CCG CGG
      A   L   P   F   W   I   D   S   E   L   S   G   G   T   W   R   A   G   E   L   S   C   K   G   A

451  CCT TTC CTG GTG GCC CTG TCC ATG AAC GTG GGC GTG CTG ATG CTG ACA TTC GTG TCT GTG GAC CGG TAC CTG GCG
     GGA AAG GAC CAC CGG GAC AGG TAC TTG CAC CCG CAC GAC TAC GAC TGT AAG CAC AGA CAC CTG GCC ATG GAC CGC
      P   F   L   V   A   L   S   M   N   V   G   V   L   M   L   T   F   V   S   V   D   R   Y   L   A

526  GTC GTG AAC CCT AGC CTG TAT AGA CGG CTG AGC AGA GCT CTC TTC ACC GGC CTG GGC TGT CTG TCC GTG TGG CTG
     CAG CAC TTG GGA TCG GAC ATA TCT GCC GAC TCG TCT CGA GAG AAG TGG CCG GAC CCG ACA GAC AGG CAC ACC GAC
      V   V   N   P   S   L   Y   R   R   L   S   R   A   L   F   T   G   L   G   C   L   S   V   W   L

601  GTC AGC CCT CTG CTG GCC CTG CCC GTG CTG AGA GCC CGG GTG CTG CAA GGC GTG GAA CTG GAG GAT GGC ACC GAG
     CAG TCG GGA GAC GAC CGG GAC GGG CAC GAC TCT CGG GCC CAC GAC GTT CCG CAC CTT GAC CTC CTA CCG TGG CTC
      V   S   P   L   L   A   L   P   V   L   R   A   R   V   L   Q   G   V   E   L   E   D   G   T   E

676  GCC ATG TGG TGC CAG GAG GAG GAC GGC AGC TCC AGC CCC GGC CAG AGC CTG CTG CTG CTG CTG TTT AGC TTT TTC
     CGG TAC ACC ACG GTC CTC CTC CTG CCG TCG AGG TCG GGG CCG GTC TCG GAC GAC GAC GAC GAC AAA TCG AAA AAG
      A   M   W   C   Q   E   E   D   G   S   S   P   G   Q   S   L   L   L   L   L   F   S   F   F
```

```
751  CTG CCT CTG CTG CTG GTG CTG CTG TGC TAC TGC AGA ATT ACC AGA ACT CTG TGC CTG CAC GCC AGG CGG AGC TCT
     GAC GGA GAC GAC GAC CAC GAC GAC ACG ATG ACG TCT TAA TGG TCT TGA GAC ACG GAC GTG CGG TCC GCC TCG AGA
      L   P   L   L   L   V   L   L   C   Y   C   R   I   T   R   T   L   C   L   H   A   R   R   S   S

826  TCT CTT GAT ACA CAC CTG TGC CGG AGC TTC AAG ATC ATC TTC CTG GTG GTG GGC GCC TTC GTG CTG TCC TGG GTC
     AGA GAA CTA TGT GTG GAC ACG GCC TCG AAG TTC TAG TAG AAG GAC CAC CAC CCG CGG AAG CAC GAC AGG ACC CAG
      S   L   D   T   H   L   C   R   S   F   K   I   I   F   L   V   V   G   A   F   V   L   S   W   V

901  CCC TTC AAC ACC TTT AGA CTG GTG GGC GTG GTG CAG CAG CTG CTG GCT AGA GAC AGC AGC CCC AGC TGC GTG GCC
     GGG AAG TTG TGG AAA TCT GAC CAC CCG CAC CAC GTC GTC GAC GAC CGA TCT CTG TCG TCG GGG TCG ACG CAC CGG
      P   F   N   T   F   R   L   V   G   V   V   Q   Q   L   L   A   R   D   S   S   P   S   C   V   A

976  CAC AAG GTG GCT CAA CTG GGA ATG GAA ATC ACC GCC CCT CTG GCT TTC GCC AAT AGC TGC GCC AAC CCC TTC ATC
     GTG TTC CAC CGA GTT GAC CCT TAC CTT TAG TGG CGG GGA GAC CGA AAG CGG TTA TCG ACG CGG TTG GGG AAG TAG
      H   K   V   A   Q   L   G   M   E   I   T   A   P   L   A   F   A   N   S   C   A   N   P   F   I

1051 TAC GCC CTG GCC GAC AGA AGC CTC AGA AGA GAT TCT GCC CGG TGC CTG TGT CCT TGC CTG GTG AAG GCC CCA GTG
     ATG CGG GAC CGG CTG TCT TCG GAG TCT TCT CTA AGA CGG GCC ACG GAC ACA GGA ACG GAC CAC TTC CGG GGT CAC
      Y   A   L   A   D   R   S   L   R   R   D   S   A   R   C   L   C   P   C   L   V   K   A   P   V

1126 TCC TGG GGA GCC ATC AGC TTC AGC AGC CAG CAG GGC GCC AGC TGG TCC GGC TGG CGC ACC AGA GAT AGA GCC GCT
     AGG ACC CCT CGG TAG TCG AAG TCG TCG GTC GTC CCG CGG TCG ACC AGG CCG ACC GCG TGG TCT CTA TCT CGG CGA
      S   W   G   A   I   S   F   S   S   Q   Q   G   A   S   W   S   G   W   R   T   R   D   R   A   A

1201 GGC GAG CCT GTG GGC CTG ACC ACC CTG CAG GAC CCA CCG GTC GGT ACC GGC GGA GGG TCT AGC AGT GGC GGT GGG
     CCG CTC GGA CAC CCG GAC TGG TGG GAC GTC CTG GGT GGC CAG CCA TGG CCG CCT CCC AGA TCG TCA CCG CCA CCC
      G   E   P   V   G   L   T   T   L   Q   D   P   P   V   G   T   G   G   G   S   S   S   G   G   G

1276 ATG GTC TTC ACA CTC GAA GAT TTC GTT GGG GAC TGG GAA CAG ACA GCC GCC TAC AAC CTG GAC CAA GTC CTT GAA
     TAC CAG AAG TGT GAG CTT CTA AAG CAA CCC CTG ACC CTT GTC TGT CGG CGG ATG TTG GAC CTG GTT CAG GAA CTT
      M   V   F   T   L   E   D   F   V   G   D   W   E   Q   T   A   A   Y   N   L   D   Q   V   L   E

1351 CAG GGA GGT GTG TCC AGT TTG CTG CAG AAT CTC GCC GTG TCC GTA ACT CCG ATC CAA AGG ATT GTC CGG AGC GGT
     GTC CCT CCA CAC AGG TCA AAC GAC GTC TTA GAG CGG CAC AGG CAT TGA GGC TAG GTT TCC TAA CAG GCC TCG CCA
      Q   G   G   V   S   S   L   L   Q   N   L   A   V   S   V   T   P   I   Q   R   I   V   R   S   G

1426 GAA AAT GCC CTG AAG ATC GAC ATC CAT GTC ATC ATC CCG TAT GAA GGT CTG AGC GCC GAC CAA ATG GCC CAG ATC
     CTT TTA CGG GAC TTC TAG CTG TAG GTA CAG TAG TAG GGC ATA CTT CCA GAC TCG CGG CTG GTT TAC CGG GTC TAG
      E   N   A   L   K   I   D   I   H   V   I   I   P   Y   E   G   L   S   A   D   Q   M   A   Q   I

1501 GAA GAG GTG TTT AAG GTG GTG TAC CCT GTG GAT GAT CAT CAC TTT AAG GTG ATC CTG CCC TAT GGC ACA CTG GTA
     CTT CTC CAC AAA TTC CAC CAC ATG GGA CAC CTA CTA GTA GTG AAA TTC CAC TAG GAC GGG ATA CCG TGT GAC CAT
      E   E   V   F   K   V   V   Y   P   V   D   D   H   H   F   K   V   I   L   P   Y   G   T   L   V

1576 ATC GAC GGG GTT ACG CCG AAC ATG CTG AAC TAT TTC GGA CGG CCG TAT GAA GGC ATC GCC GTG TTC GAC GGC AAA
     TAG CTG CCC CAA TGC GGC TTG TAC GAC TTG ATA AAG CCT GCC GGC ATA CTT CCG TAG CGG CAC AAG CTG CCG TTT
      I   D   G   V   T   P   N   M   L   N   Y   F   G   R   P   Y   E   G   I   A   V   F   D   G   K

1651 AAG ATC ACT GTA ACA GGG ACC CTG TGG AAC GGC AAC AAA ATT ATC GAC GAG CGC CTG ATC ACC CCC GAC GGC TCC
     TTC TAG TGA CAT TGT CCC TGG GAC ACC TTG CCG TTG TTT TAA TAG CTG CTC GCG GAC TAG TGG GGG CTG CCG AGG
      K   I   T   V   T   G   T   L   W   N   G   N   K   I   I   D   E   R   L   I   T   P   D   G   S

1726 ATG CTG TTC CGA GTA ACC ATC AAC AGT TAA CAT ATG
     TAC GAC AAG GCT CAT TGG TAG TTG TCA ATT GTA TAC
      M   L   F   R   V   T   I   N   S   *
```

**Fig. S3.** The nucleotide sequence and amino acid sequences of Lo-GPR15 expression constructs. The amino acid sequence of Lo-GPR15 is shown in red, that of LgBiT in blue. The signal peptide of sLgBiT is shaded.

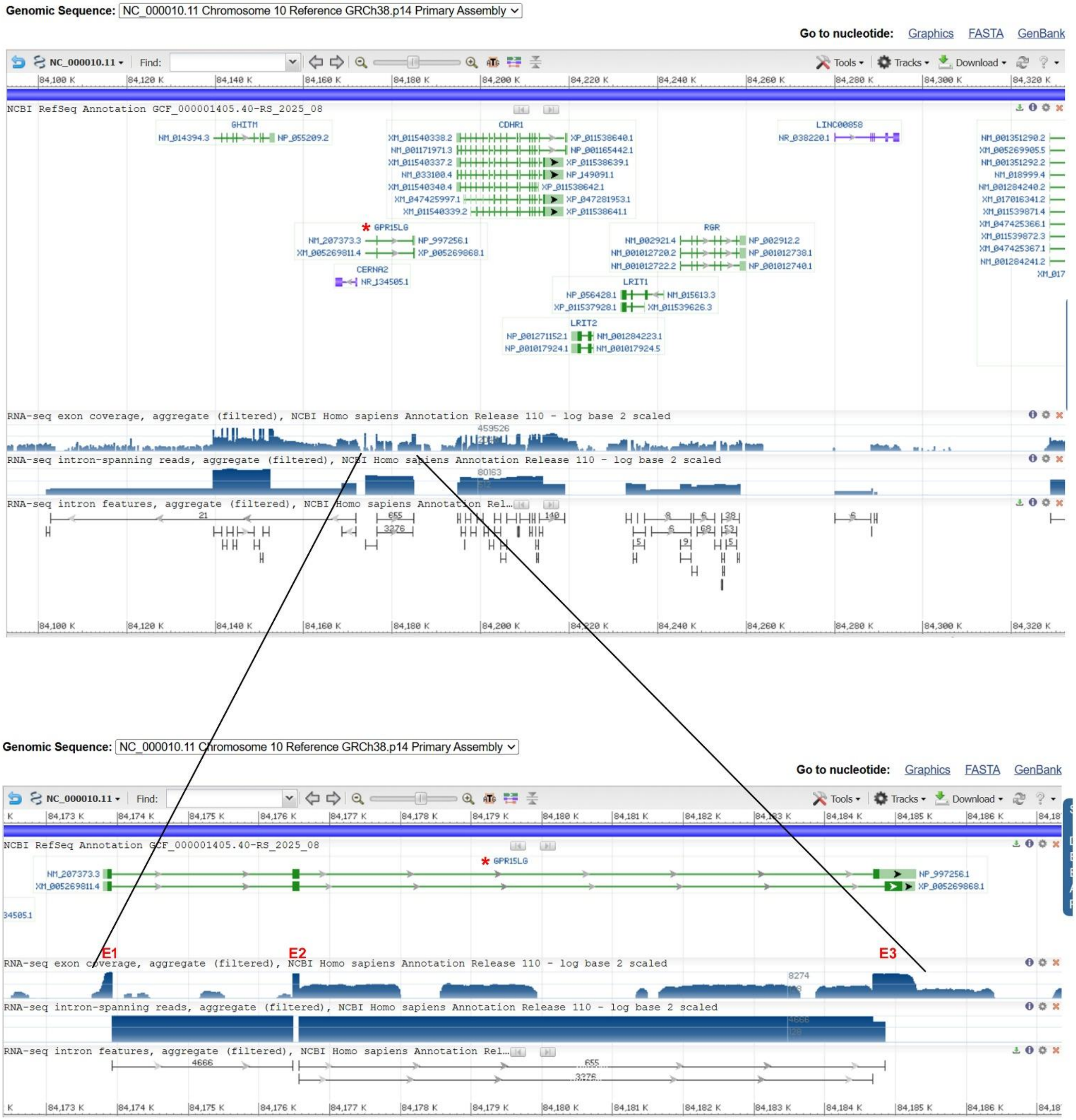


**Fig. S4.** Gene synteny and genomic architecture of human *GPR15LG* in human genome. Human *GPR15LG* gene is indicated by a red asterisk. The information is downloaded from the NCBI gene database (https://www.ncbi.nlm.nih.gov/gene/?term=Homo+sapiens+GPR15LG).

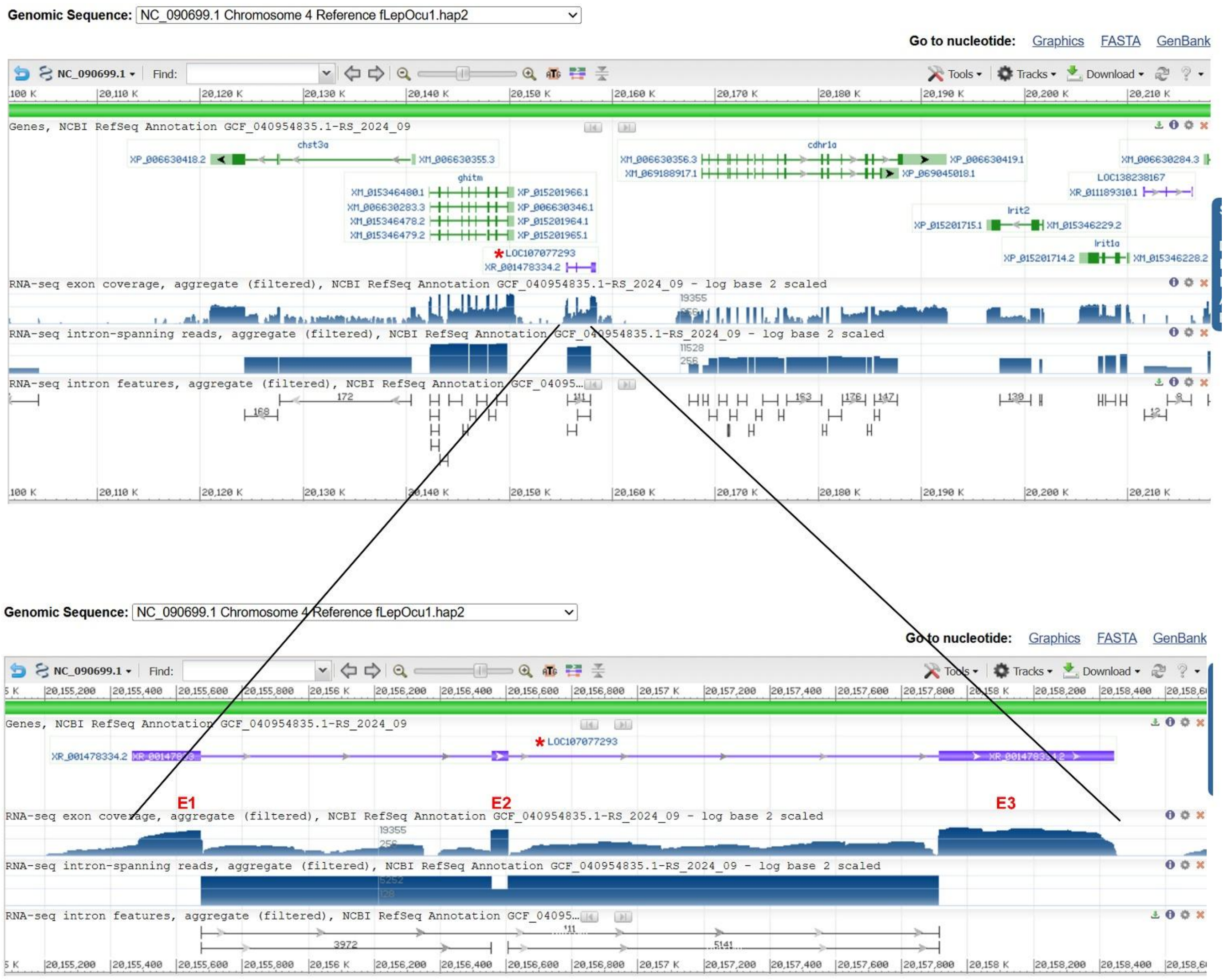


**Fig. S5.** Gene synteny and genomic architecture of the possible fish *gpr15lg* gene (LOC107077293) in the reference genome (fLepOcu1.hap2) of spotted gar (*Lepisosteus oculatus*). The possible spotted gar *gpr15lg* gene is indicated by a red asterisk. The information is downloaded from the NCBI gene database (https://www.ncbi.nlm.nih.gov/gene/?term=ghitm%2C+Lepisosteus+oculatus).

**(A) Possible spotted gar GPR15LG transcript (XR_001478334)**

```
1    CAT TGT GAG GTT GGT TCT TGG GGC AAT TAA CCA CCA CCT CCT TCT GTG AGA AAT CCA CAC AGC ACA CAG CGC AGA

76   CTG AAA CAG CTG AAG GAG GGC ATT GAG GCT CAG TTT CCA GCT CTC GAC ACA CCA GTT GAC AAA ATG AAG AGG GAG
                                                                                         M   K   R   E

151  ACA ATC CTT GTG TTG GGA CTG GTG TTA ATC CTC TCT ATG GCT GTC CTT TCA TCT GAA GGC AGA AAA CTC AAG TGT
      T   I   L   V   L   G   L   V   L   I   L   S   M   A   V   L   S   S   E   G   R   K   L   K   C

226  TGC AAA AAA TAT TTT CTG AAA CAT CAC AAG AAT GAC TCC CTC AGG CCC AAG AAT GCC AAG ACT GGC CAT CGC TGC
      C   K   K   Y   F   L   K   H   H   K   N   D   S   L   R   P   K   N   A   K   T   G   H   R   C

301  AGG CCA TGC AGA CCA AAC ATA CCT CTC CCT TCT TCG TAA TCA GCT TCT GCC ATG CTT GCG TGA CAA AGC TGG GAA
      R   P   C   R   P   N   I   P   L   P   S   S   *

376  GGC GTA TAC ACA CCA GCA TAA TAC TTT CAA GGG AAT AAA GCA TCT TGC CTA GTG ATT CAT CTT CTG AAT AAA AAA

451  AAA GTA GAT CTG ACG TTG ATG CTA TGC ACA GCC TTA ACT ATG CAC CAT ATA CTG TAT GTG TCT GGC TTT GAT AGC

526  TCA AAC TAA GAA AAG CTT TTG TAG TAT TCT TTA ATG CAA AAT TCA ACC CAC GGG AAA ATC GAA ATA TTT GTT ACC

601  CTC ACA TGC AAC TAA TGA TTT AGG TTT CTT GTT GTA ATG GGT AAT TTT TTA AAC AGC CCT TAT TAT TGA AAC TAT

676  TGT TTG ACA TTT CTA ACC TGC TTT TTT CTT ATT CTG TAT GAA AAC GGA ATA AAA ATA TAT TCT GCT ACA TGG ACA

751  CCT ATC TTG CTA CAT TAA AAT AAA AAA AAT ATT GTA CTT CT
```

**(B) Human GPR15LG transcript (NM_207373)**

```
1    ACT TCT GCA GCA CAG CTC CCT TCC CAG GAC GTG AAA ATC TGC CTT CTC ACC ATG AGG CTT CTA GTC CTT TCC AGC
                                                                         M   R   L   L   V   L   S   S

76   CTG CTC TGT ATC CTG CTT CTC TGC TTC TCC ATC TTC TCC ACA GAA GGG AAG AGG CGT CCT GCC AAG GCC TGG TCA
      L   L   C   I   L   L   L   C   F   S   I   F   S   T   E   G   K   R   R   P   A   K   A   W   S

151  GGC AGG AGA ACC AGG CTC TGC TGC CAC CGA GTC CCT AGC CCC AAC TCA ACA AAC CTG AAA GGA CAT CAT GTG AGG
      G   R   R   T   R   L   C   C   H   R   V   P   S   P   N   S   T   N   L   K   G   H   H   V   R

226  CTC TGT AAA CCA TGC AAG CTT GAG CCA GAG CCC CGC CTT TGG GTG GTG CCT GGG GCA CTC CCA CAG GTG TAG CAC
      L   C   K   P   C   K   L   E   P   E   P   R   L   W   V   V   P   G   A   L   P   Q   V   *

301  TCC CAA AGC AAG ACT CCA GAC AGC GGA GAA CCT CAT GCC TGG CAC CTG AGG TAC CCA GCA GCC TCC TGT CTC CCC

376  TTT CAG CCT TCA CAG CAG TGA GCT GCA ATG TTG GAG GGC TTC ATC TCG GGC TGC AAG GAC CCT GGG AAA GTT CCA

451  GAA CTC CAC GTC CTT GTC TCA ATT GTG CCA TCA ACT TTC AGA GCT ATC ATG AGC CAA CCT CAC CCC ACA GGG CCT

526  CAG TCG CCA CCA TGT GGG CCT CTC CAG TGC AAA CCA CCG AGC ATT CCA CCA TGA CCG GTC ACA GCT ACA AAT CCA

601  GAG ACC ATC AAT CCT GCT AGA GTG CAG GGT GGC AAG CAC CCA AGG GTG GCT GAC CAA GAC TGC AGA GTC TCC TCC

676  ATC TTC AGG TCC ATT CAG CCT CCT GGC ATT TAA CTA CCA GCA TCC AGT GGT CCC CAA GGA ATC CCT TCC TAG CCT

751  CCT GAC ATG AGT CTG CTG GAA AGA GCA TCC AAA CAA ACA AGT AAT AAA TAA ATA AAT AAA CTC AAT GCA GAC ACA
```

**Fig. S6.** The cDNA sequence and its encoded protein sequence of human or spotted gar GPR15LG. (A) Possible spotted gar GPR15LG transcript (XR_001478334). (B) Human GPR15LG transcript (NM_207373). The in-frame stop codons are shown in red. The signal peptide is shaded.

**(A) Possible GPR15LG transcript (XR_010805124) from *Amia ocellicauda***

```
1     CT ATA TCC ATC AAT CAG AAG TGC CCG GCT ACG CCA TAA ACG AAA ACT GAG TAG ATA ACT GTA AGT TGC CGT AAA

75    GTT CAG AGT TTT GCT GGC ACT CCA CTT GAC AAC ATG AGG AAG CAC ACT CTC TTA GCC CTG GGG CTG CTG CTG ATA
                                                   M   R   K   H   T   L   L   A   L   G   L   L   L   I

150   CTT TGT GTC GCT GTA CTT TCC TCT GAT GCA CGT AAA GCC AGA GCT GCT TGC AGG TGT TTT CTG AAA CAT CAT GTG
       L   C   V   A   V   L   S   S   D   A   R   K   A   R   A   A   C   R   C   F   L   K   H   H   V

225   AAA CAT CCT AAA GGA AAC TAC AGA CAC AAG AAG CTC TGC AAG AGT CCC TTT TCC TGC AGA AAA CAC TAC CCG AAC
       K   H   P   K   G   N   Y   R   H   K   K   L   C   K   S   P   F   S   C   R   K   H   Y   P   N

300   CTC CCG AAC ATC CCG ATC CCT TCT GAA TAA TCC GCT CCT CCA AGC TTG TCT CAC AGA GTT GGA AAT GGA TGG ACA
       L   P   N   I   P   I   P   S   E   *

375   ACA TTT AAG GCA GAA AAA CCC TTG CTT GAT GTT TAA TCC TGT TAA CTT GCA AAA TTA GAC TAC TGC TGA TGC AAT

450   TAA ATA CTG TAC AAC CAA ATG TCA AAA ATA TAC TTA ACT TAA AAA TAC TTA ACT TAA ACA TAA AAA ATA ATG TGC

525   CAT AGG GGA TGT CTT GTG ATC TTT ATT CTG ACC ACA ACA AAG ATG TAA TGT GTA CAG TGA TAA AGA TGA CAA TTT

600   TTG TGT ACT GGA TGA GAT TAT TCA AGC TCT TAA AAT GGC TTC TAC ATG AAC ATT TCA CAC ACC ATC ATG TTG CTT

675   TTT AAG TAG TAA ATG ATT ATA ACT GTA TCT TGG TAA TGT AAT GCC AGT CCA AGA GAC TCC TAA AGC TAT GCA ATG

750   CTG TGC TGT TAT GTT GCT TTG CTA GTT GCA TAA ACC TCT TTT TAA TGC ACT GAA TTA AAT ACA TCA GCA TGT TTA

825   ATA TC
```

**(B) Possible GPR15LG transcript (XR_010328695) *Anguilla rostrata***

```
1     CT TTG AAA AAG CAG GGG GAG GAT ACT GTG TGT TCA GAG GCG ACT GGG TGG CTC TCG GCA CTG AAA GAC GCC GGA

75    ACG TTC CCT GAA AGA TCA TTT TCC AGG AGT CTG TGA GAT ATG AAG GGA CAG ACG CTG CTG GCC TGG GGG ATG GTG
                                                       M   K   G   Q   T   L   L   A   W   G   M   V

150   CTC CTG CTC TGT GTG GCC TTG ATG TCC ACT GAA GCA AAA CAG AGC AGG ACT CAG CTC AAG TGC TGT ACT AGA AAG
       L   L   L   C   V   A   L   M   S   T   E   A   K   Q   S   R   T   Q   L   K   C   C   T   R   K

225   GAC CCG AAG GTT CGC AAA GAA ATC TAT AAA CAC AAA ATG CAA GGT TCG AGG TTC AAG CGT TCC TGC AAG ATA TGC
       D   P   K   V   R   K   E   I   Y   K   H   K   M   Q   G   S   R   F   K   R   S   C   K   I   C

300   AAA AAT CCA AAC AGG GTT TAC ATT AAC CCT GGA AAT CCA CTG CCC TCT CTC TGA TCC TCT CTG GAA CCT GGA GGC
       K   N   P   N   R   V   Y   I   N   P   G   N   P   L   P   S   L   *

375   AAG AAT TCT GGC TGA TTA ACC CTA CAC GTG CAC AAG ACA AAT TAT TTA CCT GGC TGG AGG CCA GTT CTG CGT TGG

450   TGT AAT TTT ATC AAT CAA ATA TTA ATA CAC ATT TTG TAA TAT TCT CTG CCA GGC AGT CTA ATA GAA AAA GTA AAG

525   ACC CCA ATA AAA ACT AGT AAA TGG TAG AAA AGG AAT AAA AAC ACT AAA TTG AGT GTT TAC ATG GCT GAA CCA TGT

600   AAT CTT GTG GTT CCT ACT TCA GGT TGG TCT GAT TAT AGA GAC AAA AAT GTG AAA GGT CCT CTG TAT TAT CAT TCA

675   ACA AAT GAT AAT CGT TCA TAT CTG TCC ATC TAT AAG CCA GCT GTA GAC CTA GGC CAT GTA GAA TAT CGA CCT GTT

750   TAA ATG TTC TAG CAA ATG TGT GTC AGT ACA TTC AAG CCT CAC TAT TTT ATT TCA TTT TAA AAT AAA ACC CCA TTT

825   TAT TTA CAT TTT TGT CAA TTT AAA AAG AAC CTG TAA CCT GAA GCT GAT GAT GAA TGC ATT TTC ACT TGT CAA GTT

900   CTA TCT ATA TTC AGT CTT ACC TTC TTT TTC TTC TTC TTC TAA TTA TTA TTA TTG TAT GAA AGT TTA TCT ACT CTT

975   ATT TGA TCA CAA TTG TAA AGA TGA GAA TAC AAT TTC TTA CAC TTT TAT ACT TTT TTT TAC ATG CTG TAA CAA TGT

1050  GAT AAT AAA CTA CTT GTA AAA GAT CA
```

**Fig. S7.** The cDNA sequence and its encoded protein sequence of possible fish GPR15LG orthologs from *Amia ocellicauda* (**A**) or *Anguilla rostrata* (**B**). The in-frame stop codons are shown in red. The signal peptide predicted by the SignalP6.0 algorithm is shaded.

### (A) Gene synteny and genomic architecture

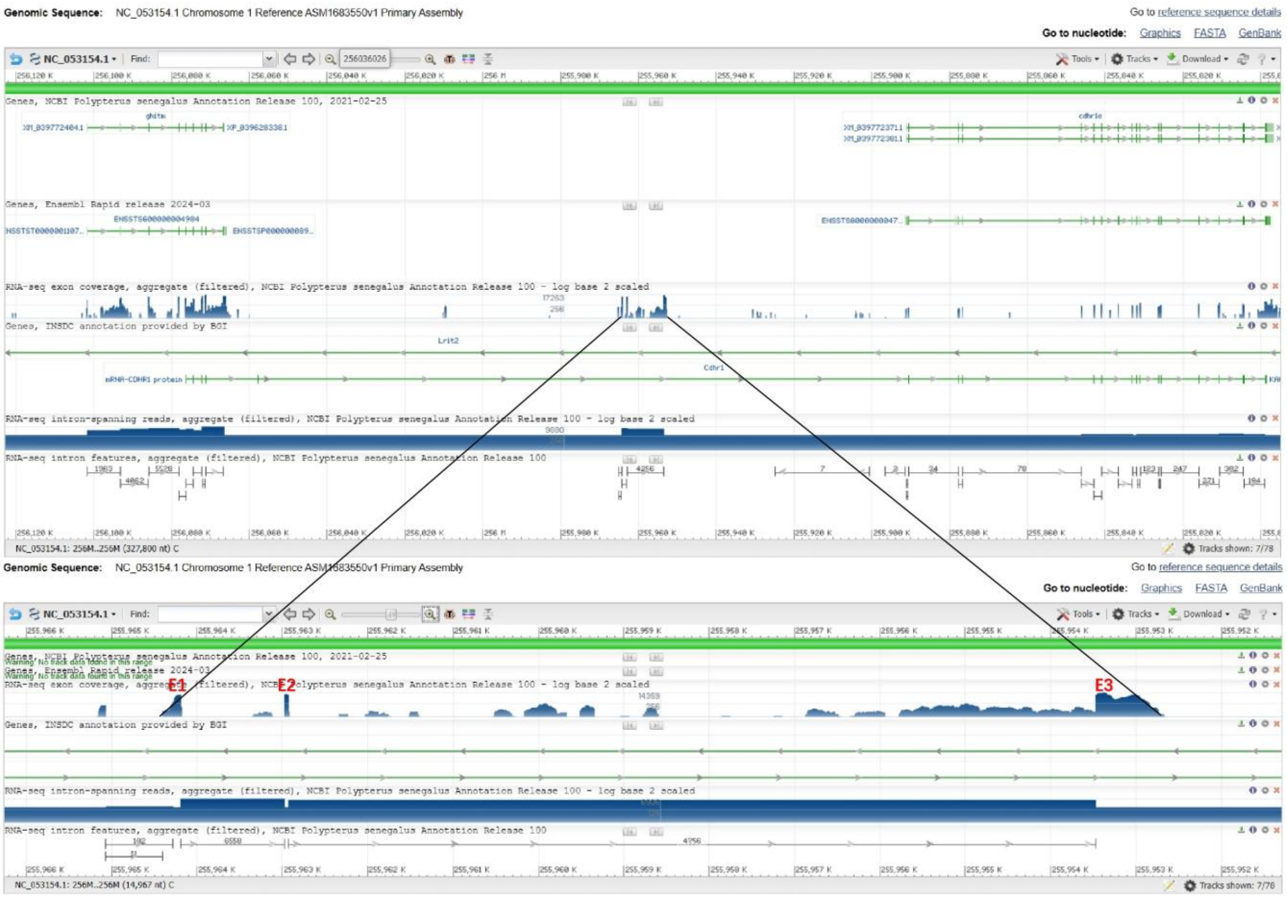


### (B) Genomic DNA sequence (exons highlighted in yellow, coding region underlined)

tgttttcttattgttttctgtgcgcttctacagtacatactgggtttaagtttcagagggctggccttaaaattgaaaaggtgaaccattgttgattttctccaatgcccctgtgcacacgtaagaacaaataaaagacagctcattcttcattcctgcagacctacagcaaccatttagaatgaaggcgca
cattttccttgtgtcagtgttggcactgatgatcacgatttctgaattttgctgtgaaggtaagaaaacttgactaagaaggaagctgattgattcataaagcattgcattaaactttgtgtaattatggaagttgaggcaggctttttatagcgtgtagtgaattgttaccttttattatacccaatagatctat
accattcatgaatttatatgcggtatatactaaaagacaggctcattagtgggacatgtttaatgatattttaacactgggaaaattaatccaagaatgaacacagcaacttaaagtgaaatcatgaaaatcatgttgttgaaaataaggcaaagtatatattgagttctattagcaaaaaactatttccta
aaactagataatttaataaaggaacatttatgttcagcatgggcaatgatgcattataaaacaaaaaaacactttaaactttgttaattatttactgagctaaagtgaaagtattgaaatctgagagttcttatagtacagtaaagcatttcacattaatttcacaggaaatctgcatagccttaacattgtgt
agtgtttaatttgaagacattttattctgttatatgcgtaaccctgataaacaattttaaaatactctgtaaaacaagcggtatctttaagttaaagccgttaatgttaattcatactgttcttaatgtaacatcccaatcaatattatatttatctctttaaatctaatgcattgaaagttttgttgcttttaactcagat
taatcaatgtgtcagcactctttgtatgcatcttatagataatgatcacacgttaaaatataaatatgtgtgagtagcttgtataggctgaccagacgtcctgatttgatcaaacagttcagtttcaggctccgcgtcccattttggctgaccagatgtcctgttttttttttccgagacaatcctgcctttatgt
atcgcgtcccggagtttaagaaactgttccgtttgtcccggttttaggtccccgaaagtagaagtggggagacttcagtttcatgagtattttggcaaggcgcgcgatactccatccgaggctttcattttatttgcgcccggcacactcgctgcattaatgagttgcgtccgtctggtcagcctgtac
acaattgtcccggtttgggactttaaaaaacctggtcaccttaatcttaaaagaattgcaacttatatttgcaattgtagtgctattctacttctgcttttcattttaggaaggagaatcagatgctgccccagacgtaagcatggtgtacaaggtaacccataaaacatttcctaattattttactcactactt
tcaacataagaatttttatgggctttaattgtattccaaaaaattgcagatggtttgtcatcaaatgtgggagactgaaactatgattcatgataattaactaattctgtttgaaacaattaatctaaggttatgcaccacaattgattaacttccttgatttggaagcaatattgtcataaaattttaacactatg
catgtgatgtcttttaaaaaaaataaaatagttccagacaatgtgtttatatttctttgataaatttggctttagatatattgtcaataacagtacacagagcaaaatatctgctggtattcttgcaaaaacagttttagctatacaaataaaattacaaatttggtttcagtcagcaaatttctttatttgaaaata
aagctgttttgatactaagactatgtcgctatattgtgcacaattgaaaggtttattcatataaaatatgtgtgtgcacagcatcatccccagtccaattgcttgaccaaaaaaggtattagtaaatgaaaaaggttaaagatagaatttttgaaattaatttgaaactatccgaatcattgtggaagatatg
aaaacttcttcaaacataaattattttttaaaggatcttttcatttgtcaatttattatagcctggacaccataacaatagtttaaagcagagaaatgtcattaaagtttttgaagagtttttttgtgttaacaataacttaattcagaattacaaaaatgaatgaacatacaaatatttaaagagcaatggatgta
aaagtagaaaaaaatacttatttttttaaagaatgaaacctatactctatcacagccagggggcgctgtaaagcaagcaattgaactgcagcaaatgcaagccttcaaaagtaaatctgagaatgtaaatgcaaagatacagttgaacacttttgatgggaacaggccattgactctccaaaataa
catcaacttcacatctgaaagtcccaaaagcccatctttctaacacactaaaaggttatttattcaaactctcaaatatttctgtgaaattattccttaaagttccaactgtgtactgtacaactatttttaatgaacaatttattttaaattagcagtggggatccactataataattccaatataagtttaatact
tcaatcatgccacctattaatctcttaacaataaataaatgcatagactttaaagaaaaaaagttgctggttcgtgatgcatgactccgagaaggtgtattaacccatctgttattcatgtgagaaaacagtatatgcaaaacaaatgagttctgcaattataaagtgcaatgtaaaataaattatcacctta
aagtgcaataaaaaaaattgtgacattgaaagacattcatatttgctccagggggtgttgtacaatggctgaccctgggcttggaaatcaataaagtatatccatccatccatccattttctaacccgctgaatccgaatagggtcacgggggtctgctggagccaatcccagccaacacagggca
caaggcaggaatcaatcctgggcagggtgccaacccaccgcaggacacacacaaacacacccacccaccaagcacacactagggccaatttagaatcaccaatccacctaaccggcatgtctttggattgtgggaggaaaccggagcgcccggaggaaacccacgcagacacggga
gaacatgaaactccacgcagggaggacccgggaagcgaacccgggtctcctaactgcgaggcaataaagtatatcaaagcaaaaaaaataaatacattttgaaccccatgtaattcaatcccaccagagacttgtgtaggtattttagcctaggtttataaatctctccatgactgtttattgaaat
ggatgatgtacataaatatgtacatgttgcatgcttttgcactgttctatttgtatttgtgtatttgtaaatctaaatttaaaaatgcatatagaaacatgcatcaatagtcaaatgtgaaagaacgttgtttgaacataacaacatgagggatggcagttgacaacaaaagagagaaaaattaaagctcct
gataacagcatttaacaataaaatatgaagctatatatattgtggtagaaatttaaaatcttattaaagaaataaatgtcttctaacaggacaaatcagttatcaggcaggagaaaagctggtcattgtatcttttaattactcctcggtaaaatgccttagagggagcattctttaaaagcagtccgttac
agaatggtcagagaaacaaagaacagaggttcattttataaactgttgaatttacattcagtgtcaatcaaggcatacatttattctggttggttgattggtttacactaaaatgatttatattaaataaaagtctaatatataatattctggttcttctcataccttttgcgatgagacaccacccttgaaattt
ctgtgcatataacaactgtccattctcatttataggacatctgctgatttcttagtcatccttcagtaaattatattacatcaacccttcttctggtacatttacttgaccatctcagtatttttcctaaaccagttgttgtatttcagtgtacattttgttgttcacttgacctttatctggtttcctggtgtgcctggac
agttttctgtcatttattaaccctttatgctttttctttaagatgaatgctatgttaccctaaaattattattccacaatgtaaatataaaaacaaaagactaaacaaatcagtctaatgcaatcgccatattattcaaaatatcaaccataataaaaggataactgtctgtatgtttgtctgtgtggcagtctggtt
gctatgtctttggcattccaaaagatgtcacatcacaaacatttttagtaataaaggatctttcaatgaatatctgagaaaaggaaggcagccatccatagaatacaccctagcagcatatgcacaaagctttcaattattaaacttaaaatctttcatcaacaacaacaatatttatttatatagcaaattt
tcatgcaaatgatgtagctcaaagtgctttgcatgatgaagaaagagaaaaaagacaaagtaagaattaaaacaagagaacactaattaacatagtaataacagtaaggtctgatgcccagggaggacagaagcaacaaaaaactccagacggctggagaaaaaataaaatctgcaggggt
accagaccattagactgcccagtcccctctaggcattctacctaacatcaatgatcagtcctcattgtattcaaggttctcatggaaggacttgatgatgatggtcacgtggacttctggactttaatccatcaatgtattaacatcatggtgctttgattaggtggtggtggcgcaggtcatcaccaca
gcaaaccgaaaaaagatcagaagagagagtacggattttagagccaccatgaatagtaatgataattaagtgaatatgcagagcaccaggattaaattaaaataagttatgagaaagccatgttaaagtaatgtggtttttgcagtttttttaaagtgctccaccgtatcagcctggcgaattcctatt
ggcaggctattccagattttaggtacaataacagcagaaggccgcctcaccacttcttttaagtttagctcttggaattctaagcagacactcatttgaggatctaaggttacgatttggaatataaggtgtcagccattccgatatataaaatggagtaagattatttaaggctttgtaaaccataagca
gtgttttaaagtcaattctgaatgacataggtaagcagtgtagtgagttcaaaactggagaaatgtgctctgattttcttttcctagttaggattctagcagctgcattctgcactagttgcagtcaatttatgactttttgggtagtcctgagaggagtgcgttacagtagtctagtcgactgaaaacaaa
agcgtaaactaatttctcagcatctttcaatgatataagaggtctaatttttgctatgtgtcttaagtgaaaaaatgctgtcctagtggtctgattaatatgctatttaaaattcaggtcacagtcaacatcgatcacatttgtcttgtttaatattgtccaaaatttatccagagcaagagccaacctgtgaac
gttgcaatcgaacttcagcctcactgggccacatattttgggtgtgcaccaaattaacaacattttggacaaaaatcagcctccgtgtcacaatcacccctaagccattaacagctgtgtttggtgaactccaaaatgggcttaaagtgaagaagaataaattgattgtaattgcctattccacacttct
agcacgtagacttattttgctcaattggaagaaacccagcccactttttataagtcagtgggtaactgatgtcctatactatttgaaattggagaaaatcaaattctcgcttagaggatctcgtcaaaagtttttaaaacatccattatccaacccactatatccgaactacaggggtctgctggagcca
accccagccaacacagggcacaagccggaaacaaaccccagcccacacacaccaagcacaatttagaatcgtcaatgcacctaacctgcatgtctgtgggaggaaaccagagtacccgaaggacaacccatgcagacacagagataacatgcaaactccacgcagggaggacccgg
gaagcgaacctgggcctcctaactgcgaggcagcagcactaccactgtgccaccgtgccacccatgcaatgttattttcttcaaataacataaaaaatgcattgcagttgttatttttaataaatggcacatcacaaacattaacacagctgttattaattccataccaaatggtatataacaaagaca

tacagtatgtggacatcaacttatcttaggccgggtttatacttcacatgaccatacgaatgctccagtggacactactgttacgcatagtgttctactatttatacttgcacgcatactttacataaatctggaagattccatcaggtggcagtgcgagatatcatcacagtgagaaaacgttcggcttc
actgtgttgtgaattgcctgaaatatccattaaattccaaggacaccttgccacaatatctctgaaaaggatgtttaatgattacatccatcaatctagagatgtgcccattccagcaagcagaatccctggatggggcatcagcttatcacaacatgaacacaagcactcacatacactagcgtcat
tttagcgtcaccacatccccaaaccttcatgtctttggaaggaaaccggagtacactgtggaaactcaccagtaaaacatgcaaacaccaggcagggaacatcagggatgtgactccctgcgagacagcagcactaccactctgccacaatgtcgtgcccacatgtatttgttaacagtattca
ttatttaaatggagttaatgatttatctgtaaaatgtaatatacatactttaatgcatttcatcctgaaagtgatatcaagtataaatcaaaggattctaaatatgcagagagctgaaatatcataagtgtaatgtgttctgagtggcgattgctgcttgtcactgctgtcaggtcaggaggacgccccaga
agtaagtaggaattaacaactgggttggttttaagatgacgtttacaacggtctacttgaatgataaaataaactacgaggttaaagtggacatttcaagactaaagccgaaatttccactttaatcacaaaatagaccttttcactgtgtccttaatttaaaatttctgtggctcaaatacagcgctgtac
attctgttgctgttgtgaagatgcaaaaaaaaaaaagacggcacagaagatggtatgtgagacttttaaaatatattgtgtcattacgatgggaaatatgcgacacttaaatataaaagcaccacaaatgcatttgtatgttgccattttgcttcaccacatcgaacatttctcaaacatcgaggcaggt
acatcaatcctgtaggatcctcaaagcggctttctgtcacatgtagatagtaaacagagactctgacgtcacattctgacttgatcacagtgcgccccctccctctcccgtctttttgctggtactgcaactcactcacatgtcacattaatttctgaggagctgctcaaaggacgcatcaaataaac
gctgggaacgcgtggcagccatgatgcgtgggtgtatgcgttctgagtatgaagtatgaactagcccttatatgtttagcatggctactagctaatataagtttacttataattctgaaatgtgccatttttcaaataacttgaccatgtaccatacagaaactatgtaatacaataacattcaggagagt
tcttgcattaatggtgcattagtaaggacagttttaacaaaaaaactgtgctatttactttcacaacattaggtcttcttgttggtaatgaaattaaatattcagatgccagtttagcatttccaatctgaaaataaactcacctactgtactgttgtaatcctgtatcttttctgaaagatagtttaacatatactg
tatgtagctaactataccagcaatatccatagtttaatccagtagcaaacttagttaatccaaatttgaggctgtgtttggtataagacatggaccaaccatgttaaacaaaaatcacacggatatatcccatatttcaacatactagtaattcaaaaataaatgggagctgattttgtttggagtggcattt
gaattctggcttgggtactgcctatctggaatctgcgcattctccttttgtttgtattgtttttctccaggttttctttccacacctcagatacttgtttattggtttaattggccacatatagatgtgttggactggcatctgtccaatgttagctcttaacttgaactcattgtttccaagataggttttagttcctg
tgatttgaataaaaagtctagaaaacacatgaaataacattttatataaacaccattatatgtaaaagtcattgcagttctgaaattgtgttttgggagtgccacaattcaaaacttgcttttaaggttatagaataatttaggctaatgttcatggaatcctgcacgttcattcatccttaactaccatctacc
aagttcatttttattgcaaggtcaacatcagaatctaaatcctatttcgctcttcatcttttggaacatttgttgtaggtgttgcctgattgcaggaaaactaaaggaaatcttttttagaagaaagttatgttatctttggatcaagaacataaaagttttaattaagatcaatatgttgaatcactaggcccta
taatagactgacacctgtttcctgctttgcatcccattactttaacttggattaaatggatttgcaaatgttaaattgaaagaaataagtacaaaataatcaaggtttaaattttaaagaactagtgatgaactttgaagggcttttgacattttgttacagagagaaagggaagttaccaaccaaaaccc
aatattttctaccagtaagatatcttggaatacatttatgggccattccagactggaatgtgagttaagtgaagcttattaatcaacagagaacggtccattgtgctaaaagctgcactaacagacagtaaaataaaaaaatgaacaaaaaaaaaaaagaataaaaaccgtaatatgctaccttcaaa
agataattaactatattgttttaatgctgtgccatttacaaagcccagagcttttgagagtattagaattatatagggatccccgagtatttaggaaaatcgcacaatttgccaaaaggaaagtggactgataatttatcattttaccaggatcaacagccatttttttgcttttccttcatatactgcaaaagt
acccacacgggaaaggtactgtgaagaagtcaaagttttgctttcattttcaatagagttacatgtttctggcatccctgatgatgtaaatatatgggaatgctaactgaaaaatgaaacacatcactcactttgacttaaactttattagcattgcatcactaggagcataatgtttttgagcaaacatta
cctgccaacgatagataatataataaatgtaacaatatgtgctgtctcttgccctacatgtacatttagctcctttttctcagtatccgcatgtgtctggacatctcccgttcggtgctccttctgtcaatcgcattactgtaaagcgtgcttctcaggctctctttattttcaaggtgccttataaatctcctgtc
caaggaagccactcttcatcatgtggcttctacacctttattcctcctcttgtttttcactcactcgcacttcctcttcgattgaatcaccaaagtcaaattgaccaatcagattgctctgacggacaggacacgctcatgcagattttagcattttagtgtaaagtagatatttatatatatttttttaaacatga
agaatgacaggtcaagagttgagacatctcccacgtgaaccccttttttacaccataaattagaattaattgataaataaatacccactgccatgattcaaagtctcttgtattaaggttcatatgtacagagaatctgctggaaggagtgggacttctggggctacatgacaaatgaggagagccat
cttcttggttgccgtgtagagaagtctataggggatgtgttgtgatcactatttcaacttgaattgagtgtacttacaggattaaagctttcctgaatataatttgagatatgtatttctgcctatctgtcaagatatgagtctgcggatgcagtaactcaaaaaaggaccattcactttaaataaaactggat
accattattaaacgtttcaggttggtttttcaacaaaatctcttcagtacattttgctctatacagacaatgatattatgatatatgtatatttgtgccacaactactacacaactaatgaatatgaatcgatatttcattgcacatgataaaatactgatgtacatgatgttattgctaaaatgtctctgtaagca
ggagtgctttggatgtttttactgcagggtaactcttttgtttgcagtgctaaggtacgcaccgggaatgactactcaaataatagttaaaaattcaaatcccttttcaagagatttgtttaaatacatcaagtgatggaaagaaactgtaaacaaagtgactaaaccctaataacaactgataaattctc
tttattcccacagtgaatggccccaagttccggaataagctgcactgctggatctgctcaaatatgaagggggcaccagctaaagcaaatggccatcctgatatacccataccaactttttaagcactttgtagctccagcgctgtaccaccatcaaacagaaatgacataaactgatctgtacaat
cctttcacttttctaacctctgcaagtaaccaaatgaatacagcaagcatgacattcatttataaggaaactgctttagttattgtctttgtattcttttatcattttatatatttgagaaagtagtctgcaaatttgcttaatcagaaatgtgtaaaaaatgtctaaaaatgaggagagaagacagactataaa
gcatgcaggatggtatctcagatgaggattcatgagagtctttcttttagtaactgtaattgaaattaattgtaaagttctagggatatggagcttggaccaatgctaacgagtgtgtaaattggacacatgaatggatgaatgagaggatttttgtgagggtgcatttccatactgcatacaaaacata
cacaaaaatgtatatataaagacttgctgttacttgtagaaatgtttaaacttctgtggttttaataaaaaaggtatgcaaataagtgtttgactttttagaaatatctgttcatatgtattttatactccctctttctttgtctttccatttttttccatatactgtagatatgtctccatcccctcattttaatgtataaag
ctttgcagttttcatgtgcttaacatgcttaaaacaattgaagaagatggtaacaaaaggcataaaaaaagcaaaacatctcttt

#### (C) cDNA sequence (coding region highlighted in yellow)

tacatactgggtttaagtttcagagggctggccttaaaattgaaaaggtgaaccattgttgattttctccaatgcccctgtgcacacgtaagaacaaataaaagacagctcattcttcattcctgcagacctacagcaaccatttagaatgaaggcgcacattttccttgtgtcagtgttggcactgatg
atcacgatttctgaattttgctgtgaaggaaggagaatcagatgctgccccagacgtaagcatggtgtacaagtgaatggccccaagttccggaataagctgcactgctggatctgctcaaatatgaagggggcaccagctaaagcaaatggccatcctgatatacccataccaactttttaagc
actttgtagctccagcgctgtaccaccatcaaacagaaatgacataaactgatctgtacaatcctttcacttttctaacctctgcaagtaaccaaatgaatacagcaagcatgacattcatttataaggaaactgctttagttattgtctttgtattcttttatcattttatatatttgagaaagtagtctgcaaa
tttgcttaatcagaaatgtgtaaaaaatgtctaaaaatgaggagagaagacagactataaagcatgcaggatggtatctcagatgaggattcatgagagtctttcttttagtaactgtaattgaaattaattgtaaagttctagggatatggagcttggaccaatgctaacgagtgtgtaaattggaca
catgaatggatgaatgagaggatttttgtgagggtgcatttccatactgcatacaaaacatacacaaaaatgtatatataaagacttgctgttacttgtagaaatgtttaaacttctgtggttttaataaaaaaggtatgcaaataagtgtttgactttttagaaatatctgttcatatgtattttatactccctct
ttctttgtctttccatttttttccatatactgtagatatgtctccatcccctcattttaatgtataaagctttgcagttttcatgtgcttaacatgcttaaaacaattgaagaag

#### (D) Amino acid sequence (signal peptide shaded)

MKAHIFLVSVLALMITISEFCCEGRRIRCCPRRKHGVQVNGPKFRNKLHCWICSNMKGAPAKANGHPDIPIPTF

**Fig. S8.** Information about possible GPR15LG from *Polypterus senegalus* (gray bichir). The information is downloaded from the NCBI gene database (https://www.ncbi.nlm.nih.gov/gene/?term=ghitm%2C+Polypterus+senegalus).

### (A) Gene synteny and genomic architecture

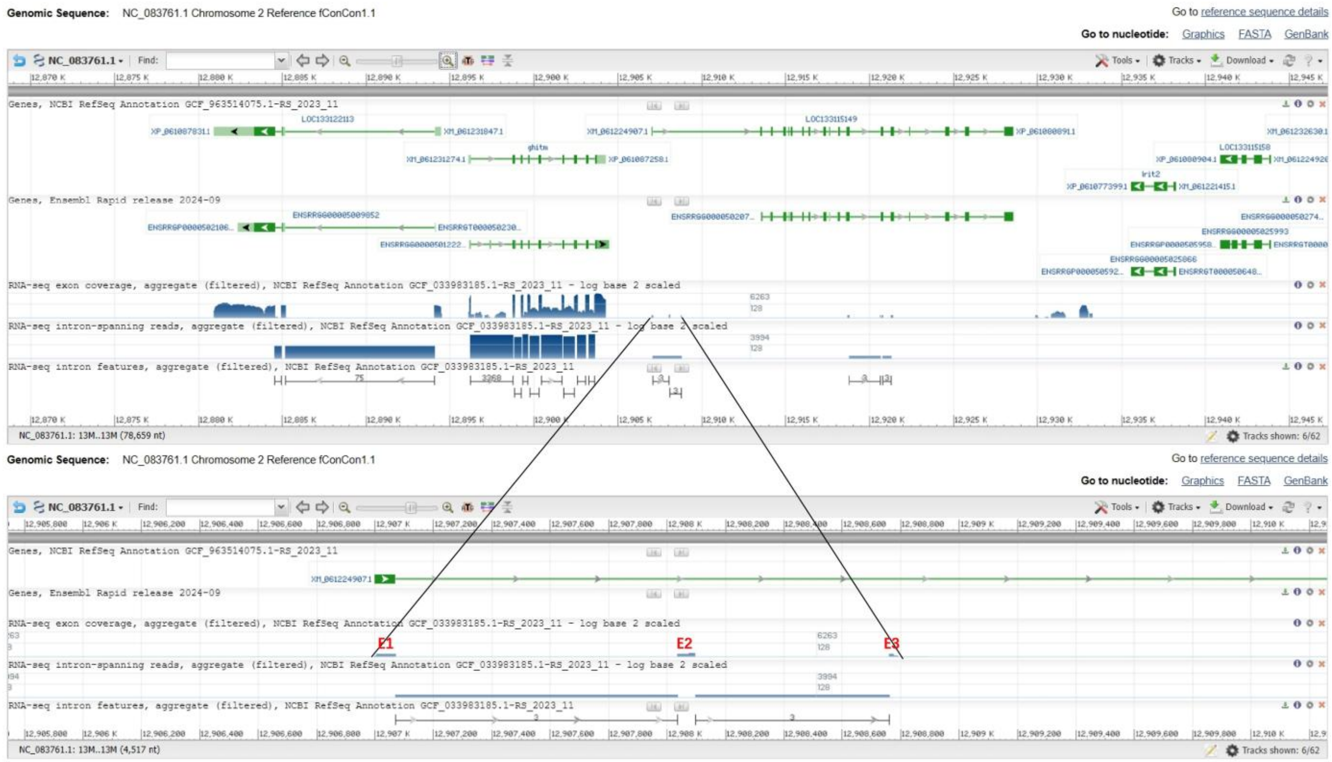


### (B) Genomic DNA sequence (exons highlighted in yellow, coding region underlined)

ccttaaagattattttccagaagcctgcgacatgaagggacagactctgctggcctgggggattgtgttcctgctttgtgtggccctgatgtccactgaaggtaaggaacgagccatttccccctacacagatacaagctgctggaacagagaaaataaatacagtcatgttgggtattttcccaaaggagttcatcgttgactatagatattggtatgtgttcattagcttatgaatagctgttcaaatatatacaggtctaaatatatacagtgacttcagaaagtattccgaccccttcaatttttgtgcacttaattttgtgatttcattttaaatggaatttttttaaatttctgcccatcaatcgacactcaataacccataatgacaaaagttaaaaagtttttaagaaatttttgcacatgaaaaattattctaaatcaaagaaaggcattaagggcagtttatccattcttccgggcagattctctcaagctctgtcagagtggatgggtagcatcttcaggtctctccatggatgttctatggggtctaggctttggccccattcatgtacaaaggctttggctgggtcactgaaggacagacagccactccagtgttgtcttggctgtatgttctgctgatttcagcttgtagttatgggttattgagtgtagactggaaaaaatgttttcatccattcaaaattaaatctacaacacaataaagtgtggaaaaagtggaggggtctgaagccactgtatacgtttacatgtataattgcatgatacagcattttatttaatgtggtgggtattgctatttggctgaaatataatgggtattttagtttttcttcatgggagctacataccattatataacatccatgggtataatgtcaatggatttgatattagctttatttgttctgtttagcttgtttccatgcagaatatccactgtataatatgatcatttagcaactttagaaaacaacaaataaaacattttgatagctctggaggtgaattatctcatcttaatttttttttttttagcaaatcacaagaggattcagctcaggtgctgtactagacgggacctgaaaattctcaaaggtaataaagatgaatatatttcctcctaacactgcctcaataggcatttaaggtgctgactggcatgtctttacaatgaatgtgtaagttgatacagcctcctctacaaatataagtcttctactgtatctttgccatcaaaaactataaagtataaaaactgcaatttcctttgaattatgggtaaacatttttttaaattccccacaattttgttaatgtggggagagtgctgtcgcagcatactgtaccgaggttgaatcccagccaagtggagtgtgcatgttccccacacgtacgctagggtttcctcccatggtgcaaagacatgcaggttaggagaactgtccctggggtatgaatgtgtgcggtggtactgtgtgtggggctccgcagtggattattgacctgaccaggatgccttcctgcctttcataataattcaaataatgaatcaaattagaccatcaaattcaaatttcaaatgaaaatgtcagatttctttacatttactacattatttcttgccctttgttatggaaacaaaatctaatttggatgcaaaattttagattttatatggccatttatcaaatgcttttctgagtactataacatgcaaatagaaatgtgttactaaccaatcactgttgctttcccaacgcagaactctataagcataaagttaataggatcaagcgtttctgcaagaagtgcaagaaatcacgaagatttgtcctcaatccactcccttctgtctgatgctcttctgagcatagaggcgggaatatcggctgattaa

### (C) cDNA sequence (coding region highlighted in yellow)

ccttaaagattattttccagaagcctgcgacatgaagggacagactctgctggcctgggggattgtgttcctgctttgtgtggccctgatgtccactgaagcaaatcacaagaggattcagctcaggtgctgtactagacgggacctgaaaattctcaaagaactctataagcataaagttaatag
gatcaagcgtttctgcaagaagtgcaagaaatcacgaagatttgtcctcaatccactcccttctgtctgatgctcttctgagcatagaggcgggaatatcggctgattaa

### (D) Amino acid sequence (signal peptide shaded)

MKGQTLLAWGIVFLLCVALMSTEANHKRIQLRCCTRRDLKILKELYKHKVNRIKRFCKKCKKSRRFVLNPLPSV

**Fig. S9.** Information about possible GPR15LG from *Conger conger* (European conger). The information is downloaded from the NCBI gene database (https://www.ncbi.nlm.nih.gov/gene/?term=ghitm%2C+Conger+conger).

#### (A) Gene synteny and genomic architecture

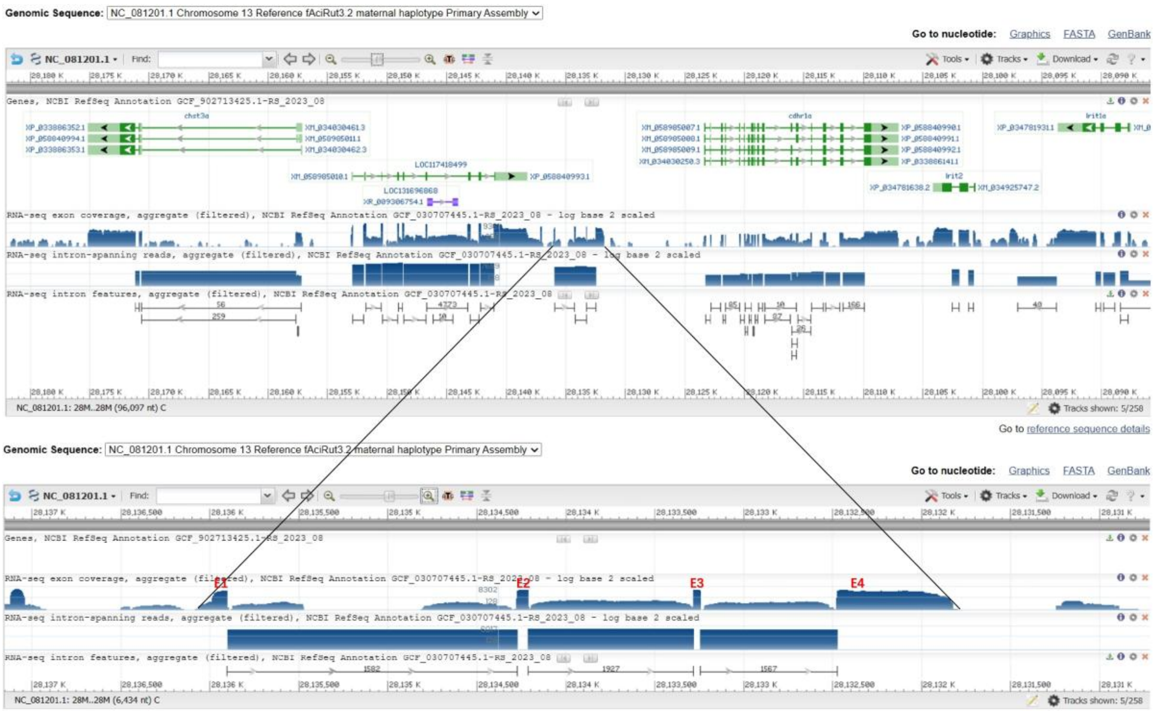

#### (B) Genomic DNA sequence (exons highlighted in yellow, coding region underlined)

ttaaaataatattctcagataacagtgtaatctgtcctgtatgtactgtgagtcaccttgtgaagggtagaagggtacaagagggttgggtcttaagaaaactttatacagcataaataaaaggtggcacacaggttctatttttcagtaccctaaacaatccagcagaaatgaggaagttgacaattg
tgggcttggtgcttgtattaatgctgtgtttggccgtactttccactgaaggtaagaacattacaattgaactttttaaatttttaattatagtaaagtgctgcatgttacagtatatattacaactaaaaggattgtattgtgataaccacattgcagtaatacattagtatacaagaacacatattctttttaga
aatgttcttgtatggatgtttcacctttcaatgagagacaagaggtttgttggattgatttgaaaataaatgtatgtttcagaacaaaataccacagctgcacagtgttatatcaatgcggtcttccaaaacattgcaaacatatctgctggctaaaaacagatttaataaaatccttccattaagatgcca
aggtcaaatgatccaaatgggatttgtcagtattctaataaaatgcatggtgatattatttctttacgagtgagactgataaattgaggccaatatatatatatatatatatatatatatatatatatatatatatatatatatatatgcatttagcaatatgcatgtagcgtgtaccataaatattttctttgctgg
ttttgtacacaattcagcctccaagttacatatagccttcatgttctggatgacaaaaaactgttgataattttttttgttactttgataacagagctaaatctttcaaccatatttttattataaaaaaaaaaaaaataatattctcatcaacttttgtttgtgcaatttctagtgcaatttataaacataggtatgttttct
atctttttctacctaagcaggatacatattctgacctgtatcacttagcaatatattgtttatttcaattttctcatcgttcagatgcaatgaccaattgcttaatgacatttcaaatttgatgatctaaatttaccatattctggttatatatatatatatatatatatatatatatatatatatatatatatatatatata
tatatatacacagtatatatatatatatatatatataagaaaaaagaaaaagagaaaaggatatttcaggtatatatatatatatatatatatacacacacacacacacacacaatacaatgtgtgcagtacctctgtagttgagaaaatattttgttggttgtcttgttcaggcatgcttctcaataaagt
aattttccatgttaagattagggactatttgctttgcactttagtcatggtgatgatcaataatctattgacgtttatgaaactgaaactttttttgttcaaacactgttcacaaatgatcatgaaaaatcacaattagatagcaataccagtgatggccaacagtaaaatgactgtattatgtcatatgcaaaa
ctgtacagttagcttatgcatgcatacaaatagactccaaattaactctgtaaattcaactttatctatagagtaagtgggtaacactttacattgagctgtgtctaacgacattgtaattacactgtaattggattgtaaatatacatgtgtaaatacaatgtacttcaaaagataccaactaccacaatatt
actttttgtagttaaacatattattaaatgtaacaatacaaccatttgttgaattaatgtgttactacagtgtaaacatatttttctccttttaaggaaagaaaataaaagacaggagatgctgtataaaatacaaacaccatcaacaagttaacaaaggtgtgacataaattactttttttctttgacatttcat
tagtttttgttttaattattattattagttgccttgcttttatttatactggcttcttcccagacattcaaatgaaaagcgttccctttctgctgcctaatgtcacagttttcaaatatgtcggacaacaaaattacagaagattacattgatttggctgtttgtccaagtccatttccttgaaatgtttccagtttaca
aataaaaccttccagctaatgagtaatacaaatccaggtgacattcactgactcatgtactgtaaataccacttcatcccaaaatgtacacatagtttaagagtaagtggtttcagctatcatgttcttacctttgatcctgactgtttttattgcaattactcaattgaatgagttcagaaaaagataaccg
acgtatgacctacataatgtgggttagtgttgtagtctaggtctgcagctctgtgtctaaccctaaagcactactctccagctttcacttaattactgtcccttcacactcccccaaatactggccagcctcctctatttataccgctaagccagaccaccattgaaatatatttttccaataattcagtttta
tgattttaccttttttcattttacaacaatacatttcctctaaaatgcacatatcaaattacacataacaaggaaacaaaccttcaaaaaaaataaatgaaccccttctacaggctagctagagccacattgccttgtgggtaacaagacaatgtggccaccccttgcaacacacacaaacaaaagcat
tttcctttttacaatggtcattttatctcctttgatattgaatttcttcattgatgtctgcacccgttcattattatgtcacaaattagcactgtaccctttttattttttcagttccaggcactgtgaagaattccttcaacaaaaacccaggtaagcacaaaaggaaatgtgatccaaagtggcagaatggcctata
cagtgcttcccattatactgctatagttgggagccatagttaagcgactgtgctgtttgttttccaccttataatgagaatactagtgttagtgaaatggcattatggggcaaatgggagccatgactgtactgccttataacctgttccgcagtataattgcttaactgatataaatggctcagtaagttta
tcaggttgatggctgctgccatcccgtatctccctttgtaaaccagccaaaaatttaaaaataccaaaccaacaagatttaggtgacaataaactgcatgagtacagcaatctccactatctatagaactgaagaatgtaatatattagttaaccaaatattaaaatacttgtttgcttttcccaggtttttg
aatgtattcacatgaattacattcattgttaaatacaaaagcatgtgaaataaaacacattctggatttcccccatgactgagcttttattgtttgcatataaagaaaagtgtatataaatataactttataggagtgccagtttttacatgcatacccttttcctttttaactgccattgttcctaacaataattgaa
tactctctgttatgtggtttagatgaagggtgggtggtctatatcgtttattagattcctgtagaatgttacgcataggatgaagtagctgatgatactaaagacaaactcagaatcatatttgtttcccttccacagtggtcagcaaatacaagaagcgttgcattgtgtggtgtccagtccggccggac
ttaccgctccctcactaatgcctaaaataaagcaggctttctgtatcatttatcttgaaaatcaactcttttgcaaataaatgaacttttttctttttttataatggaataaccaaaagcatctagctatgttatgtgttcttgcatgcagcataacaggttaagataaaaaaatacataatatgataatagcatataa
tgagcgcatatcacatgtattgtcataaatcatataattaaacattatatcattataattgcagcttaatttgctttcatagtgtgtgcttaaaaagagttttatttacaacaattacagctgagaaatgctttcaaaataaagaatgggcattcatctaacaaatgaaaagaactacaaaacaacattcacac
aaagaatcatttaaagaaaagaaagaaagaaaaacaacaacctgggcttacaaagtggtatttacattcagtggtagaaacaagattccttctgtcctgagaatacatactgtactggaaataattttataatgttactgttttgattataaaaccttataaactatataaaaatggtactgttttcatcaac
gatttgtctcctgtatttcttcataatgatttgttttatttatttatttatttttttgtgatgaaatgcagca

#### (C) cDNA sequence (coding region highlighted in yellow)

ataacagtgtaatctgtcctgtatgtactgtgagtcaccttgtgaagggtagaagggtacaagagggttgggtcttaagaaaactttatacagcataaataaaaggtggcacacaggttctatttttcagtaccctaaacaatccagcagaaatgaggaagttgacaattgtgggcttggtgcttgta
ttaatgctgtgtttggccgtactttccactgaaggaaagaaaataaaagacaggagatgctgtataaaatacaaacaccatcaacaagttaacaaagttccaggcactgtgaagaattccttcaacaaaaacccagtggtcagcaaatacaagaagcgttgcattgtgtggtgtccagtccggcc
ggacttaccgctccctcactaatgcctaaaataaagcaggctttctgtatcatttatcttgaaaatcaactcttttgcaaataaatgaactttttctttttttataatggaataaccaaaagcatctagctatgttatgtgttcttgcatgcagcataacaggttaagataaaaaaatacataatatgataatagca
tataatgagcgcatatcacatgtattgtcataaatcatataattaaacattatatcattataattgcagcttaatttgctttcatagtgtgtgcttaaaaagagttttatttacaacaattacagctgagaaatgctttcaaaataaagaatgggcattcatctaacaaatgaaaag

#### (D) Amino acid sequence (signal peptide shaded)

MRKLTIVGLVLVLMLCLAVLSTEGKKIKDRRCCIKYKHHQQVNKVPGTVKNSFNKNPVVSKYKKRCIVWCPVRPDLPLPH

**Fig. S10.** Information about possible GPR15LG (isoform a) from *Acipenser ruthenus* (sterlet). The information is downloaded from the NCBI gene database (https://www.ncbi.nlm.nih.gov/gene/117418499).

### (A) Gene synteny and genomic architecture

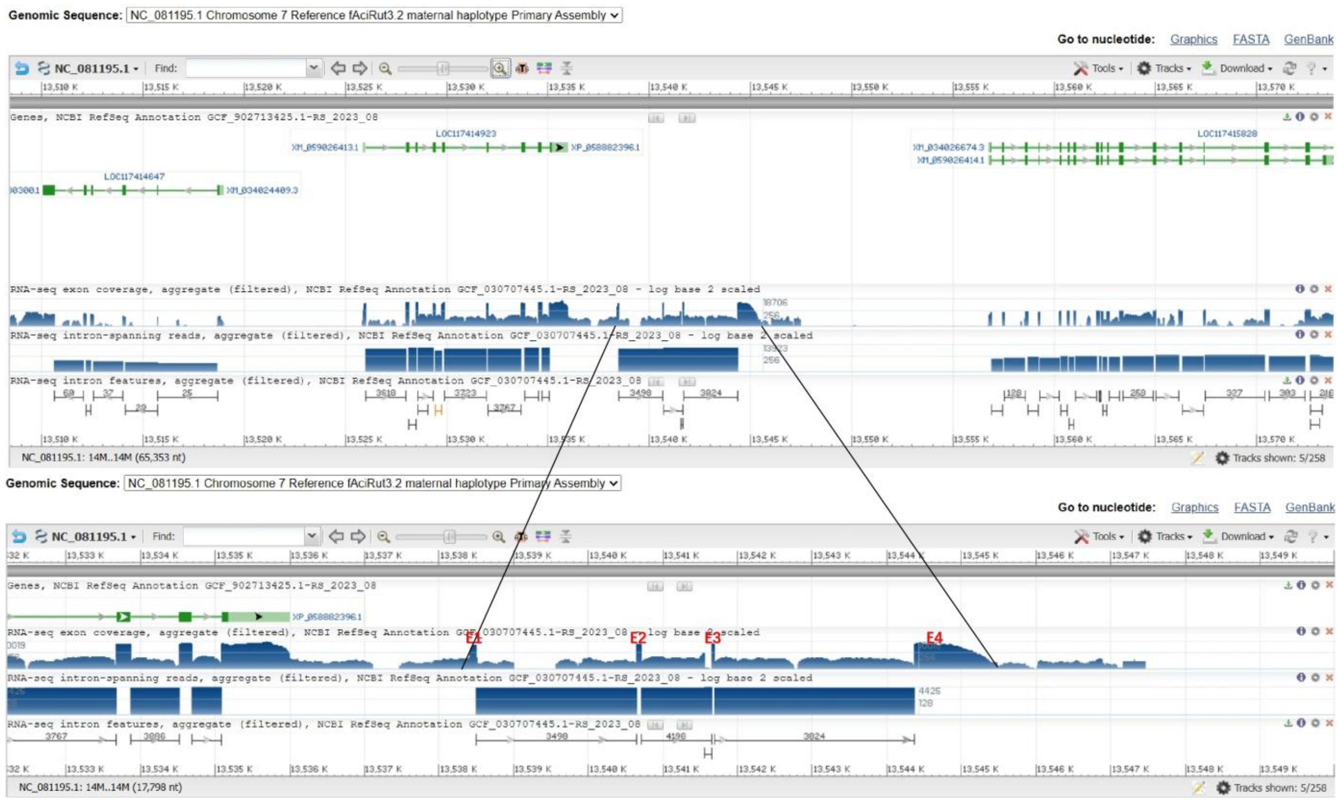


### (B) Genomic DNA sequence (exons highlighted in yellow, coding region underlined)

cttgtgaagggtagaaggtacaagagggttgggcttacgaaatctttatacagcataaataaacggtggcacacaggttctatttttcagtatcctaaacaatccagcagcaatatgaggaaggtgacaattgtgggcttggtgcttgtgttaatgctttgtttctccgtactttccactgaaggtaag
aacattagaattgatttaactattaatatttgtatttttttttttttttaattaagatataacctacaatggtatgataaaagcaaaagtactgagaaactggtaattcctatataggcaattttctatataagtgacaagcctttaataactatacatttagcttcacaaacaatagacaattatgcctttatgttgt
atgttgaaacatagggctatttaaagagaaaaggctgtttttccactggattttttttattacataaatcagctctgttgtaacctgtatgtggtgtgcaatgcccttgtccacaacactgtctatttctcatagtaaagtgtgtttgttttacgtatgttacagtatatattacacctaaggattgtattgtgataa
ccacattgtggttaatacattagtatgcaagcacacacatcatttttataaatgttattgtatggatgttgcacttttcaatgagagactcttgagtgggttgttgcattgatttgaaaataaatgtactgtatgtttcagaacaaaataccccagtttcataatgttatatcaacgcggtcttccaaaaccttga
aaacatacctgccggttaataacagatttaataacaccctgccaatgagatgccaaggtcaaatgatccaaatgagatttctgaatattctaatgattgtgatgttctttctttacgaatgagtctgctaaattgaggccaataatatatatatatatatatatatatatatatatatatatatatatatatatatat
atatatatatatataaattaaactaattaagtttaaattaagtgtttttatgcacaaaaggtttaaatgattagaacatatactgtgtatatatatatatatatatatatatatatatatatatatatatatatatatatatatatatataggacatttccgacaaaagctgaagaagaaattagaattagaaacaataa
cactgaatgacaataaatgtcagaaatattttttttacaataacatatactaagacacaattcatacacaggcataaccgctaataacagatccctcaacaaaatattacaattgtgtgtttgtactacacaaaaataaatcaatacataaacgtgtgcagtgccactgtagttgagtaaacattattttg
gtgtcttgttcaggcatgcttctgaatggagtaattttccatgctcaggtttgggacttagctttgcacctcattcatggcagatgatcaataatttgttgatgtttatggaactgaaaattaaactttttttgttttgttttacagatgacataaattattattattattattattattattattattattattattatt
tatttatttattagcagacgcccttatccagggtgacttacaatcgtaagcaaatacatttcaagtgttacaatacaagtaatacaataagagcaagaaatacaataacttttgttcaagcaaagtacaagtgtgacaaaccacaattcaataatacagcagataatagtgatagttacatcaggatatg
attaaatacaaaatactacaggttaaacacttggcagattacagtaatctgaagtacaggattaaatgcagtaaaataggggggcagataagagcaaaataaaaaaaaagttccaattctgttaacacttcacaaatgatcatgaaaaatcacaattagatatcaatactagcgatgaaagaccaaca
gtaaaattactgtattgtgacatatacaatactgtacaattagcttatgctagcatacaaatagtctccaaattaactctgtaaattcaactttatagagtaagtgggtaacactttacattgagctgtgtctaactacattataattacactgtaattgtatatgtaattacacagtaactcgattgtaaatgca
catgtgtaaatacaatgtacttacaaaagataccaactaccacaatattactttttttgtaattaaacatgttattaaatttcacaacatacctatttgttgaattagtgtgttcctacagtggaaatgtgagacacttagtatgcaataaacattatgtttctctttttaaggaaagagaagaaaacacaggag
atgctgtataaaatataaacatcatcaagaagttaacaaaggtaggacataaattactttttctttgacattctcactgtttttattagtattagttgtgctgtttttttatttatactggcttcttcccagtcattcgagtgaaaacctgttcctttctcctgcctaatgtcacggttttcaaatatgtcacattgtgt
gggattcagacaacaaaactacgattacattgatttggacctttgtccaagtccatttccttgcaatgtttccactttgcaaataaaaccttccagctaattattgatacaattccaagggatagtcactgactcatgtacagtaaatataatttcagccaaaaatatacaaatagtttaagagtaagtggtt
tcagataacatgttcttaccttttgatgtgtgcattctttctgagtgtttttttattgcaattactcaactgaatgagttcagaaaaagatacccgacgtagacctacataatgtgggttagtgttgtagtctgcgcccgcagctctgtggctaactctaaagcactactcttcagcttttacttcattactgtcc
cttcacactcccccaaatactggccagcctcctctatttataccgctaagccagaccccccattgaaatattttttccagtaattctgtttttatgatttgaccttttttcattgtataacaatacatgtcctctaaaaagcacatatcaaattacacataacaaggaaacaaacattcaaaaaaataaatgaaa
ccctttccacaggctagttagagccacattgccttgttttacccacaaggcaatgtgcccaccccctgcaacacacacaaacaaaagcattttccttttacagtggtacgtttatctccttttatattgagttacttcattaatgtctgcactcgttcactattgtgtcacaaattagcactgtaccattttattt
ttcaggtccagacaccatgatgaattcctcaaacaaaaacccaggtaagtacaaaaatgaaaattgatccaaagtggcagaactgcctattagtcatagagaaacactgtggcggatctaccttcgggcaattttttaatatatacaaaatcgtcattttgtgtgcatttcacaatgaaaatatgcaca
gaccaggtgagtttcataagggataagacacagtggcacaaagtctaaaaatatgtatgtaagattcaaagcgacaagacaacgtattaactacattgcaaatcagatgttgcaaacaggtgtactgtccttagatgtctttttaaataaatacctacaaataaaatacatcaatcttcggcactaaa
atgacttcattctattcgacacgtttaataggtctatccacacattatacattgtgctggtctctaacgtaacacagatcagacttgaaaccatgttacagacagacacccatgctaatcgggaaagttccacattttgtacgttcgttgtgtagacgcattgggaaacaaagacacatgtttaaaggca
gtgcactactacttagcatttggtagcaacggttacttattaattttgggttggtagagttttacatttatttatgttctctctacctgtctttaaagctgtaatttgcatcgtttaaccatgggctgaattaaagcgagcgcaaatattttgtagttctagtacattccttaatgaaaactggtggcttcctactca
ctgtaaggaggatatagtcactacactacacacctttgatttcgtaaaatacagatattctgtctggatacacacggcttgttaaacttaacaaataccaaaatgtagaattaacaaagaaacgacttgaagtattaattatctgtgtttgcattaaaaggggcgctatttaaattgcaatctggtgataca
ataaacctgttagcactcgagctagtagcattgcttttatctgcgagtaattgttctttagagtgaaagaaagattgaagtcaatatcgctgtaaattgatttgtttagatttagtttttatttgaatcgttttatctcttagtagtttcacaccccccccccccccccccccccccccctccccccccctcctttat
aaacgtcttgcatccctccctgtcttccttctatatgcgtcttgcatttctttttttacatttgggaactcccccattacattgcggtcaattacaaaatattcaactcaaaatgtaacaatcaacaatggtgaattgaaaaactattttctactggaagctaaaaatgaaaatgtcagtttgtcatgccgtggca
agaaacacagttatacatccattaaaacagacgttagcgtaagcttaaccttttaatctactttccttagcgattggacgtctcttctgagttcttaccgaaaacataaaatgtcactttctacttgaatccataccaatacatttttaaattccaaatacagtaatagtatgtgcctactaatggaaattatcg
tgcctcactggcctgggctgttgcagtaacagttattaaagggcatagggaataataataataataataataataataataataataataataataataataataataataataataataataataaacacgacagagaagacagcaccactaaaaatacagtttagagtatgatttctttggtttaattaaatt
aaaataagtggcatacagaacgtggactggttctttctactagtaaaacgtggttctggaagttagagattaatgtttaatgttttgccatcccacattgttaaacacacagttatcatagcgacaccagctgctaggtcaggacacatttgtaagtaaacgctgcgtctcgcatgacagtgcttggtg
ctgatactgtcatttataggttggaacaatgaacatcgccctcgcatagcaatgcttagtgctaattggttgaaacactaaatgacagcacgcatacggcaatactaattggttgaaacgttgattaacgactttcttgtaggcagtgcaaatgctcattcataaactaaagttaacgaaaaaagttgct
ggtgcggccgcacccccagcacccccactggccgtgtcccatcacgcaagtggtggaagccagccaagagctcaataataccaaaccaacaagatgtaggtgaaagtaaacgccatgcttacagcaatctccacaatctatagtactgtagaatataatgtattagtttaaaccaaatactga
aatgtatgtttgctttcgaatgtattcacatttcttacattcattgttatttggatttttttacaccctttttcttttaactgtcgttgttcctaagaataattgaatacactattaaaggaacattaatatacagtattgtaatcctcttgcatgacttacatttagtagtttactgagcatatataaataaatatatctctatc
tttcctccctctctctctctctctctctctctctctctttctctccctctctctctctctctctctctctctctctctgatgtgatttagattaatgatggatggtcttctatatagcttattagattcctgtagtatgttaataaacatatctatgaaaataatagctgatgatactaaagacaaactcagaagc
atatttgtttcccttccacagaggtcggcaaatccaagaatatgaaactgcagcgttgcaggatgtggtgtccagtccggcctgacttaccgctccctcactaatggctaaaataaagcacactttatatctttcatcttgaaaactactcttttgcaaatagatacactttttcttttttataatggaataac
caaaagcatctagctattttatgtgttttgcatgcagcataactggctaagataatatataatatggtaatatttttaatagcatataatgagcgcatatcacatgcattgccataaatcatatacattaaacattatacagcttaatttgctttcatattgtgagtttaaaaagtaaaaatctaagttgagatgta
gtaatattctatattctatcttacatattgcgctatcttggaaagggaacactataattaaaatgtccttctattctgagtgatgtgttttatttttctcaatacaagaactacagccaagaaatgctttcaaaataaagaatgggcattcatctaaccaatgaaaagagctcagaaaaaaaaaactgtgcttac
aaagtggtatttacattcagtggtagaaacaagattcattctgtcctaagaatacatactggaaatcattttattatgttactgtttcgattataaaacctaataaactatatcaaatagtacattgtttccatcaatgatttgtcgcttgtatttcttcttaatgattctttcatgatgaaatgcagcaatactaata
ggaaccactatcaaaaaagcatgttttttaaagaacatttaataaaacctactactcataatatattatatgcatctcgtttttttgtcttgtaatgtcttgctcttcaccactgctctaatccagttaaagatatgatgtactactctgaaactgctatttattgaattatatttacatcagagacaaatccatacct
atctcagtgattatacttatttttggaaatgtatgcattttaataaaaattacaaaaacaattggtgccaaattcagttctgtacctgatccctatttagatatggacttgagttattataaataggattgctttctaccataag

**(C) cDNA sequence (coding region highlighted in yellow)**

gtacaagagggttgggcttacgaaatctttatacagcataaataaacggtggcacacaggttctatttttcagtatcctaaacaatccagcagcaatatgaggaaggtgacaattgtgggcttggtgcttgtgttaatgctttgtttctccgtactttccactgaaggaaagagaagaaaacacagga
gatgctgtataaaatataaacatcatcaagaagttaacaaaggtccagacaccatgatgaattcctcaaacaaaaacccagaggtcggcaaatccaagaatatgaaactgcagcgttgcaggatgtggtgtccagtccggcctgacttaccgctccctcactaatggctaaaataaagcacact
ttatatctttcatcttgaaaactactcttttgcaaatagatacactttttctttttataatggaataaccaaaagcatctag

**(D) Amino acid sequence (signal peptide shaded)**

MRKVTIVGLVLVLMLCFSVLSTEGKRRKHRRCCIKYKHHQEVNKGPDTMMNSSNKNPEVGKSKNMKLQRCRMWCPVRPDLPLP
H

**Fig. S11.** Information about possible GPR15LG (isoform b) from *Acipenser ruthenus* (sterlet). The information is downloaded from the NCBI gene database (https://www.ncbi.nlm.nih.gov/gene/117414923).

### (A) Gene synteny and genomic architecture

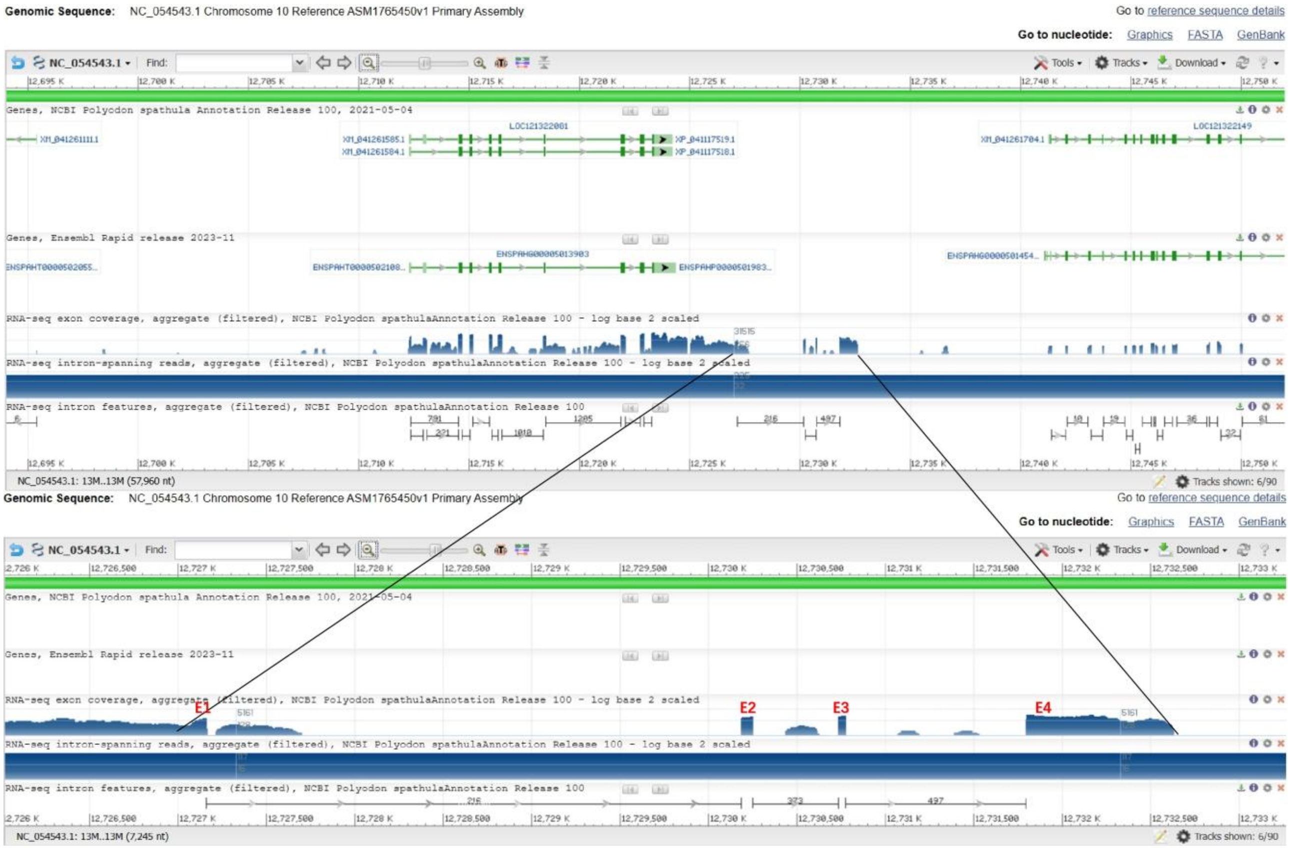


### (B) Genomic DNA sequence (exons highlighted in yellow, coding region underlined)

aaactcactggttcttttaaaattaatattatcagggaagtgtcttcggtaacagtgtaagctggcaatctgcactgtatgtattgtgagtcatcagtgaagggtacaagagggttgggtcttaagaaacctttatatagcttaagtaaaaggtggcacacaggttctatttttcagtatcaaacatacca
gttgcgatatgaggaagttgactattgtgggcttggtgcttgtgttaatgctttgtttctctgtactttccactgaaggtaagaacattagaattgatttacctttttttttttttaaattaagatataacctacaacattatgataaaatcaaaagtactgagactataaattcatatattggcagtatattgtatata
agtgacaagcctttaattatacatttagcttcacaattagacaattatgcgtgtatgttgaaccacaggtctatttaagagaaaaggcactttttccactttttttttttttttgggggggttacactaaacgattgcccttgtccaaaaacatctttctcatagtaaagtgttgcatgttacagtatatattacaa
ctaaggactgtattgggataaccacattgtagtaatacattggtatgcaagcacacaggtttttatgcatgttcttgtatgaatgtttcactttttaatgaaagactcttgagctggttgttgtattgatttgaaaataaatgtactgtatgtttcacaacaaaatagcagcatttcataatgttatatcaatgcg
gtcttccaaagccttgaaaatatatctgcggctaataacagatttagtaataacctgccaatgagatgccaaggtcaaatgagtcaagtgagatttgtcaatacacaataaaatgtattgtgatgttctttcgttacaaatgagacaagtatgttttctacctaagtaggacctgttctgaactgtatcaaa
tgcaatgacaaattgcataatgacatttcaaattatatgtttaatctcaatttatcttattgtggttagtgtgtgtatatatatatatatatatatatatatatatatatatatatatatatatatatatatatataatccaaagatccaaaatggaaaattacctatatggtaacagggtcctgggagacagtcaa
catggttttaggaaagggagatcgtgtctaactaacttgcttgatttttttgaggatgcaacatcgataatggataattgcaaagcatatgacatggtttatttagatttccagaaagcttttgacaaagtcccgcacaaaagattaattctcaaactgaacgcagttgggattcaaggaaacacatgta
catggattagggagtggttaacatgtagaaaacagaaagtactgattagaggaaaaacctcagaatggagtgtggtaaccagcggtgtaccacagggatcagtattaggtcctctgctattcctaatctacattaatgatttagattctggtatagtaagcaaacttgttaaatttgcagacgacaca
aaagtaggaggagtggcaaacactgttgcagcagcaaaggtcattcaaaatgatctagacaagattcagaactgggcagacacatggtaaatgacatttaatagagaaaagtgtaaggtactgcacgcaggaaataaaaatgtacattataaatatcatatgggagatactgaaattgaagaa
ggaatctatgaaaaagacctaggagtttttgttgactcagaaatgtcttcatctagacaatgtggggaagctataaaaaaggctaacaagatgctcggatacattgtgaaaagtgttgaatttaaatcaagggaagtaatgttaaaactgtacaatgcactagtaagacctcatcttgaatattgtgtg
cagttctggtcacctcgctataaaaaagatattgctgctctagaaagagtgcaaagaagagcgaccagaattattccgggcttaaaaggcatgtcatatgcagacaggctaaaagaattgaatctgttcagtcttgaacaaagaagactacgtggcgacctaattcaagcattcaaaattctaaaa
ggtattgacagtgtcgacccaagggactttttcagcctgaaaaaagaaacaaggaccaggggtcacaaatggagtttagacaaaggggcattcagaacagaaaataggagacactttttacacagagaattgtgagggtctggaatcaactccccagtaatgttgttgaagctgacaccctg
ggatccttcaagaagctgcttgatgagattttgggatcaataagctactaacaaccaaacgagcaagatgggccgaatggcctcctctcgtttgtaaactttcttatgttcttttatatatatatatatatatatatatatatatatatatatatatatatatatatatatagtatatatatataagatataatctttctc
accagcagattggtcgtcaatatttaagatacccgacatgtcttcatatacattttcttaatttaatgtggtcctctctgaattttacagtgtagacttgatatactgctgatgatttagaagcatgttattaacagttatatatattatatatatatatatatatatatatatatatatatatatatatatatatatgta
tattagaaacaataacactgaatgacaataaatgtcagatatgtttttcacaataacatatactaagacacaattcatacatgggtataaacactaatggcagattcctcaactaaatgttacaattgtgtgttggtacgatacacacacagacacacaaagtgcagtgccactgcagttgagtgaac
attattttggtgtcttgtacaggcgtgcttctgaataattttccatgttaaggttagactattagctttgcacttcattcatggcagatgatcagtaattattgatgtttatagaactaaaaatgtaactttttttgttttgttttacagatgacataaagtttcaatactgttaacactgcacaacttatgaaaaatc
acaattagatatcaatactatgatgaaagaccaacagtaaaattactttattgtgatatacaatagactccacatgaactctagaaattaaactttatgtacagagtaagtggctaacacttaacattgagctctgtctaactacattgtaattatttaattaatcaattaaagacacttggtatgcaataaac
atgtttctatttttaaggaaagataagaaaacacagaagatgctgtataaaatacaaacataatcaacaacttaacaaaggtatgatataaatttatttttctttgacattctcactatttttttatttgtattggttgtcctgctttttatttatactggcatcttcacagacattgaaacaaaaccctgtcgctttctc
aaatttgtcacattgcgtggtattcagacaacaaaactacatacaattacattaatttggatgtttgtccaagtccatttccttgaaatgtttccagttttcaaattaaaccttccagttaataattaacacaattcaaagtgacattcatgtacagcaaatataatttcagccaacaatgcactagtagtttaa
gagtaagtgatttcaacatgttcaatgcaattttagccaccccctgcaacacacacaaacaaaagcattatccttttacagtggtacgtttatctccttttatattgaattaatgtctgcactcattcactattatgtcacagattagcactgtaccctttttatttttcaggtgtatacaccctgaggaattcttca
aacaaaaacgcaggtaagtagaaaaggaaaattcatccaaagtggcagatctgcctatttgtcatagaaaaaacactggcagatctagcctactgtgtgtacgtgtgtgtgtgtgtgtgtgtgtgtgtgtatatatatatatatatatatatatatatatatatatatatatatatatatataaa
atgacaggcagttacaaattaagagcagtgccacaggttttatgactggaaaatctggaccagtttcaggtctctacatttttaaagtaggtgcttttgcacccataatgccacatgcacacaaagctatattgtgatatatttcttaatttaattaattaatataaatggctcaggaatagtttatcagattg
atgactgctgccgtcacgcaactccctttttgtaagccagccaagagctcaataataccaaaccaacaagatgtaggtgaaaataaacagcatgcttacagcaatctccacaatatgtagtactgtagaatgtaatgtattagtctaaaccaaatactgaaatgtttgtttgcttttcccaggcttttgaa
tgtattcacatttcttgcattcattgttaaatacaaaagcatgtgaagtaaaacacactctggattttccccatgatggaaattttatattctacatataaagaaaagtgtatccagtataacttattatatgagtgccagtttaagtgcacaccctttttcttttaactggcattgttcctaagaataattgagtac
accattaaaggaacattaatatacagtattgtatattagaaatttactgagcatatataaatatatctgtcctctctctctctctctctctctgtcgctctctctctctcatgtgattaacgctggatggtcttctagatagtttattagattcctgtagtaagttaattagcatatgtctgaaaatactagttcatg
atactaaacagctgatgccactaaagacaaactcaaaacatacttgtttcctttccacagaggttgacaaatccaagaatttgaaacagcagcgttgcaggatgtgtcgagtccagcggccggacttaccgctccctcactaatggctaaaataaagcacactttatatcttttaacttgaaaatcta
ctattttgcaaatagatacactttttctttttataatggaataaccaaaagcacctagctagtttgtgtgttttgcatgcagcctaactggctaaaaacatgcaatataatatttttatagcatataatgagcgcatatcacatgcattgccataaatcatatacattaaacattatacagcttaatttgctttca
ttttgtgtgtttaaaaagtaaaaatctaagttgagatggaatcttggaaagggaacactataattaaaatgtaatatatccttgtatcctgaggtgatgttttattttctcagtacaagaactacagccgagaaatgctttcaaaataaagaatgggcattcacctaaccaaggaaaagagctcagaaaca
accaggttcatacaaagtaccaaatcattgaaagaaaaaaaaaaaaaaaacaacctgtgcttacaaagtggtatttacattcagtggtagaaacaggattcattctgtcctaagaatacacactggaaatcattttattatgttaccgtttcaattataaaacctaataaactatatcaaatagtacattgt
ttccatcaacgacttgtctcctgtattttctcttaatgatttgttttcgtgatgaaatgctgcaatactaataggaaccactagcataaaaaaaaaaaaatgctttttaaaagaaatatatcctactactcataatatatgtatcttttttttttttttttttttttttttgtcttgtaattaacttcttgctcttcattactgct
ctaatccaattaaagatatgacgtactactctgaaactgct

### (C) cDNA sequence (coding region highlighted in yellow)

tgtaagctggcaatctgcactgtatgtattgtgagtcatcagtgaagggtacaagagggttgggtcttaagaaacctttatatagcttaagtaaaaggtggcacacaggttctatttttcagtatcaaacataccagttgcgatatgaggaagttgactattgtgggcttggtgcttgtgttaatgctttgt

ttctctgtactttccactgaaggaaagataagaaaacacagaagatgctgtataaaatacaaacataatcaacaacttaacaaaggtgtatacaccctgaggaattcttcaaacaaaaacgcagaggttgacaaatccaagaatttgaaacagcagcgttgcaggatgtgtcgagtccagcggc
cggacttaccgctccctcactaatggctaaaataaagcacactttatatcttttaacttgaaaatctactattttgcaaatagatacactttttctttttttataatggaataaccaaaagcacctagctagtttgtgtgttttgcatgcagcctaactggctaaaaacatgcaatataatattttttatagcatata
atgagcgcatatcacatgcattgccataaatcatatacattaaacattatacagcttaatttgctttcattttgtgtgtttaaaaagtaaaaatctaagttgagatggaatcttggaaagggaacactataattaaaatgtaatatatccttgtatcctgag

**(D) Amino acid sequence (signal peptide shaded)**

MRKLTIVGLVLVLMLCFSVLSTEGKIRKHRRCCIKYKHNQQLNKGVYTLRNSSNKNAEVDKSKNLKQQRCRMCRVQRPDLPLPH

**Fig. S12.** Information about possible GPR15LG (isoform a) from *Polyodon spathula* (Mississippi paddlefish). The information is downloaded from the NCBI gene database (https://www.ncbi.nlm.nih.gov/gene/121322081).

#### (A) Gene synteny and genomic architecture

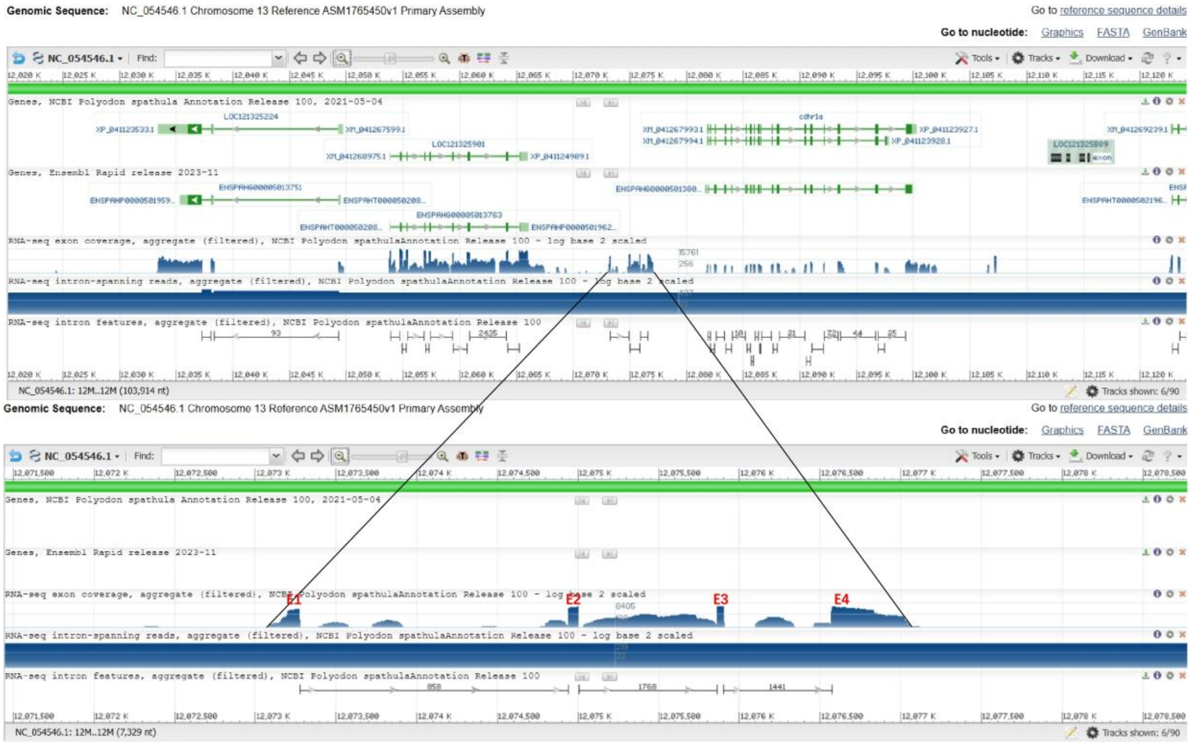


#### (B) Genomic DNA sequence (exons highlighted in yellow, coding region underlined)

atctgccctgtatgtattgtgagtcatcttgtgaagggtagaaggtacaagagggttgggtcttaagaaaactttatacagcataaataaaaggtggcacacaagttctatttttcagtaccctaaacagtccagcagaaatgaggaagttgacaattgtgggcttggtgcttgtgttaatgctttgttt
gtccatactttccactgaaggtaagaatgttacaattgctttaacttttaaatttttaaaattttaattaagatataacctacagtgttatgataaaagcaaaaatactgaggaactagtaattcatatacaggcaatattttttatatgagtgacaagcctttaataactatacatttatcttgacaaataataga
caattttccctgtatgttgtatgttgaacaataggcctgtttaaacagaaaaggctgtttttttcactggactttttaaaaattattattattgtaacctgtatgcggtgtgcaatgccattttctacaacactgtctagttctcatagtaaagtgctgcatgttacagtgtatattggaactaaaaggattgtatt
gtgataaccacattacagtaatgcattcgtatacaagcacacattctttttagacatgtttttgtatgggtgtttcacttttcaatgagagatgagaggtttgttggatttatttgaaaataaatgtatgtttcagaacaaagtaccacagctgcacagtgttatatcactatatagtgtgtactataaatattttc
tttgctggttttgtacaaattcagcctccgagttacatatagcctacctgtgctggacgacaaaaaactggtggcattattttgttactttcatagttaagtgacaaagctaaatctttcaaccatatttttattataaaaaataaaaataatattctcatcaacttttgtttgtgcaatgtatagtgcaatttaaaa
acataagtatgttttctatctgttttctacctaagcagaatatgtattctgacctgtatcacttagcaatatatcatttatttcaagctgaaagacactagcagatgaattcaatattaattaagttaagatacccgacatgtcttcatgtacatttgttattaacagttatatattatgattagaaacaataacact
aaattacagtagttttttttttttttttttttttttgctataacttatactaagatacaatcaacacacaggtataaccactaataacagatccctaaactaaagtgttacaattgtgggtttgtactacacacagaaaacaatgtgtgcagtgccactgtagttgaataaatattgtgttgttgtcttgttacagg
aatgctcctgaatagggtctaattagggactatttgctttgcacttcagtcaatttattgatgttaactgaaaatgaaactttttttgttcaaatactgttcacaaatgatcatgaaaaatcatagttaggtagcaatactagtgatgaccaacagtaaaattattgtattatgtcatatacaatactgtacaatta
gcttatgctagcatacaaatagactcaaaattaactgaataaattcaactttatctatagagtaagtgggtaactctttacattgagctgtgtctaactacattgtaattacacggtaaatgtatatgtaattacacagtaacagaatagtaactaacaaaagataccaactacctgaatattactttttgtag
ttaaacatattaaatatacaagtacaaccatttattgaattagtgtgttactacagtgtaagtgagaggcacttagtgtgcaataaacataatgtttctctttttaaggaaagaaaagaaaacacaggggatgctgtataaaacacaaacatcatcaacaacttaacaaaggtatgtcataaatgactttttt
ttctttgacattctcactattttttttattttatttttttattattgttatagttattggttgccctgctttttatttatactggcttcttcccagacattcaaattaaaagcggtccctttcttctgcctaatgtccatttccctgaaatgtttccattttgcaaataaaacctttcagctaatgattaatgcaattccaagtgaca
ttcgctgactcatgtacagtaaatatcacttcagcccaaaatgtacaaatagattaagagtaagtggtttcagataacatatccttatctgtgattcagtttttattgcaattcctcaatcgaattcagaaaaaatatacctgatgtatgacctacataacatgggttggtgttatagtctgggcccgctgct
ctgtgtctaaccctaaagcactactatccagctttcacttcattactgtccctttacactcccccaaatactggcgagactcctttatttataccgctaagccagaccaccatttaaatatatttttccaataattcagtttttatgatttcaccttttttttcattttacatttcctctaaaaaacacatatcaaattac
acatagcaaggaaacaaacattccaaaaaaaataaatgaacccctttctataggctagctagagtcacattgccttgtgttacccacaaggcaatatggccaccccccggcaacacacaaacaaaaacatttttcttttacactggtcattttatgtccttttatattgaattacttcattaatgtctgcactt
gttcactattatgtcacaaattagcactgtacccttttatttcagtttcagactccatgaggaattccttaaacaaaaacccaggtaagtaacccagtggcagaatggcctagaatggcctattagttatacggatttagcctactgcatgttagcctactgcatgtttttatatatatatatatatatatatat
atatatatatatatatatatatatatatatatatatatatatatatattacatatataaggcaataccataaagagcagtgccacagattttaggactggaaacttttgaccagtttccacgtttttaaagtagatgcttttgcacatagaatggcacatgcatacaaagctatatttgtttaatctttcttcaattgc
ttaactgatttaaatggctcatgataagcaaaacaacaacaacgtatctccctttgcaaaccagccaaaagtttaacaatgccaaaccagtaagatttaggtgaaaataaactgcacgagtacagcaatctcccactagctatagaactgtagaatgtaatatattagttaaccaaatgaactggcatt
gttccaagaataattgaatactgtctcttatgtgatttagatgaatggtggatagtctatatcgtttattagattcctgtagaatgttaagcatatgaataaagttgctgatgataccaaacagctgatgatactaaagacaaactcagaaacatgtttgtttcccttccacagaggtcagcaaatacaaga
aacagcagcgttgcagggtgtgtccagtccggccggacttacctctccctcactaatggctaaaataaagcacgctttatatataatttatcttgaaaatcaactcttttgcaaataaatgcactttttttacatataataaccaaatgcacctagctattttatgtgttcttgcatgcagcataacagattaa
ggtaaaaatatataatatgctaatatttttaatagcatataatgagcgcaaaacaagattccttctgtcataagaatacatactgtactggaaatcattttatgatgttactgtttcgattacaaaacctaataaactatataaaaatagtacattgttttcatcaattcctgtatttcttcataatgatttgttttattt
acttgtttttttttctgatgaaatgcagcaataatattagggaccataaaagagcatttaataaaacatactactcatagtagatgtatctcgttgattttttttgttttatatgaatgtcttgctctttattactgctctaatccaattaaagatatgatgtactacttggaaactgtatttttttaaaaaaattataatca
atatgcac

#### (C) cDNA sequence (coding region highlighted in yellow)

agggttgggtcttaagaaaactttatacagcataaataaaaggtggcacacaagttctatttttcagtaccctaaacagtccagcagaaatgaggaagttgacaattgtgggcttggtgcttgtgttaatgctttgtttgtccatactttccactgaaggaaagaaaagaaaacacaggggatgctgt
ataaaacacaaacatcatcaacaacttaacaaagtttcagactccatgaggaattccttaaacaaaaacccagaggtcagcaaatacaagaaacagcagcgttgcagggtgtgtccagtccggccggacttacctctccctcactaatggctaaaataaagcacgctttatatataatttatcttga
aaatcaactcttttgcaaataaatgcactttttttacatataataaccaaatgcacctagctattttatgtgttcttgcatgcagcataacagattaaggtaaaaatatataatatgctaatatttttaatagcatataatgagcgcaaaacaag

#### (D) Amino acid sequence (signal peptide shaded)

MRKLTIVGLVLVLMLCLSILSTEGKKRKHRGCCIKHKHHQQLNKVSDSMRNSLNKNPEVSKYKKQQRCRVCPVRPDLPLPH

**Fig. S13.** Information about possible GPR15LG (isoform b) from *Polyodon spathula*. (Mississippi paddlefish). The information is downloaded from the NCBI gene database (https://www.ncbi.nlm.nih.gov/gene/121325901).

### (A) Gene synteny and genomic architecture

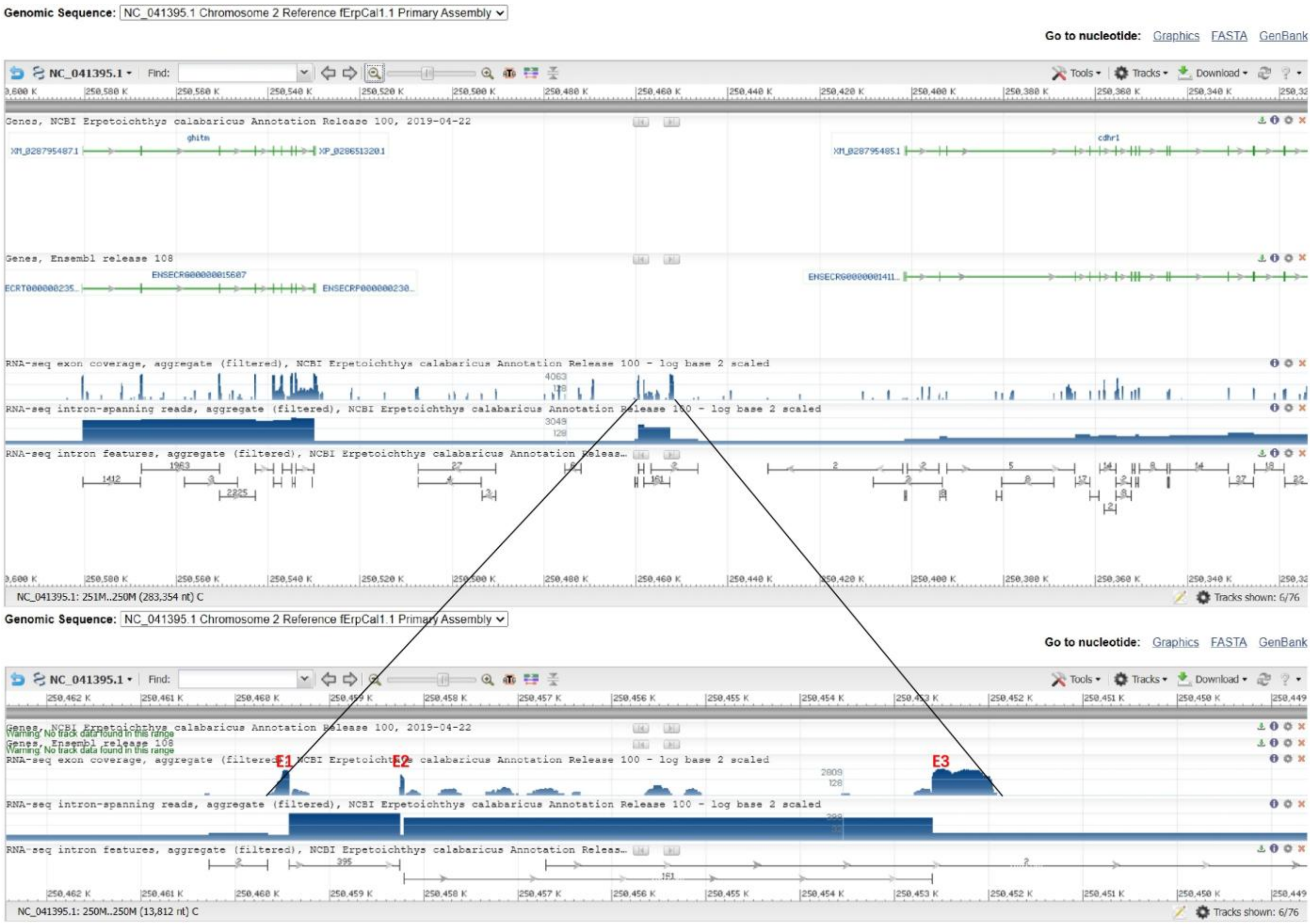


### (B) Genomic DNA sequence (exons highlighted in yellow, coding region underlined)

ttttaccatgtaacatataattggataaaagcccaatcacataatgtatatactgtaatataaaaaaataacaagcttgagcctttcagatcatgagagaaacagctagataaaatcattaccttaaagaatatttcagatctttgctttgtactacctggaaatcacctgttgctgttcttattgttttctgtgt
gtttctacagtacatactgggtttaagtttcagagggctggccttaaaattgaaaaggtgaaccattgttgattttctccaatgcccccatgcacacttaagaacaaataaaagacagctcattcttcattccttcggcagacttacagcaagcattcagaatgaaggcacacgttttcctggtgtcggt
gttggcactgatgatcactatttctgaattttgctgtgaaggtaagaaaactgacaaagaaggaagttgattgattcataaagcattggattaaactttgtgtaattatggaagttgaggcaggctttttatagcatgtagtgaattgttaccttttattatacccaatagatctataccattcatgaatttatat
acagtatatactaaaggacaggctcattagtgggaaaattaatccaagaatgaacacagcaacttaaagtgaaatcatgaaaatcatgtttttgaaaataaggcaaagtgtatattgagttctattagaaaaaaacaatttcctaaaactggataatttaataaaggaaccattatgttcaacatgggc
aatgatgcattataaaaagaaacactttaaagattgttaattatttactgagctaaagtgaaagtattgaaatctgagagttcttataatacagtaaggcatttcacattaattttacaggaaatctgcataaccttaacattctgttgtgtttaatttcaagacattctgttatattatatgtgtaaccctgataaa
caattttaaaatactctgtaaatcaaagcggtatctttaagttaaagccgttaatgttaattcatactgctctgaatgtaatatcccaatcaagttatctctttaaatctaatgcattgaaagttttgttacttttaactcaaattaatcaatgtgtcagcactcattgtatgcattttatagataatgatcacacgtta
aaatataaatatgtgtgtgtagcttgtataggctgaccagacgtcctgattttgaccacaacagtcccagtttcaggctatgcgtcccattttggctgaccagacgtcccgtttttccgagacagtcctatttttacgtatcacgtcccggagtttaagaaactgttccgtttgtcccggctttaggtctcc
gaaactagaagcagggagacttcagtttcatgtgcacttttacaaggcgcgcgatactccatccgagtctgtcattttatttgcgcccggcacactcgctgcattaatgagttgtgtccgtctgctcagcctgtagacaagtgtcccggtttgggattttaaaaacctggtcaccctactcttaaacga
attgcaactgatatttacaattgtactgctattctaattattctttttcatttaaggaaggagaatcaaatgctgtcacagacttaagaatggtatacaaggtaacccataaaacattttgtaattattttactcactaagttcaacataagaatttttatgggctttaattgtattccaaaaaataacagatggctt
gtcttcaaatgagggagactgaagctatgattcatgataattagctagatctgcaccacaattgattagcttccttgatttggactgcggtgggttggcgccctgctcgggattggttcctaccttgcgccctgtgttggctgggattggctccagcagacccccgtgaccctgtgttcggattcagc
gggttggaaaatggatggatggatgtgatgtcttttaaaaaattaaaatgttccaggacaatgtgtttatatttcttttataaatttggctttagatacattgtcaataacagtacacagaacaacatatcttctggttttcttgcaaaaacagttttagctatacaaataaagttacaaatttggtttcagtcag
caaatttcttttttgaaaataaaattgttgtgatactaagactatgtcgctgtattctgcacaattgaaaggtttatgcatataaaatatgtgtgtgtgcagcctcatcctcacaattgcttgaccaaaaagatattagtaagtgaaaaaggttaaagatagcgcttttgaaattaatttgaaactatctgaat
cattgtggaatatatgaaaacttcttcaaacataaattatttttaaaggatcttttcatttgtcattttattacagcttggacagcataacaataatttaaaacaaagaaatttcataaaagttttttaagagtttttttgtgttaacaataacttaattcagaaacacaaaaattgatgtacatacaaatattcaaa
gtgtaatggatgtaaaagtagaaaaaatactttttttattttttctaaagaatgaaacctatactctatcacagccagagggcgctgtaaagcaagtaattgaactgcagcaaatacaagccttcaaaagtaaatctgagaatgtaaatgtaaagataaagtaaaactgtacattagaacacttttgatgg
gaacaggccatcgactctccaaaataacatcaacttcacatctgaaagtccctaaagcccatctttctaccacactaaaaggtaatttattcaaagtgtatggtactttgtgtgaaaaaaactctcaaatatttctgtgaaattaggccttaaaattccagctgtttacaactatttttgttgaagaatttattt
taaagttacagtggggatccactataataattctaatataattttaatacttcagtcatgctgcctcttaatctcttaacagtaaataaatgcatagactttaaagaaaaaagttgctggttcatgatgcatgactccgagaaggtgtattaacctacctgttattcacgtgaggaatacagtatatgcaaaa
caaatgtagagttctgcaattataaagtgcaatgtaaaataaattgtcaccttaaagtgcaataaaaaattgtgacattgaaagacattcttatttgctccaggggtgttgtacaatggttgaccaagcacttgggcattaataaagtaaatcaaagcaaaaaaaaattttttaaccccacgtaattcaat
ctcacctgagtcttgtttaggcattttagcctaggtttataaatctctccatgactgtttattgaaatggatgatgtacataataaatatgtacatgttccatgcttttgatattgatttgtatttgtgtatttgtaaatttaaaaattcatatagaaacatgcatatcaatagtcaaacatgtgaaagaaagttgtttg
agcataacaacatgattgatagcagttgacaacaaaagagagaaaaattaaagctcctgataacagcatttaacaataaaatatgaagctatatatattgtggtagaaatttaacattttattaaagaaacaaatgtctcctaacaggacaaatcagtaatcagacaggagaaaagctggtcattata
tcttttaattactcctcggtaaaatgtcttagagggagcattcttcaaaagcagtccgttacagcatagtcaagaatagaggttcatttttaaactgttgaatttacattcagtatcaattaatgcatacatttattctggttggttcattggtttacacataaataatttatataaaataaaagtctattttattata
gtctggtttttcttatacctttgagttgagctaccacccttgaaatttctgtgcatataacaactctccattcttatttatatgacatctgctgatttcttagtcatctcttggtcatccttcagtacattttgtatgttacatcaaccttcttctagtacatttacttgaccatctcagtatttttcctaaaccagttctta
tatttcagtttacattttgttgttcacttgacctttatttggtttcctgttgtgcctgaacagttttctgacatttattaaccctttatgctttttctttaagatgaatgctatgtcaccctaaaatcattcttccacaatgtaaatataaaaacaaaagactaaacaaatcagtctaatgctatcgccatattattcaaa
atatcatcctcaataaaaggataactgtctgcatgtttgtctgtgtgtccgtctggttgctaggtctctggcattccaaaagatggcacatcacaaacatttttagtaataaagggtctttcaattaatatctgagaaaaggaaggcagccatccatagaatacacccttgcagcatatgcacaaaacttt
caatcattcaacttaaaatctttcatcgatcacatttatcttgtttaatattgtccaaaatgttttcagagcaagagccaacctgtgaacattgcaatcgaactccagcctcactgggccacatattttgggtgtgcaccaaattaacaacattttggacaaaaatcagcctccgtgtcacaatcaccccta
acccattaagagctgtgtttggtgaactcccagatgagcttaaagtgaagaaggttaaatttattgtaattgattgcttattccacacttatagacgtagacttattttgctcaactggaagaaacccagccccccttttataagtcagtgggtaactgatgtcctatactatttgaaattggaaaaaatcta
attctcgcttagaggatctcgtcaaaagtttttaaaacacagcaggagctaattaataaaatgttacaataagcatttatattgggaaaaggacattttccctttttttttctttctcctatttttgcctatgctgttggttctcctctctcttaggtggggttgaagtctgtctgattaagctttgtctctttttgttcct
gctttcttttttttttcttatttgactaagaagtcatttgttgtataagtttgacttgattgtttgcaatgttattttcttcaaataacataaaaaaatgaattgcatttgttattttaatagatggtgcatcacaaacattaacacaatgtttatgaattccataccaaatggcatataacagagacatatgtggagatc
aacccatcttaggccgggtttatacttcacgtgatgttacacatgctccagtggacactactgttatgcatagtgttgtactatttatacttgcgtgcgtactttacataaatctggaagattccatcagttggcagtgtgatatatcatcacagtgagaaaacgtttgactttattgtgttgtgaattgcctg
acacatccattaaattctgaggaaaccttgccacaatatctctgaaaaggatgtttaatgattacatctatcgatccagggatgcacccattccaccaagcaccgggctcgagacagaaacaatccctggacagggcatcagctcatcacaaggtgaacacaagcactcacatacactaacgtc
attttagcatcaccaaatccacaaaccttcatgtctttgggaggaaaccggagcatactgtggaaactcgccagggaacatgcaaactccaggcagggaacaccagcgacatgactccctgtgagacagcagctctaccactttgctaccgtgccgtgcccacgtgtgtaattattaacagtat
tcattatttgaataatgttaacgatttatctgtaaaatgtaacatacatactttaatgcatttcatcatgaaagtgatatcaagtatagtggaacctcggtttgcgagtaacttggtttatgagtgttttgcaggacgagcaaaaatttttaataaattttgacttgataaacgagcgaggttttgcagcacgag

tagtatgtatactctttgtctgctgagcgtcatgtgatcacaactgagctgatggttcttctctctctcgctgcgggattgtgggcaatcgtctcctattctctgtctgagtcggcatgcctcactcatagtaaacatccgtacgagcatatactgtttactacagcattatagcattgtgactgtgtgtgcg
cgcatgcttgtgtgtgtatgtgtgtgcgtgtgctcgtacgtgcgtgtgtgctgtgacgtgcgagtcccagtcttgcacactaaaacacaaagcttgagtctcagtactttagctacatcagctttattttacttgaaatagacacagcagtcaggcagggccctgcacatttataatgttccttgttcctt
gtagcacccatcgacggcaggctcttatagcatgtccgcgatcttttcggattcacttttacggtgaactgctacagccctgggagactgcgattgctttgggacgctcttccgcgtgtcgtcccgttgggtggaatcccacaagagtttagaaactcactcacaccagccatgattcttttcaaag
gtaaagtgcaggttaatttgttttatgtatttttactttatattttgcattaatcatttttatatgaatagttttgggttgtggaacaaatcatctgagtttccattatttcttattgggaaattcactttgatatacgagtgctttggactacgagcacgtttccggaacgaattatgctcgcaaatcgaggttccact
gtataaatctaaagattctaaatgtgcagagagtatcataagtgtaatgtgttctgtgtggtgatctattgctgcttgttgctgctgtcaggtcaggaggaagccctagaagcacatagtgattaacaactgggttttaagatgatgtttacgatgatctactttcatgataaaataaactacaaggttaaa
gtggacattttgagagccgaaatttccactttaatcacaaaatagaccttgggaaagttgtctctgtaagcaggagtgctttagatatcttttactgtggggtaactcttttgtttgcaagttacgcaccaggaaagactgctcaaataatagttaaaagctcaaaacccttctcaaaacatttgtttaaat
acatcaagtgatggaaagaaactgtaaacaaagtgactaaaccctaataacaactgatcaattctctttattcccacagtgaatggcccaaagttccggaacaagcgccactgctggatctgctcaaatatgaagggggcaccggctaaagcaaatggccattctgatatacccataccaactttt
taagtaccttgtagctccagcgctgtaccgccatcaaacagaaatgacataaactgatctgtacaatcctttcatttttctaaccgctgcaacaagcatgtcattcatttataaggaaactgctttagttattttctttgtattcttttatcaatttatatatttgagaaagttacaaagtagtccgcaaatttgctt
aatcagaaaagtgtataaaatgtctaaacattaggagagaagacagactataaagcatgaaggacgctatctcagatgaggattcaatagagtctttgttttagtaactgtaattgaaattaattgtaaagttctagggatatggagcttggaccaatgctaacgagtgtgtaaattggacacatgga
tggatgaatgagtggatttttatggatttttgtgagggtgcatttccatactgcatacaaaacatacacaaaaatgtatatataagactttgcttgtagaaatgtttaaacgtctgcagttttaataaaaaatgtatgcaaataagtgtttgactttttataaatctctgttcatatgtattttatactccctctctctt
tgtctttccatttctgtctctatccattttttccagatagatatgtctccatcccctcattataatgtataaagctctgcagttttcatgtgcttaatgtgtttaatac

**(C) cDNA sequence (coding region highlighted in yellow)**

gtgaaccattgttgattttctccaatgcccccatgcacacttaagaacaaataaaagacagctcattcttcattccttcggcagacttacagcaagcattcagaatgaaggcacacgttttcctggtgtcggtgttggcactgatgatcactatttctgaattttgctgtgaaggaaggagaatcaaatg
ctgtcacagacttaagaatggtatacaagtgaatggcccaaagttccggaacaagcgccactgctggatctgctcaaatatgaagggggcaccggctaaagcaaatggccattctgatatacccataccaactttttaagtaccttgtagctccagcgctgtaccgccatcaaacagaaatgac
ataaactgatctgtacaatcctttcatttttctaaccgctgcaacaagcatgtcattcatttataaggaaactgctttagttattttctttgtattcttttatcaatttatatatttgagaaag

**(D) Amino acid sequence (signal peptide shaded)**

MKAHVFLVSVLALMITISEFCCEGRRIKCCHRLKNGIQVNGPKFRNKRHCWICSNMKGAPAKANGHSDIPIPTF

**Fig. S14.** Information about possible GPR15LG from *Erpetoichthys calabaricus* (reedfish). The information is downloaded from the NCBI gene database (https://www.ncbi.nlm.nih.gov/gene/?term=growth+hormone-inducible+transmembrane+protein%2C+Erpetoichthys+calabaricus).

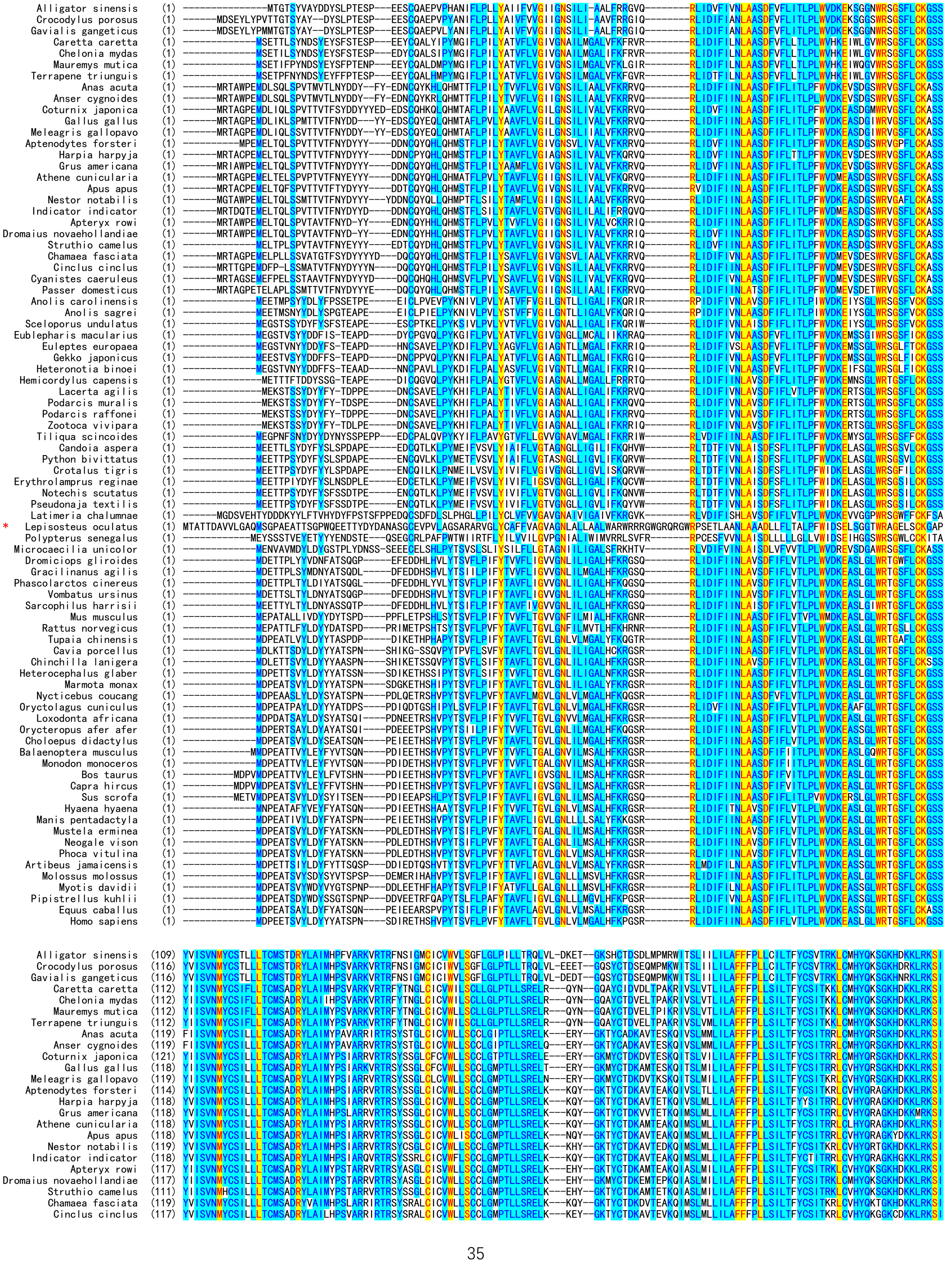
Alligator sinensis
Crocodylus porosus
Gavialis gangeticus
Caretta caretta
Chelonia mydas
Mauremys mutica
Terrapene triunguis
Anas acuta
Anser cygnoides
Coturnix japonica
Gallus gallus
Meleagris gallopavo
Aptenodytes forsteri
Harpia harpyja
Grus americana
Athene cunicularia
Apus apus
Nestor notabilis
Indicator indicator
Apteryx rowi
Dromaius novaehollandiae
Struthio camelus
Chamaea fasciata
Cinclus cinclus
Cyanistes caeruleus
Passer domesticus
Anolis carolinensis
Anolis sagrei
Sceloporus undulatus
Eublepharis macularius
Euleptes europaea
Gekko japonicus
Heteronotia binoei
Hemicordylus capensis
Lacerta agilis
Podarcis muralis
Podarcis raffonei
Zootoca vivipara
Tiliqua scincoides
Candoia aspera
Python bivittatus
Crotalus tigris
Erythrolamprus reginae
Notechis scutatus
Pseudonaja textilis
Latimeria chalumnae
* Lepisosteus oculatus
Polypterus senegalus
Microcaecilia unicolor
Dromiciops gliroides
Gracilinanus agilis
Phascolarctos cinereus
Vombatus ursinus
Sarcophilus harrisii
Mus musculus
Rattus norvegicus
Tupaia chinensis
Cavia porcellus
Chinchilla lanigera
Heterocephalus glaber
Marmota monax
Nycticebus coucang
Oryctolagus cuniculus
Loxodonta africana
Orycteropus afer afer
Choloepus didactylus
Balaenoptera musculus
Monodon monoceros
Bos taurus
Capra hircus
Sus scrofa
Hyaena hyaena
Manis pentadactyla
Mustela erminea
Neogale vison
Phoca vitulina
Artibeus jamaicensis
Molossus molossus
Myotis davidii
Pipistrellus kuhlii
Equus caballus
Homo sapiens

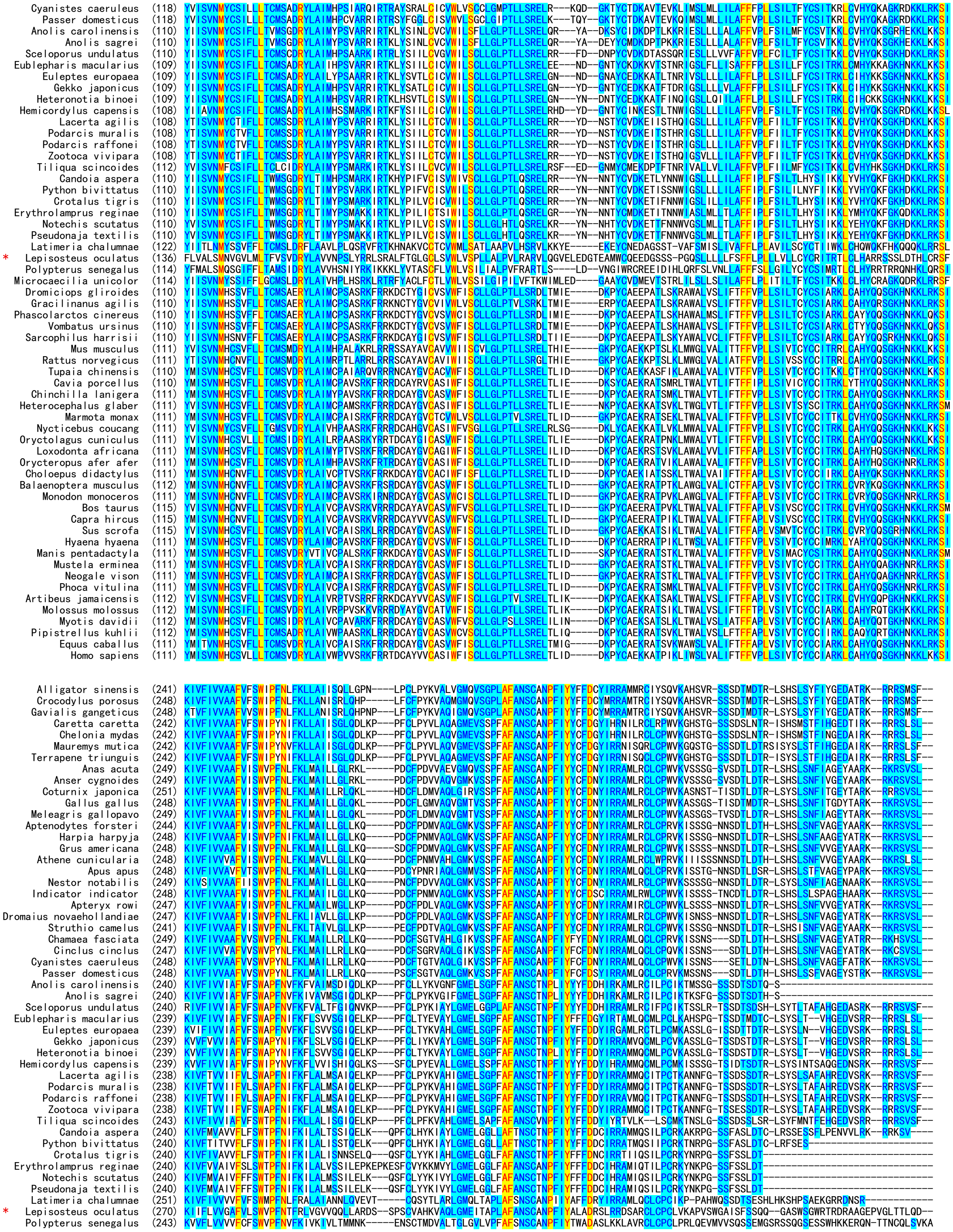
Cyanistes caeruleus (118)
Passer domesticus (118)
Anolis carolinensis (110)
Anolis sagrei (110)
Sceloporus undulatus (110)
Eublepharis macularius (109)
Euleptes europaea (109)
Gekko japonicus (109)
Heteronotia binoei (109)
Hemicordylus capensis (108)
Lacerta agilis (108)
Podarcis muralis (108)
Podarcis raffonei (108)
Zootoca vivipara (108)
Tiliqua scincoides (112)
Candoia aspera (110)
Python bivittatus (110)
Crotalus tigris (110)
Erythrolamprus reginae (110)
Notechis scutatus (110)
Pseudonaja textilis (110)
Latimeria chalumnae (122)
* Lepisosteus oculatus (136)
Polypterus senegalus (114)
Microcaecilia unicolor (114)
Dromiciops gliroides (110)
Gracilinanus agilis (110)
Phascolarctos cinereus (110)
Vombatus ursinus (110)
Sarcophilus harrisii (110)
Mus musculus (111)
Rattus norvegicus (111)
Tupaia chinensis (110)
Cavia porcellus (110)
Chinchilla lanigera (111)
Heterocephalus glaber (111)
Marmota monax (111)
Nycticebus coucang (111)
Oryctolagus cuniculus (111)
Loxodonta africana (111)
Orycteropus afer afer (111)
Choloepus didactylus (111)
Balaenoptera musculus (112)
Monodon monoceros (111)
Bos taurus (115)
Capra hircus (115)
Sus scrofa (115)
Hyaena hyaena (111)
Manis pentadactyla (111)
Mustela erminea (111)
Neogale vison (111)
Phoca vitulina (111)
Artibeus jamaicensis (112)
Molossus molossus (112)
Myotis davidii (112)
Pipistrellus kuhlii (112)
Equus caballus (111)
Homo sapiens (111)
Alligator sinensis (241)
Crocodylus porosus (248)
Gavialis gangeticus (248)
Caretta caretta (242)
Chelonia mydas (242)
Mauremys mutica (242)
Terrapene triunguis (242)
Anas acuta (249)
Anser cygnoides (249)
Coturnix japonica (251)
Gallus gallus (248)
Meleagris gallopavo (249)
Aptenodytes forsteri (244)
Harpia harpyja (248)
Grus americana (248)
Athene cunicularia (248)
Apus apus (248)
Nestor notabilis (249)
Indicator indicator (248)
Apteryx rowi (247)
Dromaius novaehollandiae (247)
Struthio camelus (241)
Chamaea fasciata (249)
Cinclus cinclus (247)
Cyanistes caeruleus (248)
Passer domesticus (248)
Anolis carolinensis (240)
Anolis sagrei (240)
Sceloporus undulatus (240)
Eublepharis macularius (239)
Euleptes europaea (239)
Gekko japonicus (239)
Heteronotia binoei (239)
Hemicordylus capensis (239)
Lacerta agilis (238)
Podarcis muralis (238)
Podarcis raffonei (238)
Zootoca vivipara (238)
Tiliqua scincoides (243)
Candoia aspera (240)
Python bivittatus (240)
Crotalus tigris (240)
Erythrolamprus reginae (240)
Notechis scutatus (240)
Pseudonaja textilis (240)
Latimeria chalumnae (251)
* Lepisosteus oculatus (270)
Polypterus senegalus (243)

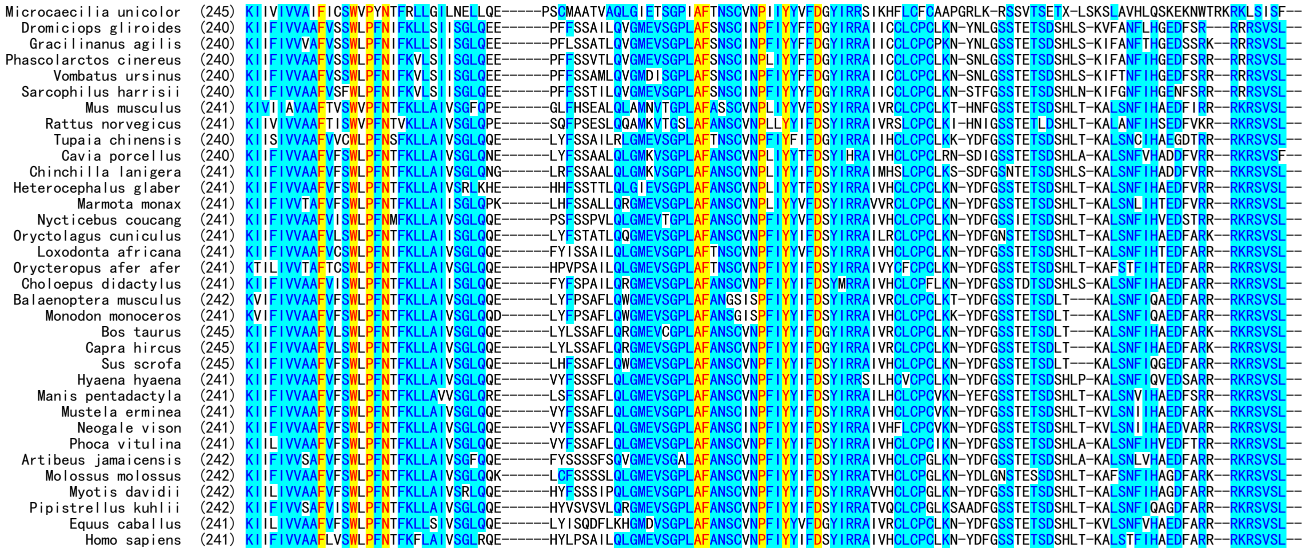


**Fig. S15.** Amino acid sequence alignment of GPR15 orthologs from different species. Their information is listed in Table S3. The spotted gar GPR15 is indicated by a red asterisk.
